# Structural basis to design multi-epitope vaccines against Novel Coronavirus 19 (COVID19) infection, the ongoing pandemic emergency: an in silico approach

**DOI:** 10.1101/2020.04.01.019299

**Authors:** Sukrit Srivastava, Sonia Verma, Mohit Kamthania, Rupinder Kaur, Ruchi Kiran Badyal, Ajay Kumar Saxena, Ho-Joon Shin, Michael Kolbe, Kailash C Pandey

## Abstract

The 2019 novel coronavirus (COVID19 / Wuhan coronavirus), officially named as Severe Acute Respiratory Syndrome Coronavirus 2 (SARS-CoV-2), is a positive-sense single-stranded RNA coronavirus. SARS-CoV-2 causes the contagious COVID19 disease also known as 2019-nCoV acute respiratory disease and has led to the ongoing 2019–20 pandemic COVID19 outbreak. The effective counter measures against SARS-CoV-2 infection require the design and development of specific and effective vaccine candidate. In the present study, we have screened and shortlisted 38 CTL, 33 HTL and 12 B cell epitopes from the eleven Protein sequences of SARS-CoV-2 by utilizing different in silico tools. The screened epitopes were further validated for their binding with their respective HLA allele binders and TAP (Transporter associated with antigen processing) molecule by molecular docking. The shortlisted screened epitopes were further utilized to design novel two multi-epitope vaccines (MEVs) composed of CTL, HTL and B cell epitopes overlaps with potential to elicit humoral as well as cellular immune response against SARS-CoV-2. To enhance the immune response for our vaccine design, truncated (residues 10-153) Onchocerca volvulus activation-associated secreted protein-1 (Ov-ASP-1) has been utilized as an adjuvant at N terminal of both the MEVs. Further molecular models for both the MEVs were prepared and validated for their stable molecular interactions with Toll-Like Receptor 3 (TLR 3). The codon-optimized cDNA of both the MEVs were further analyzed for their potential of high level of expression in a human cell line. The present study is very significant in terms of molecular designing of prospective CTL and HTL vaccine against SARS-CoV-2 infection with the potential to elicit cellular as well as humoral immune response. (SARS-CoV-2), Coronavirus, Human Transporter associated with antigen processing (TAP), Toll-Like Receptor (TLR), Epitope, Immunoinformatics, Molecular Docking, Molecular dynamics simulation, Multi-epitope Vaccine

**Graphical abstract:** The designed CTL (Cytotoxic T lymphocyte) and HTL (Helper T lymphocyte) multi-epitope vaccines (MEV) against COVID19 infection. Both the CTL and HTL MEV models show a very stable and well fit conformational complex formation tendency with the Toll like receptor 3. CTL and HTL MEVs: *ribbon*; Toll like receptor 3: *gray cartoon*; Adjuvant [truncated (residues 10-153) Onchocerca volvulus activation-associated secreted protein-1]: *orange ribbon regions*; Epitopes: *cyan ribbons regions*; 6xHis Tag: *magenta ribbon regions*.

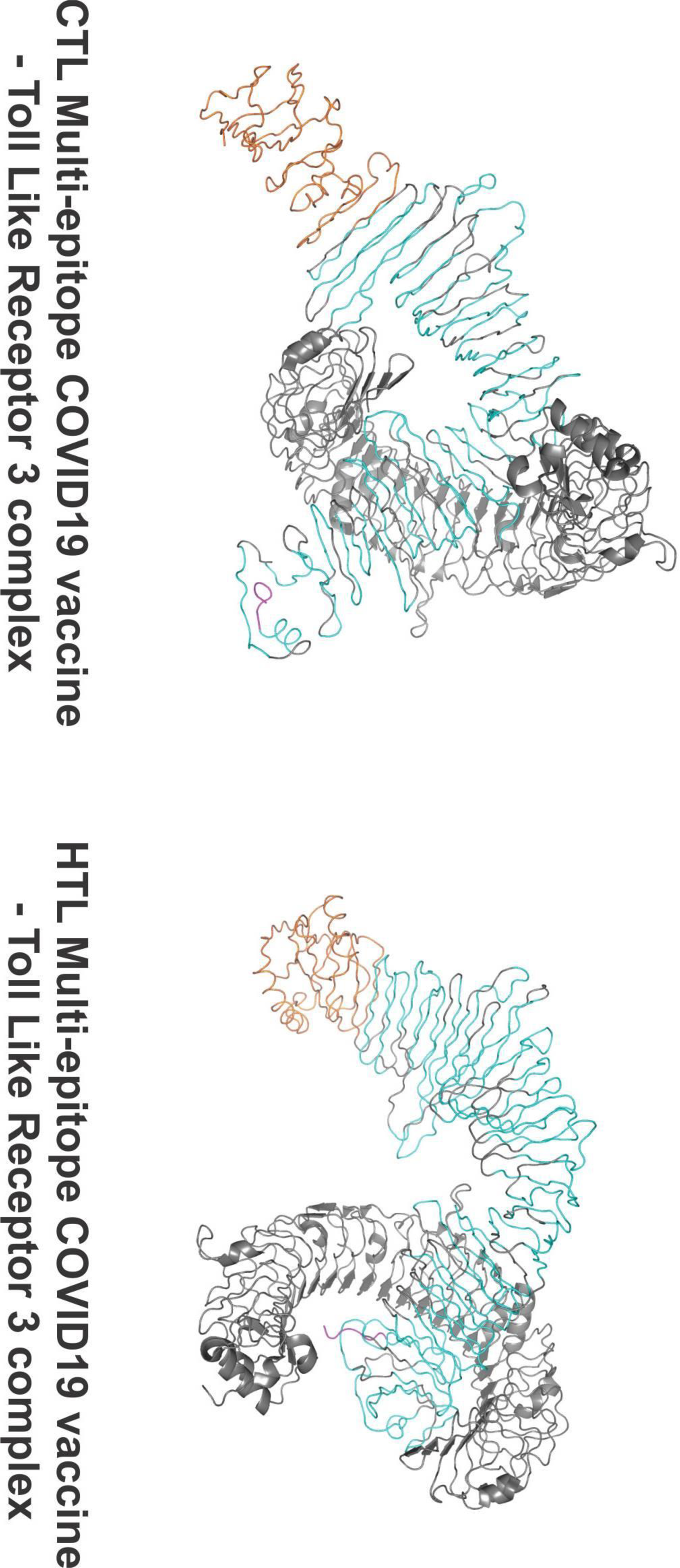

## Introduction

The novel coronavirus (COVID19), officially named as Severe Acute Respiratory Syndrome Coronavirus 2 (SARS-CoV-2) has caused the ongoing outbreak of a severe form of flu leading to death with a mortality rate of 3.4 %. The SARS-CoV-2 is a novel coronavirus associated with a respiratory disease initiated from the Wuhan of Hubei province, China. The disease is highly contagious and has spread to 182 countries/ Territories since its outbreak in China in December 2019 till 21^st^ of March 2020. Worldwide, as of 21^st^ March 2020, the total confirmed cases have been reported to be 2,66,073 and total death count reported is 11,184 (WHO Situation report 21^st^ March 2020). Overall the SARS-CoV-2 infection has put a global emergency condition. The economic impact of COVID-19 is even harsher and has put the world on economic risk. As of 9^th^ March, the downside scenario sees a $2 trillion shortfall in global income with a $220 billion hit to developing countries. The COVID-19 shock will cause a recession in several countries and depress global annual growth this year to 2.5 percent lower, the recessionary threshold for the world economy (UNCTAD report 9^th^ March 2020).

The infection mechanism and pathogenesis of SARS-CoV-2 is largely unknown yet. The proteome of SARS-CoV-2 is composed of 11 structural and non-structural proteins. These include polyprotein (ORF1ab), Surface protein (S Protein), ORF3, Envelope Protein (E Protein), Membrane Protein (M Protein), ORF6, ORF7a, ORF7b, ORF8, Nucleocapsid Protein (N Protein), and ORF10 (NCBI protein sequence database). The actual function and pathogenic or proliferative role of these SARS-CoV-2 coronavirus proteins are largely not known yet.

The SARS-CoV-2 coronavirus polyprotein (ORF1ab) with length of 7,096 amino acid (aa) is composed of 16 different expressed protein viz. leader protein (nsp1, location: 1-180 aa); nsp2 (location: 181-818); nsp3 (former nsp1, carry conserved domains - N-terminal acidic, predicted phosphoesterase, papain-like proteinase, Y-domain, transmembrane domain 1 (TM1) and adenosine diphosphate-ribose 1’’-phosphatase, location: 819-2,763); nsp4 (contains transmembrane domain 2 (TM2), location: 2,764-3,263); 3C-like proteinase (nsp5, main proteinase, mediates cleavages downstream of nsp4, location: 3,264-3,569); nsp6 (putative transmembrane domain, location: 3,570-3,859); nsp7 (location: 3,860-3,942); nsp8 (location: 3,943-4,140); nsp9 (ssRNA-binding protein, location: 4,141-4,253); nsp10 (formerly known as growth-factor-like protein, location: 4,254-4,392); nsp11 (location: 4,393-4,405); RNA-dependent RNA polymerase (nsp12, location: 4,393-5,324); Helicase (nsp13; zinc-binding domain, NTPase/helicase domain, RNA 5’-triphosphatase, location: 5,325-5,925); 3’-to-5’ exonuclease (nsp14, location: 5,926-6,452); endo RNAse (nsp15, location: 6,453-6,798); and 2’-O-ribose methyltransferase (nsp16; location: 6,799-7,096).

The SARS-CoV-2 coronavirus surface glycoprotein (S Protein) is a structural protein and acts as spike protein, its location is 21,563-25,384 aa, and length is 1273 aa); The SARS-CoV-2 coronavirus ORF3a protein has location 25,393-26,220 aa and length 275 aa. The SARS-CoV-2 coronavirus envelope protein (E Protein) (ORF4) is a structural protein and has location 26,245-26,472 aa and length 75 aa. The SARS-CoV-2 coronavirus membrane glycoprotein (M Protein) (ORF5) is a structural protein and has location 26,523-27,191aa and length 222 aa. The SARS-CoV-2 coronavirus ORF6 protein has location 27,202-27,387 aa and length 61 aa. The SARS-CoV-2 coronavirus ORF7a protein has location 27,394-27,759 aa and length 121 aa. The SARS-CoV-2 coronavirus ORF7b protein has location 27,756-27,887 aa and length 43 aa. The SARS-CoV-2 coronavirus ORF8 protein has location 27,894-28,259 aa and length 121 aa. The SARS-CoV-2 coronavirus nucleocapsid phosphoprotein (N Protein) (ORF9) is a structural protein with location 28,274-29,533 aa and has a length of 419 aa. The SARS-CoV-2 coronavirus ORF10 protein has location 29,558-29,674 aa and length 38 aa (NCBI protein sequence database).

Although the exact mechanism and roles of all the above-mentioned proteins of SARS-CoV-2 coronavirus proteome are not well known yet, but these proteins could act as potential vaccine candidates against the SARS-CoV-2 coronavirus infection. In the present study, we have screened highly potential epitopes from all the above-mentioned protein and further, we have also designed and proposed CTL (Cytotoxic T lymphocyte) and HTL (Helper T lymphocyte) multi-epitope based vaccine candidates against the SARS-CoV-2 coronavirus infection.

## Methodology

In the present study on SARS-CoV-2 coronavirus, we have screened potential epitopes and have designed and proposed two multi-epitope vaccines (MEVs) composed of screened CTL (Cytotoxic T lymphocyte) and HTL (Helper T lymphocyte) epitopes with overlapping regions of B cell epitopes. Hence the proposed MEVs are supposed to have the potential to elicit both the humoral as well as cellular immune response. To enhance immune response, truncated (residues 10-153) Onchocerca volvulus activation-associated secreted protein-1 (Ov-ASP-1) has been utilized as an adjuvant at N-terminal of both the MEVs. The truncated Ov-ASP-1 was chosen due to its potential to activate antigen-processing cells (APCs) (MacDonald et al., 2005; Guo et al., 2015; He et al., 2009). All SARS-CoV-2 coronavirus proteins mentioned in the introduction were utilized to screen the potential CTL, HTL and B cell epitopes. The screened epitopes were further studied for overlapping consensus regions amongst them. The epitopes showing overlapping regions in partial or complete were chosen for detailed further studies.

The chosen CTL and HTL epitopes were analyzed for their molecular interaction with their respective HLA allele binders. Moreover, the chosen CTL epitopes were also analyzed for their molecular interaction with TAP (Transporter associated with antigen processing) transporter cavity to observe their smooth passage from cytoplasm to endoplasmic reticulum lumen (Oldham et al., 2016; Abele et al., 2004). The tertiary model for both MEVs were generated and refined. Both the MEVs models were further utilized to screened B Cell linear and discontinuous epitopes as well as IFN-γ inducing epitopes.

The molecular signaling by multiple TLRs, is an essential component of the innate immune response against SARS-CoV-2 coronavirus. Since the rOv-ASP-1 primarily binds APCs among human PBMCs and trigger pro-inflammatory cytokine production via Toll-like receptor 3 (TLR3), hence both the CTL and HTL MEV models were further analyzed for their molecular interaction with the TLR-3 by molecular docking studies (Antoniou et al., 2003; Delneste et al., 2007; Totura et al., 2015; Farina et al., 2005). Further, the codon-optimized cDNA of both the MEVs were analyzed to have a high level of expression in mammalian cell line (human), which would facilitate *in-vivo* expression, experimentation and trials (Fig.1).

**Figure 1.**
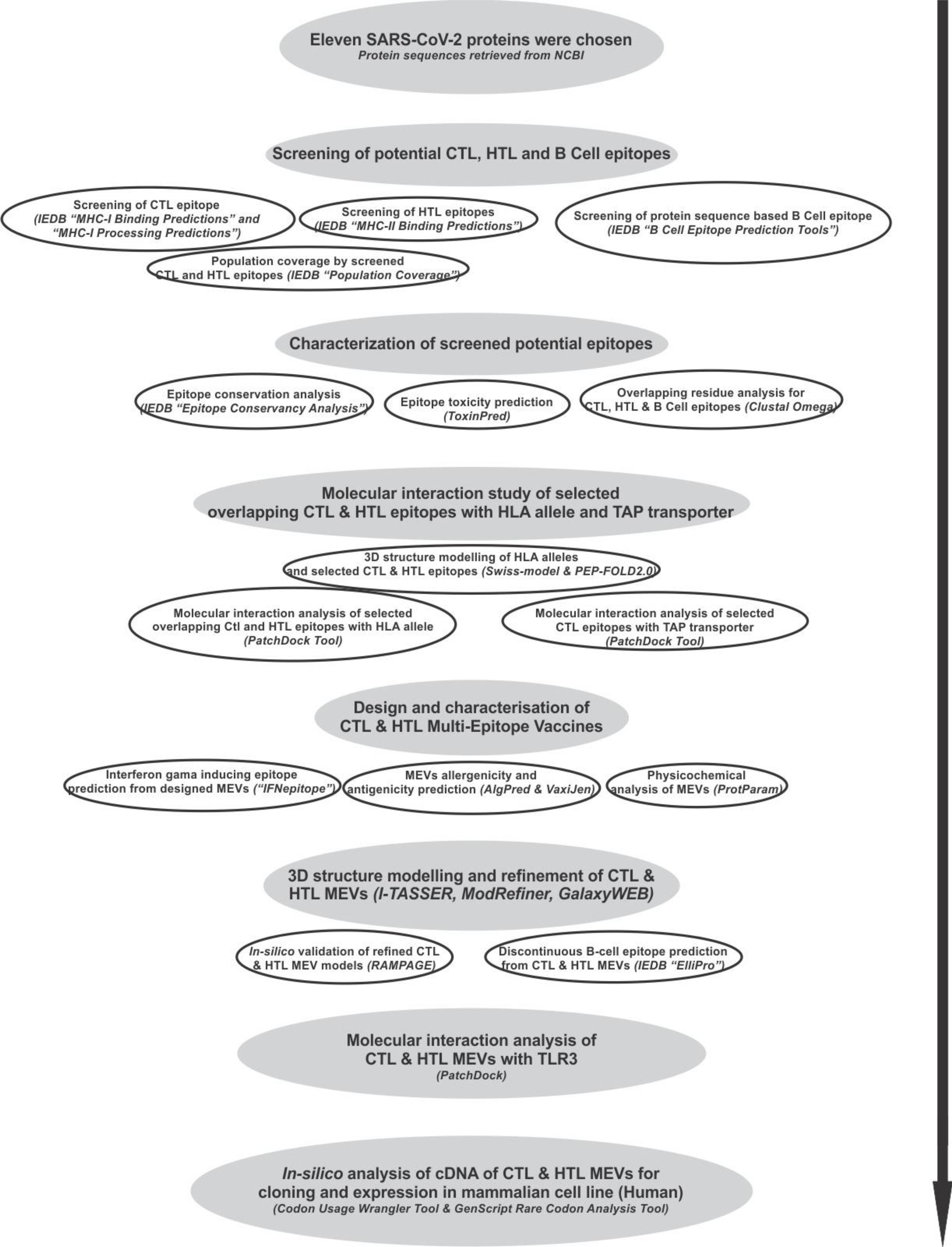
Workflow chart.

### Screening of Potential Epitopes

#### T cell Epitope Prediction

##### Screening of Cytotoxic T lymphocyte (CTL) Epitope

The screening of Cytotoxic T lymphocyte epitopes was performed by the IEDB (Immune Epitope Database) tools “MHC-I Binding Predictions” (http://tools.iedb.org/mhci/) and “MHC-I Processing Predictions” (http://tools.iedb.org/processing/) (Tenzer et al., 2005; Peters et al., 2003; Hoof et al., 2009). Both the tools use six different methods (viz. Consensus, NN-align, SMM-align, Combinatorial library, Sturniolo and NetMHCIIpan) and generate “Percentile rank” and a “total score” respectively.

The screening is based on the total amount of cleavage sites in the protein. TAP score estimates an effective log -(IC50) values (half maximal inhibitory concentration (IC50) for binding to TAP of a peptide or its N-terminal prolonged precursors. The MHC binding prediction score is -log(IC50) values for binding to MHC of a peptide (Calis et al., 2013). The IC(50) (nM) value for each epitope and MHC allele binding pairs were also obtained by this IEDB tool. Epitopes having high, intermediate, and least affinity of binding to their HLA allele binders have IC50 values < 50 nM, < 500 nM and < 5000 nM, respectively.

Immunogenicity of all the screened CTL epitopes was also obtained by using “MHC I Immunogenicity” tool of IEDB (http://tools.iedb.org/immunogenicity/) with all the parameters set to default analyzing 1st, 2nd, and C-terminus amino acids of the given screened epitope (Calis et al., 2013). The tool predicts the immunogenicity of a given peptide-MHC (pMHC) complex based on the physiochemical properties of constituting amino acid and their position within the peptide sequence.

##### Screening of Helper T lymphocyte (HTL) Epitopes

To screen out the Helper T lymphocyte epitopes from SARS-CoV-2 proteins, the IEDB tool “MHC-II Binding Predictions” (http://tools.iedb.org/mhcii/) was used. The tool generates “Percentile rank” for each potential peptide. The lower the value of percentile, higher would be the affinity. This percentile rank is generated by the combination of three different methods viz. combinatorial library, SMM_align & Sturniolo; and by comparing the score of the peptide against the scores of other random five million 15-mer peptides of SWISSPROT database (Wang et al., 2010; Sidney et al., 2008; Nielsen et al., 2007; Sturniolo et al., 1999). The rank from the consensus of all three methods was generated by the median percentile rank of the three methods.

##### Population Coverage by CTL and HTL epitopes

The “Population Coverage” tool of IEDB (http://tools.iedb.org/population/) was used to elucidate the world human population coverage by the shortlisted 38 CTL and 33 HTL epitopes derived from nine SARS-CoV-2 proteins (Bui et al., 2006). T cells recognize the complex between a specific major MHC molecule and a particular pathogen-derived epitope. The given epitope will elicit a response only in an individual that express an MHC molecule, which is capable of binding that particular epitope. This denominated MHC restriction of T cell responses and the MHC polymorphism provides the basis for population coverage study. The MHC types are expressed at dramatically different frequencies in different ethnicities. Hence a vaccine with larger population coverage could be of greater importance (Sturniolo et al., 1999). Clinical administration of multiple-epitopes involving both the CTL and the HTL epitopes are predicted here to have a greater probability of larger human population coverage worldwide.

### B Cell Epitope Prediction

#### Sequence-based B Cell epitope prediction

Protein sequence-based method “Bepipred Linear Epitope Prediction” was utilized to screen linear B cell epitopes from eleven different SARS-CoV-2 proteins. The tool “B Cell Epitope Prediction Tools” of IEDB server (http://tools.iedb.org/bcell/) was utilized. In this screening, the parameters such as hydrophilicity, flexibility, accessibility, turns, exposed surface, polarity and the antigenic propensity of the polypeptides is correlated with its location in the protein. This allows the search for continuous epitopes prediction from protein sequence. The prediction is base on the propensity scales for each of the 20 amino acids. For a window size n, the i - (n-1)/2 neighboring residues on each side of residue i are used to compute the score for the residue i. The method “Bepipred Linear Epitope Prediction” utilized here is based on the propensity scale method as well as the physiochemical properties of the given antigenic sequence to screen potential epitopes (Larsen et al., 2006).

### Characterization of potential epitopes

#### Epitope conservation analysis

The shortlisted CTL, HTL and B cell epitopes screened from eleven SARS-CoV-2 proteins were analyzed for the conservancy of their amino acid sequence by “Epitope Conservancy Analysis” tool (http://tools.iedb.org/conservancy/) of IEDB. The epitope conservancy is the number of protein sequences (retrieved from NCBI) that contain that particular epitope. The analysis was done against their entire respective source protein sequences of SARS-CoV-2 proteins retrieved from the NCBI protein database (Bui et al., 2007).

#### Epitope Toxicity prediction

The tool ToxinPred (http://crdd.osdd.net/raghava/toxinpred/multi_submit.php) was used to analyze the toxicity of shortlisted CTL, HTL and B cell epitopes. The tool allows to identify highly toxic or non-toxic short peptides. The toxicity check analysis was done by the “SVM (Swiss-Prot) based” (support vector machine) method utilizing dataset of 1805 sequences as positive, 3593 negative sequences from Swissprot as well as an alternative dataset comprises the same 1805 positive sequences and 12541 negative sequences from TrEMBLE (Gupta et al., 2013).

#### Overlapping residue analysis

The overlapping residue analysis for the shortlisted 38 CTL, 33 HTL and the 12 B cell linear epitopes was performed by the Multiple Sequence Alignment (MSA) analysis by Clustal Omega tool (https://www.ebi.ac.uk/Tools/msa/clustalo/) of EBI (European Bioinformatics Institute) (Sievers et al., 2011). The Clustal Omega multiple sequence alignment tool virtually aligns any number of protein sequences and delivers an accurate alignment.

#### Epitope selected for molecular interaction study with HLA allele and TAP transporter

Based on the overlapping residue analysis of shortlisted CTL, HTL and linear B cell epitopes few numbers of CTL and HTL epitopes were chosen for further analysis. The chosen epitopes are encircled shown in Fig.2. These epitopes were chosen based on partial or full overlapping sequence region amongst all three types of epitopes (CTL, HTL and B Cell). The chosen epitopes were further analyzed for their interactions with their respective HLA allele binders and TAP cavity interaction.

**Figure 2.**
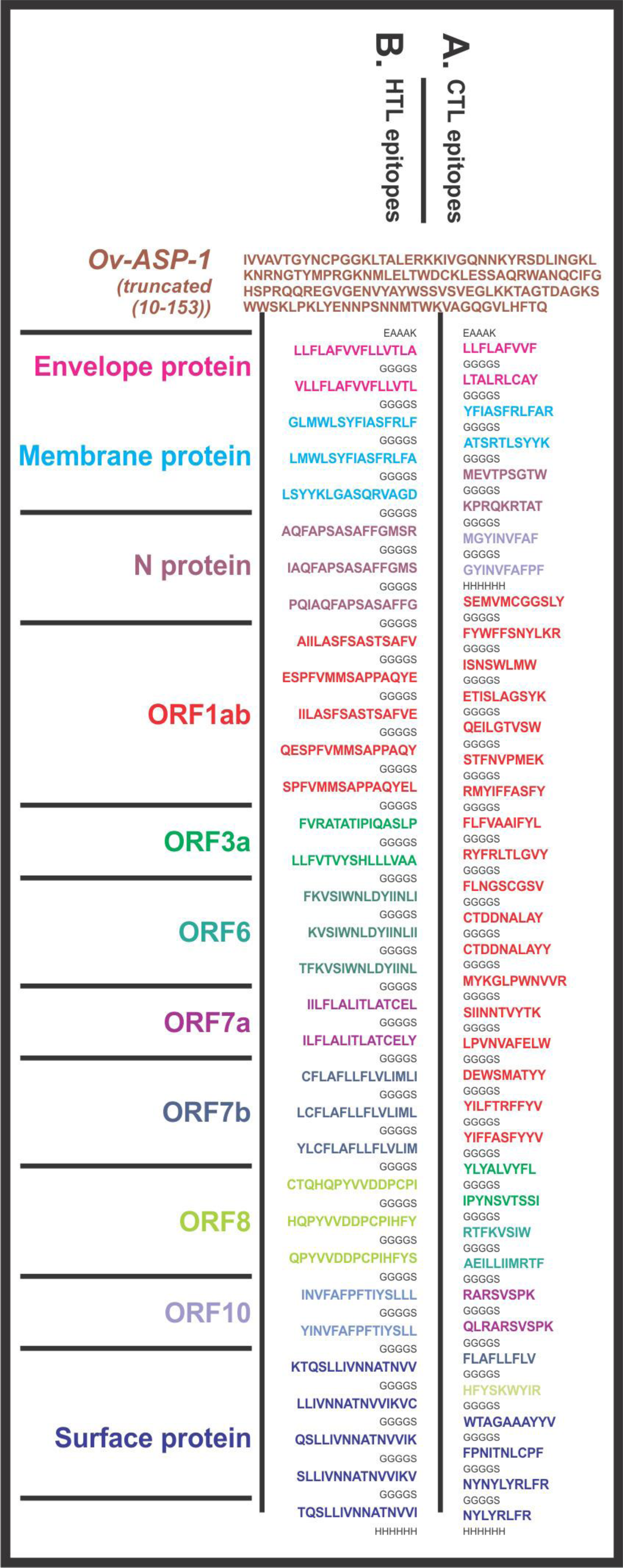
Design of SARS-CoV-2 Multi-Epitope Vaccine (MEVs). (A) CTL and (B) HTL epitopes were linked by the short peptide linker ‘GGGGS’. Truncated (residues 10-153) Onchocerca volvulus activation-associated secreted protein-1 (Ov-ASP-1) has been utilized at the N terminal of both the MEVs. The short peptide EAAAK was used to link Ov-ASP-1 at N terminal. Epitopes from different proteins were colored in different colors. C terminal 6xHis is designed as His tag.

### Molecular interaction analysis of selected epitopes with HLA allele and TAP transporter

#### Tertiary structure modeling of HLA alleles and selected T cell epitopes

The Swiss-model was used for homology modeling of the HLA class I and II allele binders of the chosen epitopes (Arnold et al., 2006). The amino acid sequences of the HLA allele binders were retrieved from Immuno Polymorphism Database (IPD-IMGT/HLA) (https://www.ebi.ac.uk/ipd/imgt/hla/allele.html). Templates for homology modeling were chosen based on the highest amino acid sequence similarity. All the generated HLA allele models had acceptable QMEAN value (cutoff −4.0) (Supplementary table S1). The QMEAN value gives a composite quality estimate involving both global as well as local analysis of the model (Benkert et al., 2008). The PEP-FOLD 2.0 a de novo structure prediction tool at RPBS Web Portal (https://mobyle.rpbs.univ-paris-diderot.fr/cgi-bin/portal.py#forms::PEP-FOLD) was utilized to generate tertiary structures for the chosen CTL and HTL epitopes (Shen et al., 2014).

**Table 1.**
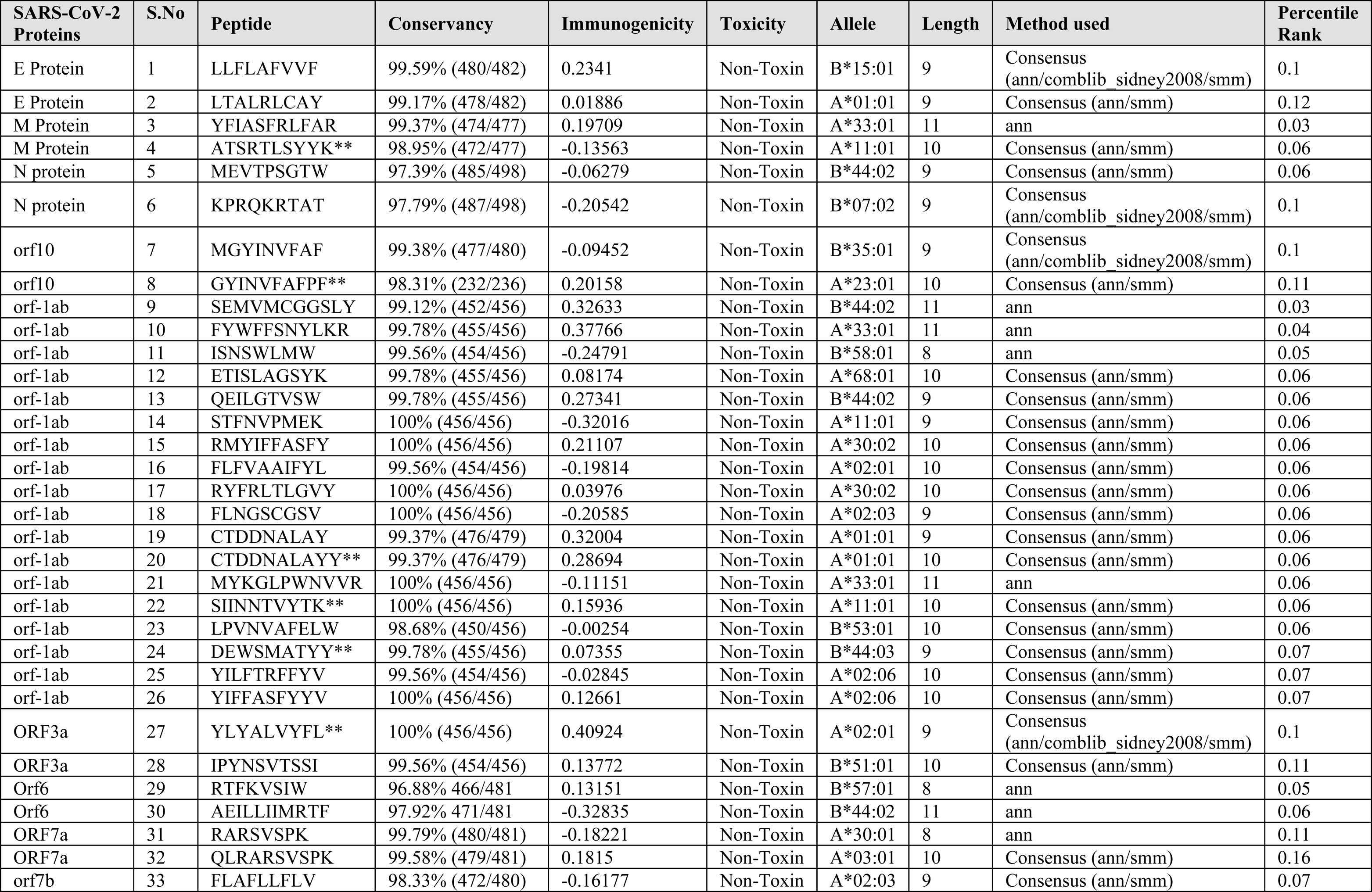
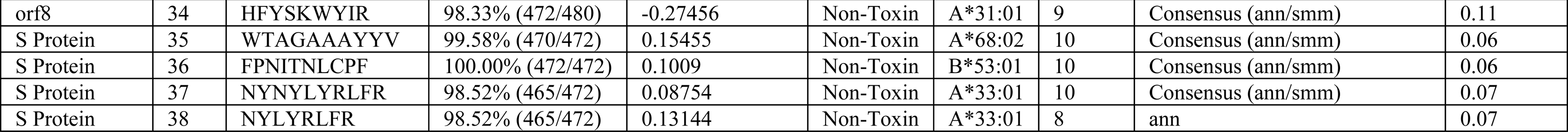
Shortlisted high percentile ranking SARS-CoV-2 CTL epitopes. Selected high percentile CTL epitopes and their respective HLA alleles binders are listed. In-silico analysis has shown all the selected epitopes to be non-toxic (Non-Toxin) as well as they show significant conservancy and high immunogenicity. Six epitopes shown with ** show exact match with the epitopes published by Grifoni et al., 2020, indicate consensus in results.

#### Molecular interaction analysis of chosen CTL and HTL epitopes with HLA alleles

The PatchDock tool (http://bioinfo3d.cs.tau.ac.il/PatchDock/) was utilized for *in-silico* molecular docking study of the selected CTL and HTL epitopes with their respective HLA class I and II allele binders (Bell et al., 2005; Duhovny et al., 2002; Schneidman-Duhovny et al., 2005). PatchDock utilizes an algorithm for unbound (real-life) docking of molecules for protein-protein complex formation. The algorithm carries out the rigid docking, with the surface variability/flexibility implicitly addressed through liberal intermolecular penetration. The algorithm focuses on the (i) initial molecular surface fitting on localized, curvature-based surface patches (ii) use of Geometric Hashing and Pose Clustering for initial transformation detection (iii) computation of shape complementarity utilizing the Distance Transform (iv) efficient steric clash detection and geometric fit scoring based on a multi-resolution shape representation and (v) utilization of biological information by focusing on hot-spot rich surface patches (Bell et al., 2005; Duhovny et al., 2002; Schneidman-Duhovny et al., 2005).

#### Molecular interaction analysis of selected CTL epitopes with TAP transporter

TAP transporter plays an important role in the presentation of CTL epitope. From the cytosol after proteasome processing, the fragmented peptide of foreign protein gets transported to endoplasmic reticulum (ER) through the TAP transporter. From the ER these short peptides reach to Golgi bodies and then get presented on the cell surface (Abele et al., 2004). Molecular interaction study of the chosen CTL epitopes within the TAP transporter cavity was performed by molecular docking study utilizing the PatchDock tool. For accurate prediction, the cryo-EM structure of TAP transporter (PDB ID: 5u1d) was used by removing the antigen from TAP cavity of the original structure (Oldham et al., 2016).

### Design, characterization and molecular interaction analysis of Multi-Epitope Vaccines with immune receptor

#### Design of Multi-Epitope Vaccines

The screened and shortlisted high scoring 38 CTL and 33 HTL epitopes were utilized to design CTL and HTL Multi-Epitope vaccines (Table 1 and 2). Short peptide EAAAK and GGGGS were used as rigid and flexible linkers respectively (Fig.2). The GGGGS linker provides proper conformational flexibility to the vaccine tertiary structure and hence facilitates stable conformation to the vaccine. The EAAAK linker facilitates in domain formation and hence facilitates the vaccine to obtain its final stable structure. Truncated (residues 10-153) Onchocerca volvulus activation-associated secreted protein-1 (Ov-ASP-1) has been utilized as an adjuvant at N terminal of both the CTL and HTL MEVs (MacDonald et al., 2005; Guo et al., 2015; He et al., 2009; Hu et al., 2004; Hajighahramani et al., 2017; Chen et al., 2013; Srivastava et al., 2019; Srivastava et al., 2018).

### Characterization of designed Multi-Epitope Vaccines

#### Physicochemical property analysis of designed MEVs

The ProtParam (https://web.expasy.org/protparam/) tool was utilized to analyze the physiochemical properties of the amino acid sequence of the designed CTL and HTL MEVs (Gasteiger et al., 2005). The ProtParam analysis performs an empirical investigation for the given query amino acid sequence. ProtParam computes various physicochemical properties derived from a given protein sequence.

#### Interferon-gamma inducing epitope prediction

From the designed amino acid sequence of both the MEVs potential interferon-gamma (IFN-γ) epitopes were screened by “IFN epitope” server (http://crdd.osdd.net/raghava/ifnepitope/scan.php) using “Motif and SVM hybrid”, (MERCI: Motif-EmeRging and with Classes-Identification, and SVM: support vector machine) method. The tool predicts peptides from protein sequences having the capacity to induce IFN-gamma release from CD4+ T cells. This module generates overlapping peptides from the query sequence and predicts IFN-gamma inducing peptides. For the screening, IEDB database with 3705 IFN-gamma inducing and 6728 non-inducing MHC class II binders are utilized (Nagpal et al., 2015; Dhanda et al., 2013).

#### MEVs allergenicity and antigenicity prediction

Both the designed MEVs were further analyzed for allergenicity and antigenicity prediction by utilizing the AlgPred (http://crdd.osdd.net/raghava/algpred/submission.html) and the Vaxigen (http://www.ddg-pharmfac.net/vaxijen/VaxiJen/VaxiJen.html) tools respectively (Saha et al., 2006; Doytchinova and Flower 2007). The AlgPred prediction is based on the similarity of already known epitope with any region of the submitted protein. For the screening of allergenicity, the Swiss-prot dataset consisting of 101725 non-allergens and 323 allergens is utilized. The VaxiJen utilizes an alignment-free approach, solely based on the physicochemical properties of the query amino acid sequence. For the prediction of antigenicity, the Bacterial, viral and the tumor protein datasets are used by VaxiJen to derive models for the prediction of whole protein antigenicity. Every set consisted of known 100 antigens and 100 non-antigens.

#### Tertiary structure modeling and refinement of MEVs

The tertiary structure of both the designed CTL and HTL MEVs were generated by homology modeling utilizing the I-TASSER modeling tool (https://zhanglab.ccmb.med.umich.edu/I-TASSER/). The I-TASSER is a tool for protein structure prediction based on the sequence-to-structure-to-function paradigm (Roy et al., 2010). The tool generates three-dimensional (3D) atomic models from multiple threading alignments and iterative structural assembly simulations for a submitted amino acid sequence. I-TASSER works based on the structure templates identified by LOMETS, a meta-server, from the PDB library. I-TASSER only uses the templates of the highest Z-score which is the difference between the raw and average scores in the unit of standard deviation. For each target model, the I-TASSER simulations generate a large ensemble of structural conformations, called decoys. To select the final models, I-TASSER uses the SPICKER program to cluster all the decoys based on the pair-wise structure similarity and reports up to five models. The Normalized Z-score >1 mean a good alignment and vice versa. The Cov represents the coverage of the threading alignment and is equal to the number of aligned residues divided by the length of query protein. Ranking of templet proteins is based on TM-score of the structural alignment between the query structure model and known structures. The RMSD is the RMSD between templet residues and query residues that are structurally aligned by TM-align.

The refinement of both the generated MEV models was performed by ModRefiner (https://zhanglab.ccmb.med.umich.edu/ModRefiner/) and GalaxyRefine tool (http://galaxy.seoklab.org/cgi-bin/submit.cgi?type=REFINE) (Dong et al., 2011). TM-score generated by ModRefiner indicates the structural similarity of the refined model with the original input model. Closer the TM-Score to 1, higher would be the similarity of original and the refined model. RMSD value of the refined model shows the conformational deviation from the initial input models.

The GalaxyRefine tool refines the query tertiary structure by repeated structure perturbation as well as by utilizing the subsequent structural relaxation by the molecular dynamics simulation. The tool GalaxyRefine generates reliable core structures from multiple templates and then re-builds unreliable loops or termini by using an optimization-based refinement method (Ko et al., 2012; Wang et al., 2013; Shin et al., 2014). To avoid any breaks in the 3D model GalaxyRefine uses the triaxial loop closure method. The MolProbity score generated for a given refined model indicates the log-weighted combination of the clash score, the percentage of Ramachandran not favored residues and the percentage of bad side-chain rotamers.

#### Validation of CTL and HTL MEVs refined models

Both the refined CTL and HTL MEV 3D models were further validated by RAMPAGE analysis tool (http://mordred.bioc.cam.ac.uk/~rapper/rampage.php) (S.C. Lovell et al., 2008; Ramakrishnan, C. et al., 1965). The generated Ramachandran plots for the MEV models show the sterically allowed and disallowed residues along with their dihedral psi (ψ) and phi (φ) angles.

#### Linear and Discontinuous B-cell epitope prediction from MEVs

The Ellipro (ElliPro: Antibody Epitope Prediction tool; http://tools.iedb.org/ellipro/) method available at IEDB, was used to screen the linear and the discontinuous B cell epitopes from the MEVs vaccine models. The ElliPro method analyses based on the location of residue in the protein’s 3D structure. The residues lying outside of an ellipsoid covering 90% of the inner core protein residues score highest Protrusion Index (PI) of 0.9; and so on. The discontinuous epitopes predicted by the ElliPro tool are clustered based on the distance “R” in Å between two residue’s centers of mass lying outside of the largest possible ellipsoid. The larger value of R indicates larger distant residues (residue discontinuity) are screened in the epitopes (Kringelum et al., 2012; Ponomarenko et al., 2008).

#### Molecular interaction analysis of MEVs with immunological receptor

##### Molecular docking study of MEVs and TLR-3

Molecular interaction analysis of both the designed MEVs with Toll-Like receptor-3 (TLR-3), was performed by molecular docking and molecular dynamics simulation. Molecular docking was performed by PatchDock server (http://bioinfo3d.cs.tau.ac.il/PatchDock/) (Schneidman-Duhovny et al., 2005). PatchDock utilizes an algorithm for unbound (mimicking real-world environment) docking of molecules for protein-protein complex formation as explained earlier (Bell et al., 2005; Duhovny et al., 2002). For molecular docking, the 3D structure of human TLR-3 ectodomain (ECD) was retrieved from PDB databank (PDB ID: 2A0Z). The study provides dynamical properties of the designed system with MEVs-TLR3 complexes with a guess at the interactions between the molecules, and also it gives ‘exact’ predictions of bulk properties including the hydrogen bond formation and conformation of the molecules forming the complex.

##### Molecular Dynamics (MD) Simulations study of MEVs and TLR-3 complex

The MEVs-TLR3 molecular interactions were further evaluated using molecular dynamics simulations analysis. MD simulation studies were performed for 10 ns by using YASARA (Yet Another Scientifc Artifcial Reality Application) (Krieger, E. and Vriend, G., et al., 2015).The simulations were carried out in an explicit water environment in a dodecahedron simulation box at a constant temperature (298K) and pressure (1 atm) and pH 7.4 with periodic cell boundary condition. The solvated systems were neutralized with counter ions (NaCl) (conc. 0.9 M). The AMBER14 force field have been applied on to the systems during simulation (Maier, J.A., et al., 2015; Case, D.A., et al., 2014). Long-range electrostatic energy and forces were calculated using particle-mesh-based Ewald method (Toukmaji, A., et al., 2000). The solvated structures was minimized by the steepest descent method at a temperature of 298K and a constant pressure. Then the complexes were equilibrated for a 1ns period. After equilibration, a production MD was run for 10 ns at a constant temperature and pressure and time frames were saved at every 10 ps for each simulation. The RMSD and RMSF values for Cα, Back bone and all the atoms of both the MEV complexes were analyzed for each simulation conducted.

### *In-silico* analysis of MEVs for cloning and expression potency

#### Analysis of cDNA of both the MEVs for cloning and expression in the mammalian host cell line

Complementary DNA of both the MEVs, codon-optimized for expression in Mammalian cell line (Human) was generated by Java Codon Adaptation Tool (http://www.jcat.de/). The generated cDNA of both the MEVs was further analyzed by GenScript Rare Codon Analysis Tool (https://www.genscript.com/tools/rare-codon-analysis). The tool analyses the GC content, Codon Adaptation Index (CAI) and the Tandem rare codon frequency for a given cDNA (Morla et al., 2016; Wu et al., 2010). The CAI indicates the possibility of cDNA expression in a chosen expression system. The tandem rare codon frequency indicates the presence of low-frequency codons in the given cDNA.

## RESULTS & DISCUSSION

### Screening of potential epitopes

#### T cell Epitope Prediction

##### Screening of Cytotoxic T lymphocyte (CTL) Epitope

Cytotoxic T lymphocyte (CTL) epitopes were screened by “MHC-I Binding Predictions” and “MHC-I Binding Predictions” IEDB tools. These epitopes are shortlisted based on the total amount of cleavage site in the protein, low IC(50) (nM) value for epitope-HLA class I allele pairs, and for binding to TAP cavity. The 38 epitopes predicted by “MHC-I Binding Predictions” tool with the highest “Percentile Rank” were shortlisted for multi-epitope vaccine design and are listed in table 1. Rest 101 epitopes-HLA I allele pairs are listed in Supplementary table S8. The 67 epitopes-HLA I allele pairs predicted by “MHC-I Processing Predictions” tool with the highest “Total score” are listed in Supplementary table 9.

The immunogenicity of the shortlisted CTL epitopes was also determined and are mentioned in table 1, Supplementary table S8, and S9. The higher immunogenicity score indicates the greater immunogenic potential of the given epitope.

##### Screening of Helper T lymphocyte (HTL) epitopes

The screening of helper T lymphocyte (HTL) epitopes from eleven different proteins of SARS-CoV-2 was performed based on “Percentile rank”. The smaller the value of percentile rank the higher would be the affinity of the peptide with its respective HLA allele binders. The 33 epitopes with high percentile ranking were shortlisted (Table 2). Another 180 potential HTL cell epitopes-HLA allele II pairs with high Percentile rank, screened in our study are listed in Supplementary table S10.

**Table 2.**
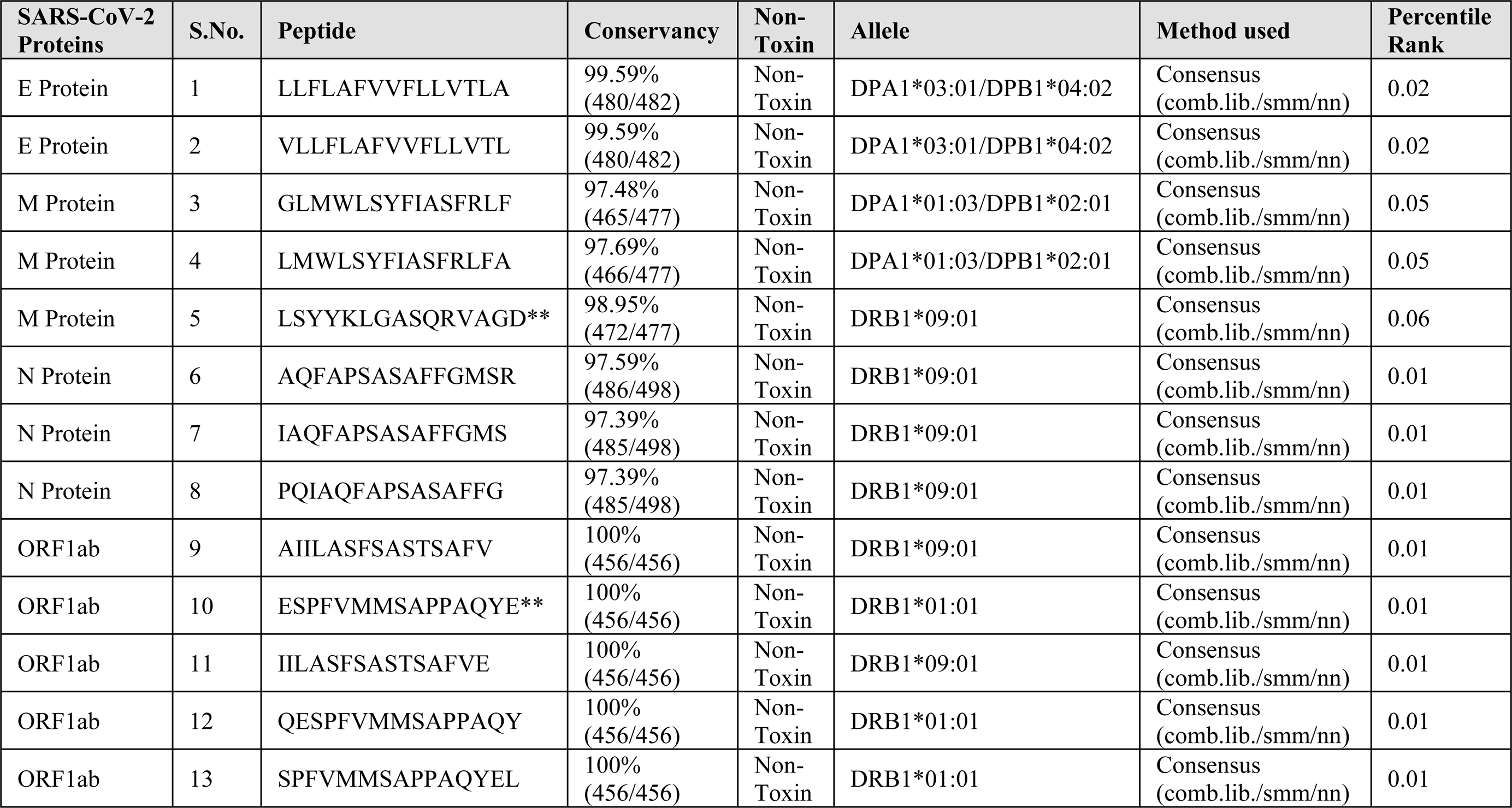
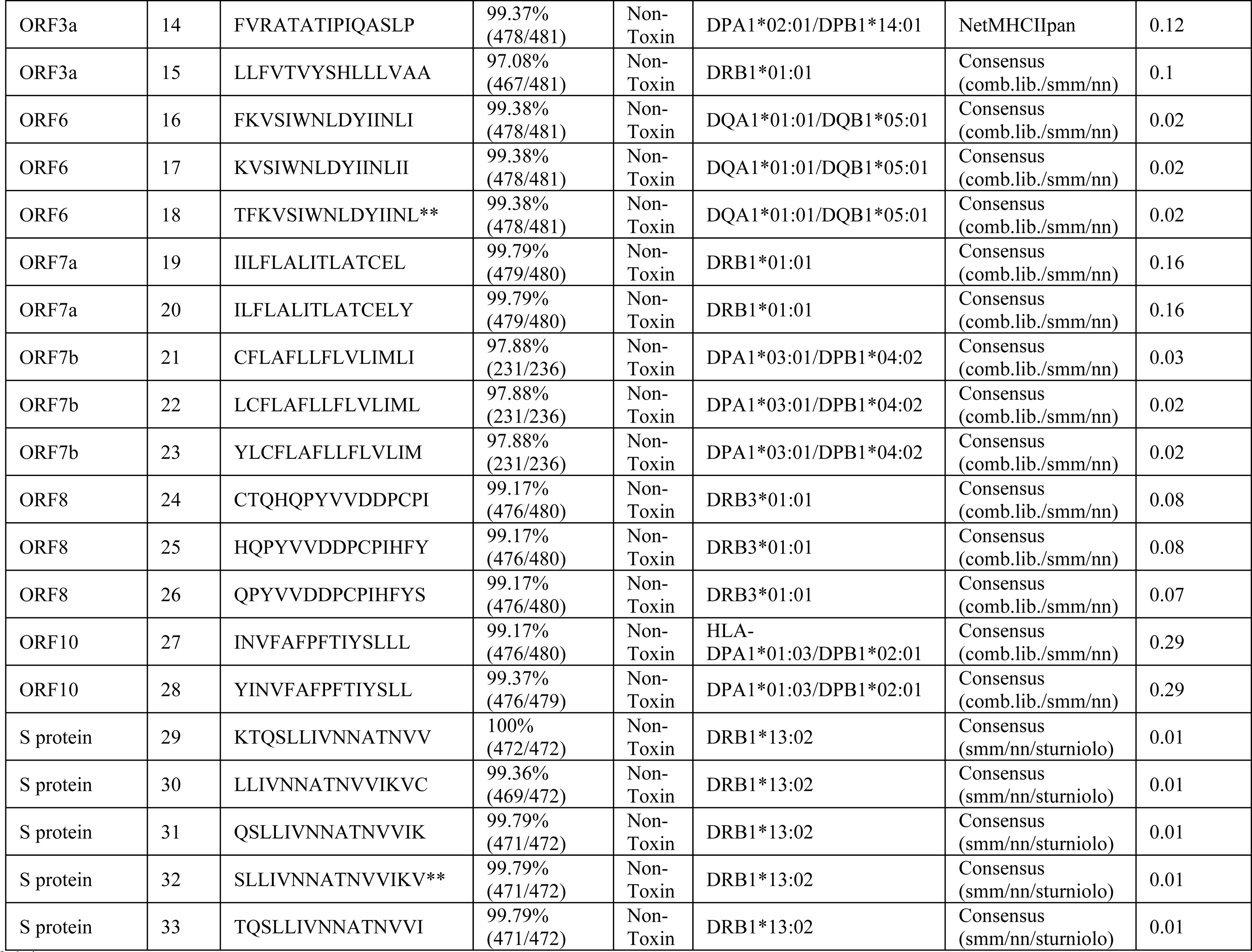
Shortlisted high scoring SARS-CoV-2 HTL epitopes. Selected high scoring HTL epitopes and their respective HLA alleles binders are listed. In-silico analysis has shown all the selected epitopes to be non-toxic (Non-Toxin) as well as they show significant conservancy. Four epitopes shown with ** show exact match with the epitopes published by Grifoni et al., 2020, indicate consensus in results.

##### Population Coverage by CTL and HTL epitopes

The population coverage by the shortlisted epitopes was also studied, in particular involving China, France, Italy, United States of America, South Asia, East Asia, Northeast Asia, and the Middle East. From this study, we may conclude that the combined use of all the shortlisted CTL and HTL epitopes would have an average worldwide population coverage as high as 96.10%, with a standard deviation of 23.74 (Table 3).

**Table 3.**
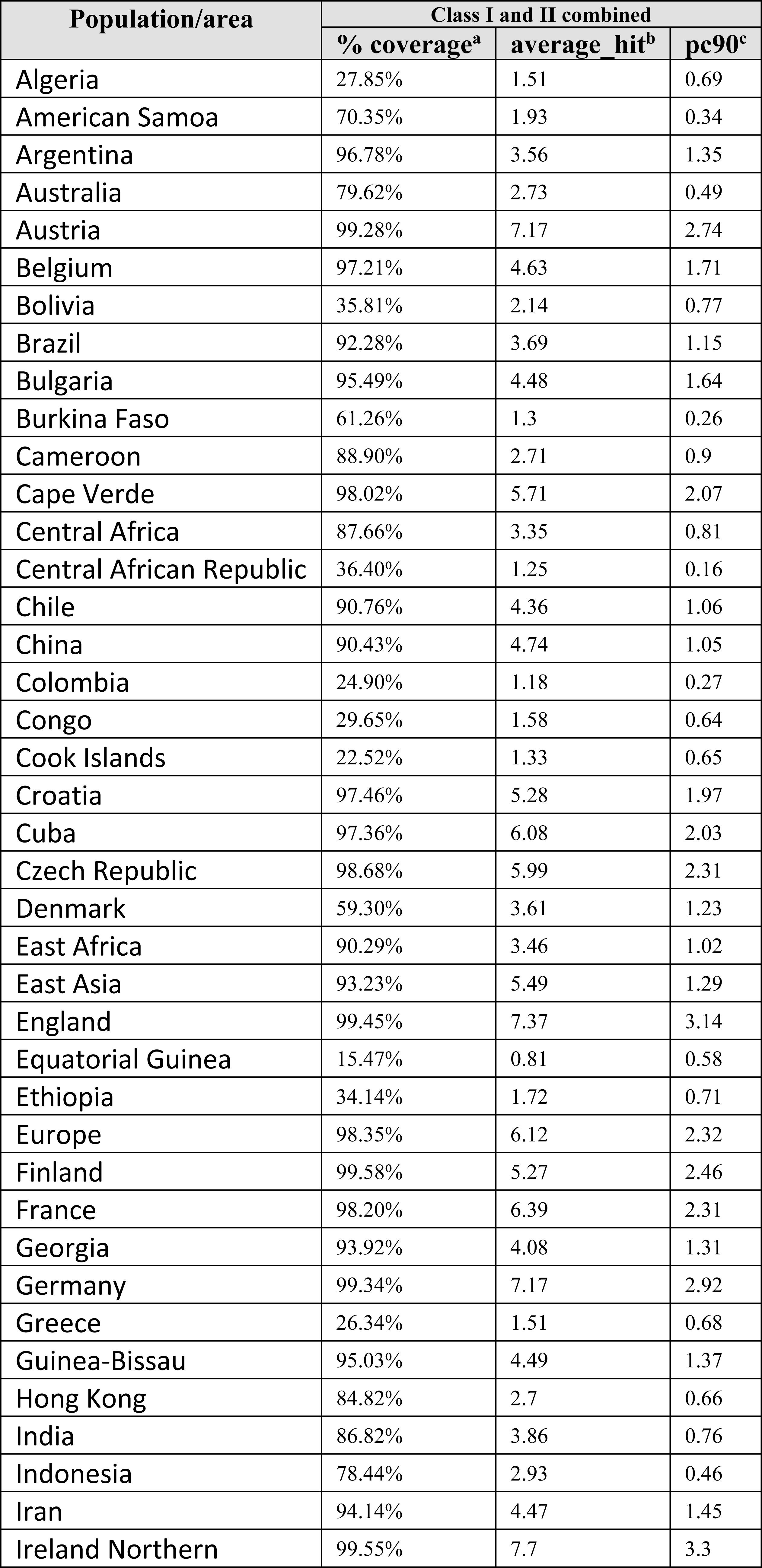
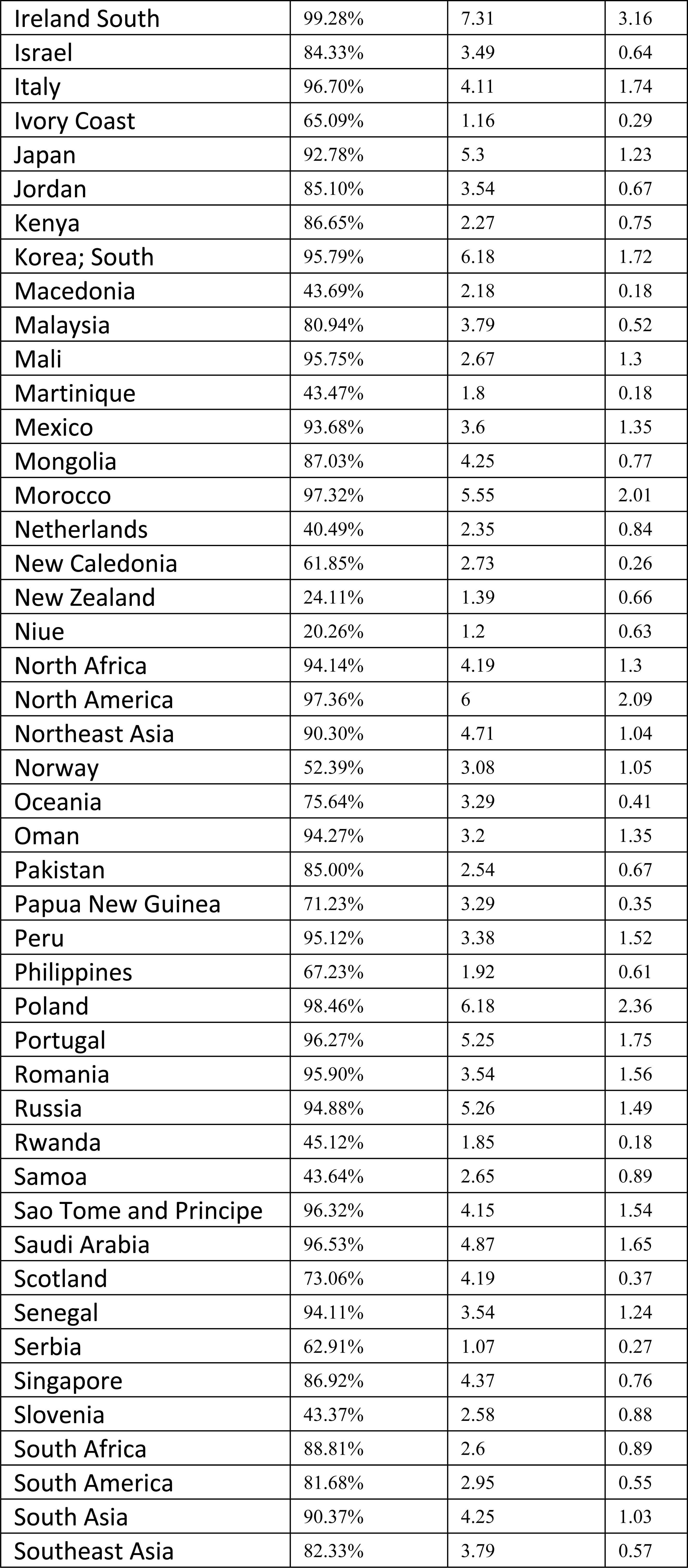
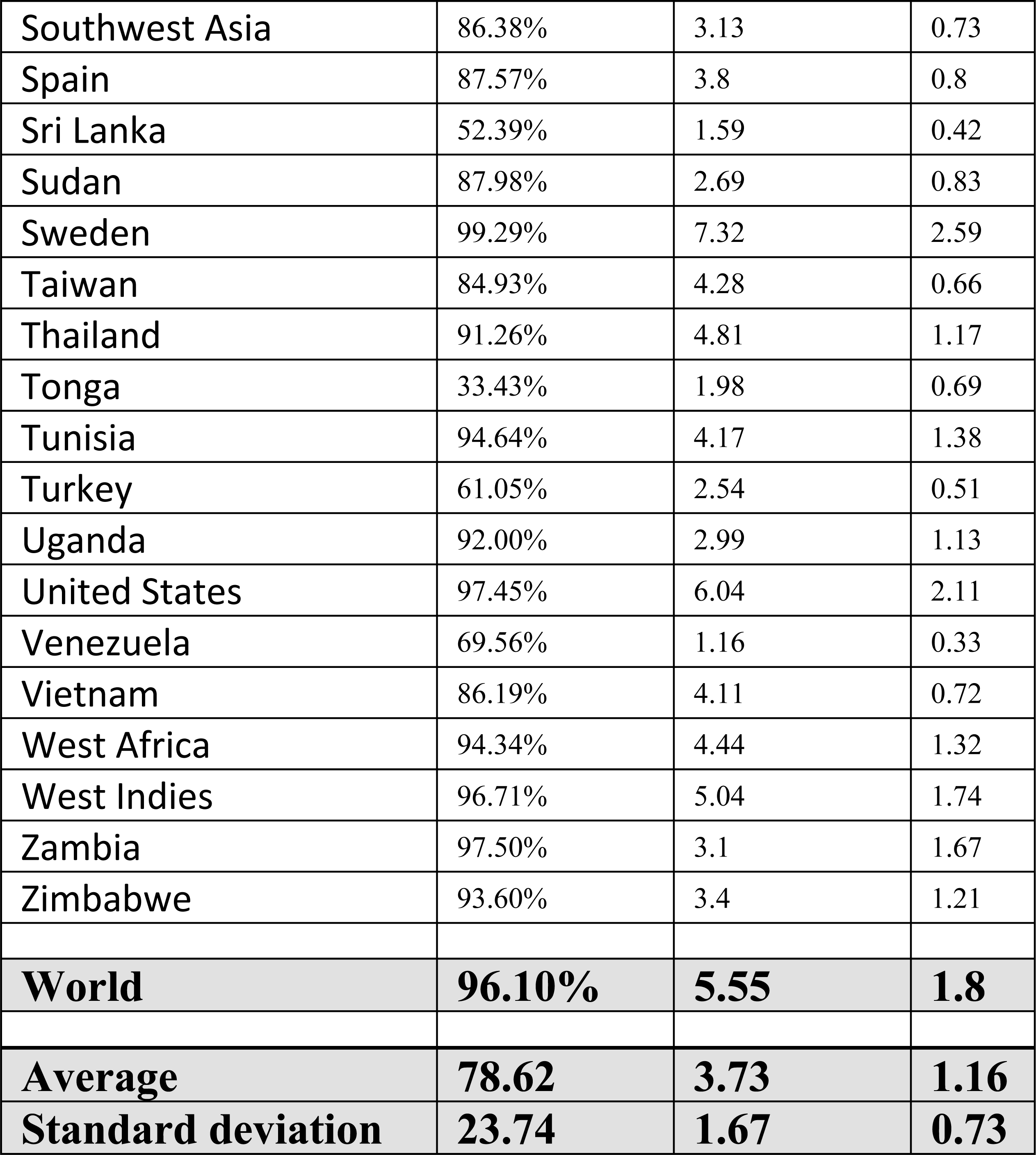
World population coverage by the shortlisted SARS-CoV-2 CTL and HTL epitopes combined. With a standard deviation of 23.74 on an average 96.10 % of the world population could be covered by the joint administration of selected CTL and HTL epitopes (given in Table 1 and 2) as vaccine candidates.

| Population/area | Class I and II combined |  |  |
| --- | --- | --- | --- |
|  | % coverage<sup>a</sup> | average_hit<sup>b</sup> | pc90<sup>c</sup> |
| Algeria | 27.85% | 1.51 | 0.69 |
| American Samoa | 70.35% | 1.93 | 0.34 |
| Argentina | 96.78% | 3.56 | 1.35 |
| Australia | 79.62% | 2.73 | 0.49 |
| Austria | 99.28% | 7.17 | 2.74 |
| Belgium | 97.21% | 4.63 | 1.71 |
| Bolivia | 35.81% | 2.14 | 0.77 |
| Brazil | 92.28% | 3.69 | 1.15 |
| Bulgaria | 95.49% | 4.48 | 1.64 |
| Burkina Faso | 61.26% | 1.3 | 0.26 |
| Cameroon | 88.90% | 2.71 | 0.9 |
| Cape Verde | 98.02% | 5.71 | 2.07 |
| Central Africa | 87.66% | 3.35 | 0.81 |
| Central African Republic | 36.40% | 1.25 | 0.16 |
| Chile | 90.76% | 4.36 | 1.06 |
| China | 90.43% | 4.74 | 1.05 |
| Colombia | 24.90% | 1.18 | 0.27 |
| Congo | 29.65% | 1.58 | 0.64 |
| Cook Islands | 22.52% | 1.33 | 0.65 |
| Croatia | 97.46% | 5.28 | 1.97 |
| Cuba | 97.36% | 6.08 | 2.03 |
| Czech Republic | 98.68% | 5.99 | 2.31 |
| Denmark | 59.30% | 3.61 | 1.23 |
| East Africa | 90.29% | 3.46 | 1.02 |
| East Asia | 93.23% | 5.49 | 1.29 |
| England | 99.45% | 7.37 | 3.14 |
| Equatorial Guinea | 15.47% | 0.81 | 0.58 |
| Ethiopia | 34.14% | 1.72 | 0.71 |
| Europe | 98.35% | 6.12 | 2.32 |
| Finland | 99.58% | 5.27 | 2.46 |
| France | 98.20% | 6.39 | 2.31 |
| Georgia | 93.92% | 4.08 | 1.31 |
| Germany | 99.34% | 7.17 | 2.92 |
| Greece | 26.34% | 1.51 | 0.68 |
| Guinea-Bissau | 95.03% | 4.49 | 1.37 |
| Hong Kong | 84.82% | 2.7 | 0.66 |
| India | 86.82% | 3.86 | 0.76 |
| Indonesia | 78.44% | 2.93 | 0.46 |
| Iran | 94.14% | 4.47 | 1.45 |
| Ireland Northern | 99.55% | 7.7 | 3.3 |
| Ireland South | 99.28% | 7.31 | 3.16 |
| Israel | 84.33% | 3.49 | 0.64 |
| Italy | 96.70% | 4.11 | 1.74 |
| Ivory Coast | 65.09% | 1.16 | 0.29 |
| Japan | 92.78% | 5.3 | 1.23 |
| Jordan | 85.10% | 3.54 | 0.67 |
| Kenya | 86.65% | 2.27 | 0.75 |
| Korea; South | 95.79% | 6.18 | 1.72 |
| Macedonia | 43.69% | 2.18 | 0.18 |
| Malaysia | 80.94% | 3.79 | 0.52 |
| Mali | 95.75% | 2.67 | 1.3 |
| Martinique | 43.47% | 1.8 | 0.18 |
| Mexico | 93.68% | 3.6 | 1.35 |
| Mongolia | 87.03% | 4.25 | 0.77 |
| Morocco | 97.32% | 5.55 | 2.01 |
| Netherlands | 40.49% | 2.35 | 0.84 |
| New Caledonia | 61.85% | 2.73 | 0.26 |
| New Zealand | 24.11% | 1.39 | 0.66 |
| Niue | 20.26% | 1.2 | 0.63 |
| North Africa | 94.14% | 4.19 | 1.3 |
| North America | 97.36% | 6 | 2.09 |
| Northeast Asia | 90.30% | 4.71 | 1.04 |
| Norway | 52.39% | 3.08 | 1.05 |
| Oceania | 75.64% | 3.29 | 0.41 |
| Oman | 94.27% | 3.2 | 1.35 |
| Pakistan | 85.00% | 2.54 | 0.67 |
| Papua New Guinea | 71.23% | 3.29 | 0.35 |
| Peru | 95.12% | 3.38 | 1.52 |
| Philippines | 67.23% | 1.92 | 0.61 |
| Poland | 98.46% | 6.18 | 2.36 |
| Portugal | 96.27% | 5.25 | 1.75 |
| Romania | 95.90% | 3.54 | 1.56 |
| Russia | 94.88% | 5.26 | 1.49 |
| Rwanda | 45.12% | 1.85 | 0.18 |
| Samoa | 43.64% | 2.65 | 0.89 |
| Sao Tome and Principe | 96.32% | 4.15 | 1.54 |
| Saudi Arabia | 96.53% | 4.87 | 1.65 |
| Scotland | 73.06% | 4.19 | 0.37 |
| Senegal | 94.11% | 3.54 | 1.24 |
| Serbia | 62.91% | 1.07 | 0.27 |
| Singapore | 86.92% | 4.37 | 0.76 |
| Slovenia | 43.37% | 2.58 | 0.88 |
| South Africa | 88.81% | 2.6 | 0.89 |
| South America | 81.68% | 2.95 | 0.55 |
| South Asia | 90.37% | 4.25 | 1.03 |
| Southeast Asia | 82.33% | 3.79 | 0.57 |
| Southwest Asia | 86.38% | 3.13 | 0.73 |
| Spain | 87.57% | 3.8 | 0.8 |
| Sri Lanka | 52.39% | 1.59 | 0.42 |
| Sudan | 87.98% | 2.69 | 0.83 |
| Sweden | 99.29% | 7.32 | 2.59 |
| Taiwan | 84.93% | 4.28 | 0.66 |
| Thailand | 91.26% | 4.81 | 1.17 |
| Tonga | 33.43% | 1.98 | 0.69 |
| Tunisia | 94.64% | 4.17 | 1.38 |
| Turkey | 61.05% | 2.54 | 0.51 |
| Uganda | 92.00% | 2.99 | 1.13 |
| United States | 97.45% | 6.04 | 2.11 |
| Venezuela | 69.56% | 1.16 | 0.33 |
| Vietnam | 86.19% | 4.11 | 0.72 |
| West Africa | 94.34% | 4.44 | 1.32 |
| West Indies | 96.71% | 5.04 | 1.74 |
| Zambia | 97.50% | 3.1 | 1.67 |
| Zimbabwe | 93.60% | 3.4 | 1.21 |
| **World** | **96.10%** | **5.55** | **1.8** |
| **Average** | **78.62** | **3.73** | **1.16** |
| **Standard deviation** | **23.74** | **1.67** | **0.73** |

### B Cell epitope prediction

#### Sequence-based B Cell epitope prediction

To screen B cell epitopes we utilized the Bepipred Linear Epitope Prediction method. In our study, we screened 12 B cell epitopes, from eleven SARS-CoV-2 proteins, which show partial or complete overlap with shortlisted CTL and HTL epitope (Table 4). Another 206 B Cell epitope, with the epitope length of at least four amino acids and maximum 20 amino acids were screened and are listed in Supplementary table S11.

**Table 4.**
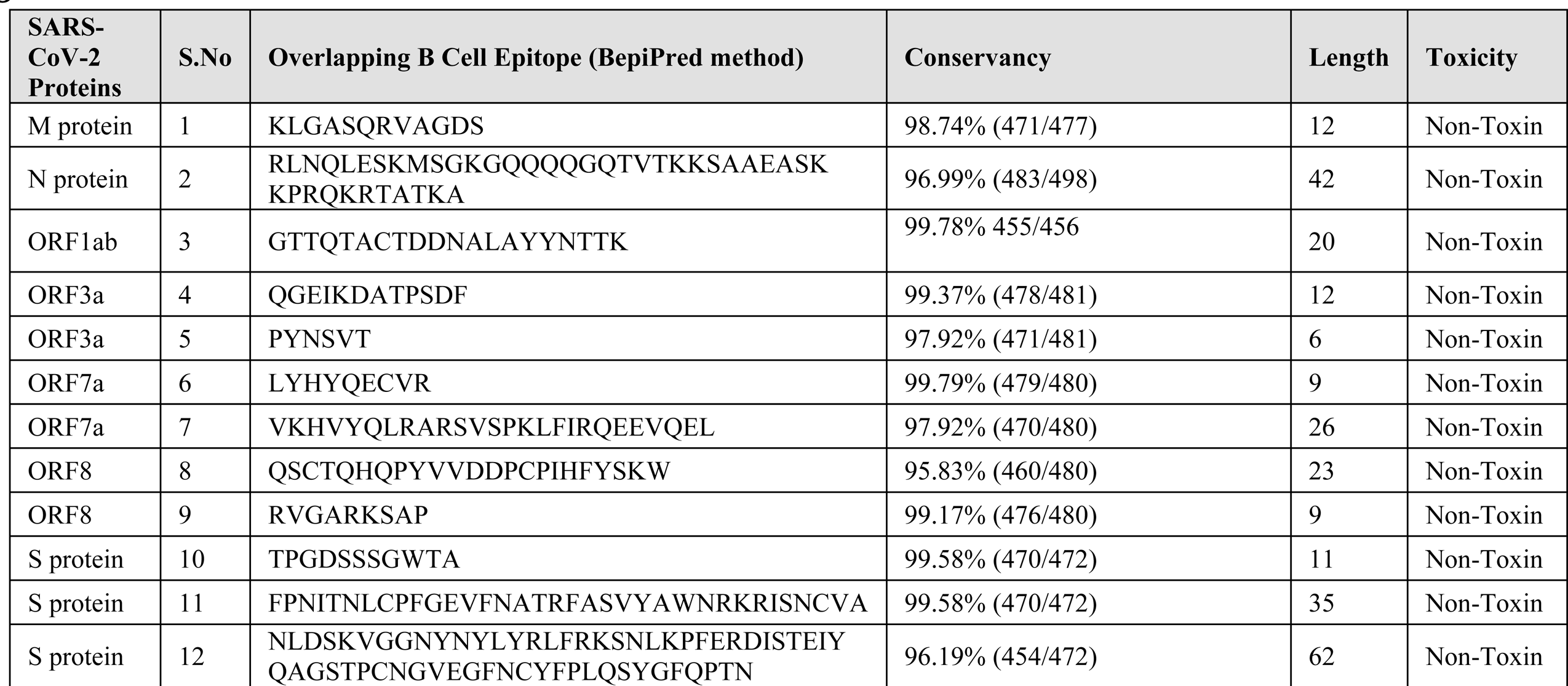
Shortlisted SARS-CoV-2 B Cell epitopes. BepiPred Linear B Cell epitopes showing sequence overlap with CTL and HTL epitopes are shortlisted. In-silico analysis has shown all the selected epitopes to be non-toxic (Non-Toxin) as well as they show significant amino acid sequence conservancy.

### Characterization of potential epitopes

#### Epitope conservation analysis

Sequence conservation analysis of the screened CTL, HTL and B cell epitopes have shown highly conserved nature of the shortlisted epitopes. Both the CTL epitopes and the HTL epitope were found to be significantly conserved with their 100% amino acid sequence amongst the NCBI retrieved protein sequences of SARS-CoV-2 (CTL epitopes 96.88% to 100% conserved and HTL epitopes were 97.08% to 100% conserved (Table 1, 2 4, Supplementary table S8, S9, S10 & S11).

#### Epitope toxicity prediction

Toxicity analysis of all the screened CTL, HTL and B Cell epitopes was also performed. The ToxinPred study of all the shortlisted epitopes shows that they all are non-toxic in nature (Table 1, 2 4, Supplementary table S8, S9, S10 & S11).

#### Overlapping residue analysis

Amino acid sequence overlap analysis amongst the shortlisted CTL, HTL and B cell epitopes from eleven SARS-CoV-2 proteins was performed by the Multiple Sequence Alignment (MSA) analysis tool Clustal Omega. The analysis has shown that several epitopes of CTL, HTL and B cell were having amino acid sequences overlap. The CTL, HTL and B cell epitopes having two or more than two amino acid residues overlap are shown in Fig.2.

#### Epitope selected for molecular interaction study with HLA allele and TAP transporter

The epitopes showing overlap amongst all the three types of epitopes i.e CTL, HTL and B cell epitopes have been encircled in Fig.2 and are chosen for further study for their interaction with HLA allele and TAP (Transporter Associated with Antigen Processing) transporter.

### Molecular interaction analysis of selected epitopes with HLA allele and TAP transporter

#### Molecular interaction analysis of chosen CTL and HTL epitopes with HLA alleles

The molecular docking study of chosen CTL and HTL epitopes with their respective HLA class I and II allele binders was performed by PatchDock tool. The study revealed a significant molecular interaction between all the chosen epitopes and their HLA allele binders showing multiple hydrogen bond formations (Fig.3). Furthermore, the B-factor analysis of all the epitope-HLA allele complexes has also shown the epitope ligand to have stable (blue) binding conformation in complex with the HLA allele molecule (VIBGYOR color

**Figure 3.**
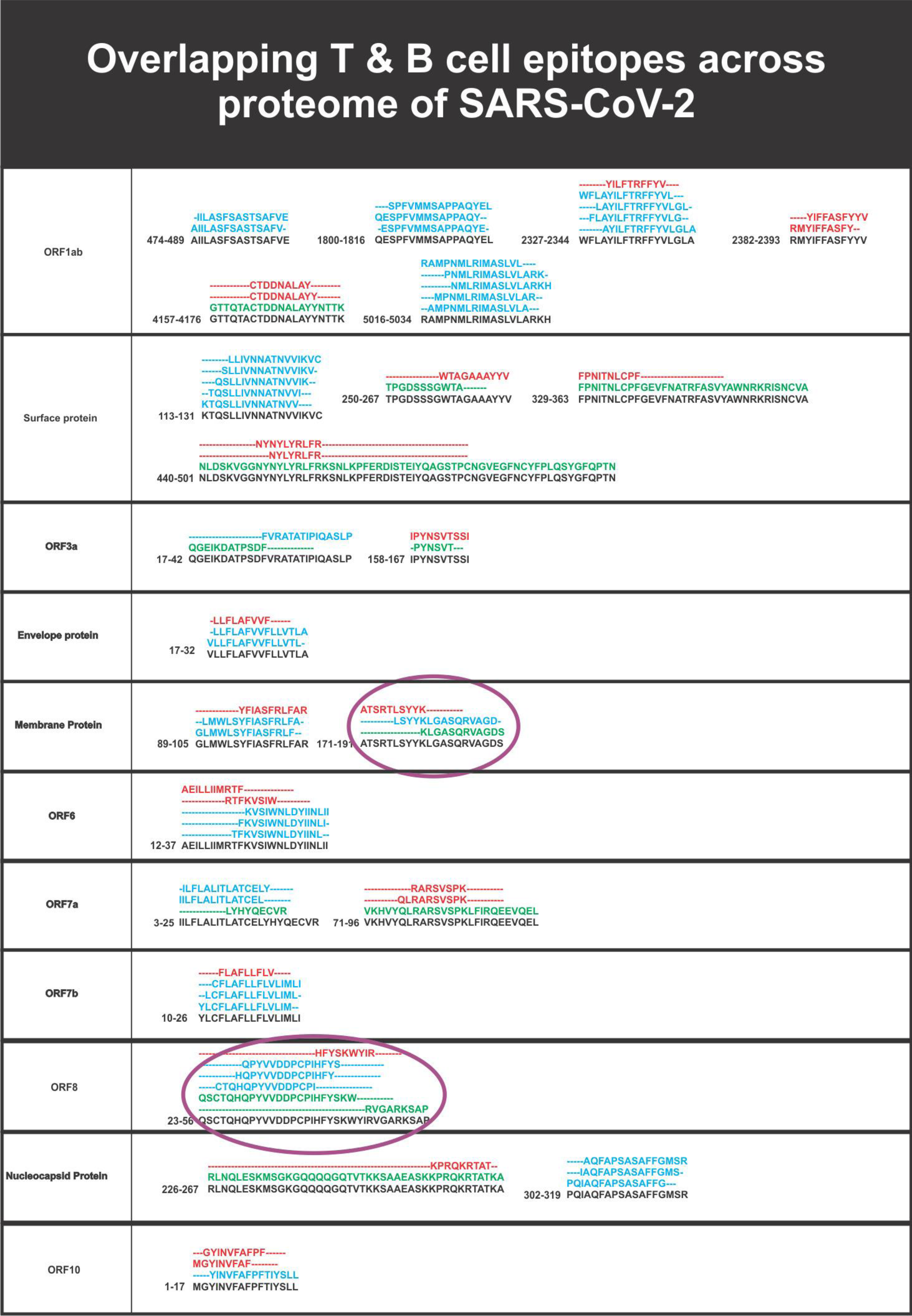
Overlapping SARS-CoV-2 CTL, HTL and B cell epitopes. Multiple sequence alignment performed by Clustal Omega at EBI to identify the consensus overlapping regions of CTL (red), HTL (blue) and B cell epitopes (green) amongst shortlisted epitopes. Epitopes with overlapping regions amongst all the three types of epitopes (CTL, HTL and B Cell epitopes) were chosen for further studies (encircled).

**Figure 4.**
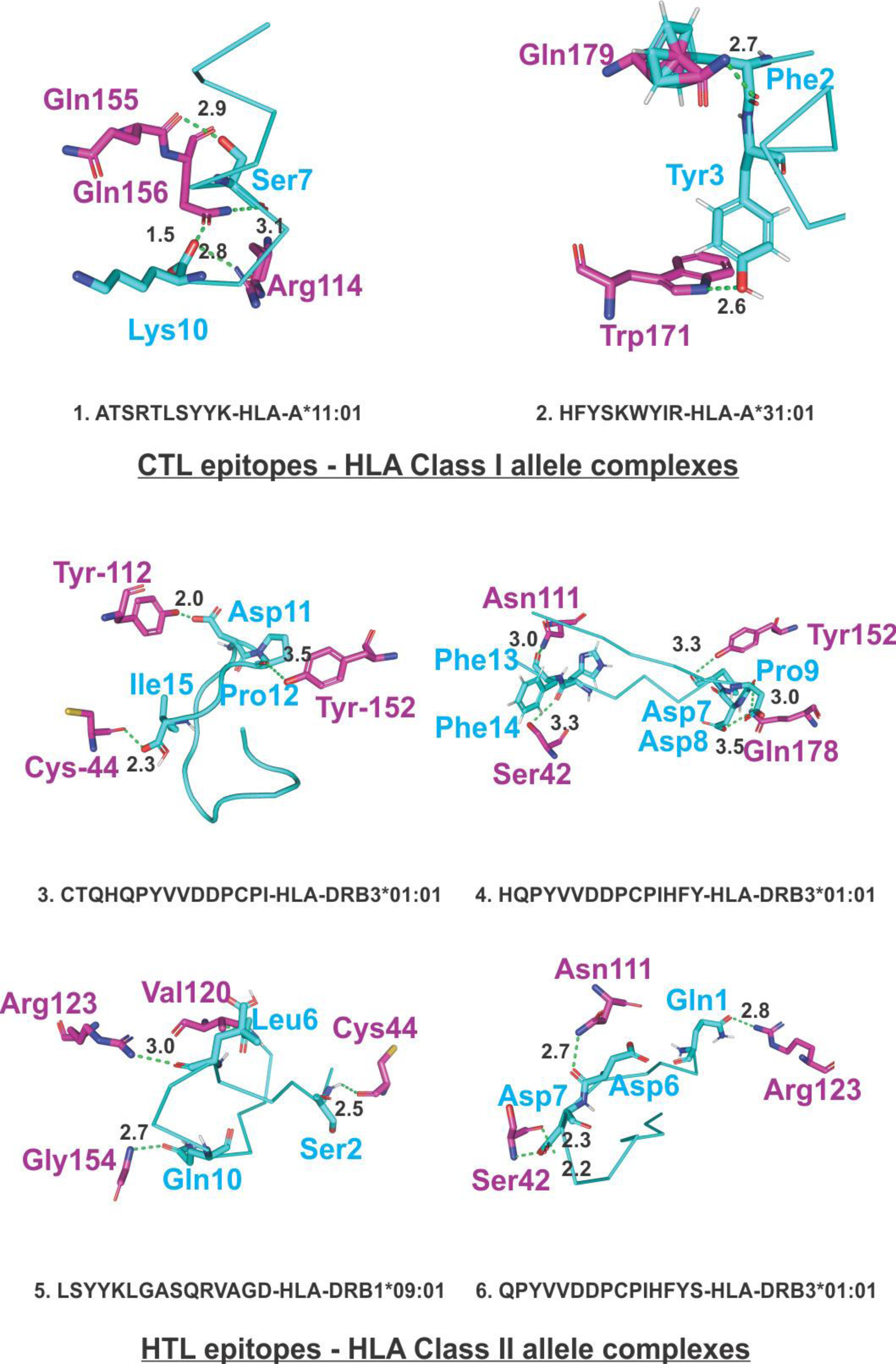
Molecular Docking analysis of SARS-CoV-2 CTL epitopes and HLA alleles. Molecular docking of chosen CTL and HTL epitopes (cyan sticks) with binding amino acid residues of their respective HLA class I and class II allele binders (magenta sticks). The study shows the docked complexes to form a stable complex with multiple hydrogen bonds (green dots, lengths in Angstroms) formation. Images were generated by the PyMOL Molecular Graphics System, Version 2.0 Schrödinger, LLC.

#### Molecular interaction analysis of selected CTL epitopes with TAP transporter

The molecular docking interaction analysis of the chosen CTL epitopes with the TAP transporter cavity has shown a significantly strong molecular interaction with several hydrogen bonds formation at different sites of the TAP transporter cavity. Two sites of interaction were of particular interest, one closer to the cytoplasmic end and another closer to the ER lumen (Fig.5). This study confirms the feasibility of transportation of chosen CTL epitopes from the cytoplasm to the ER lumen which is an essential event for the representation of epitope by the HLA allele molecules on the surface of antigen-presenting cells.

**Figure 5.**
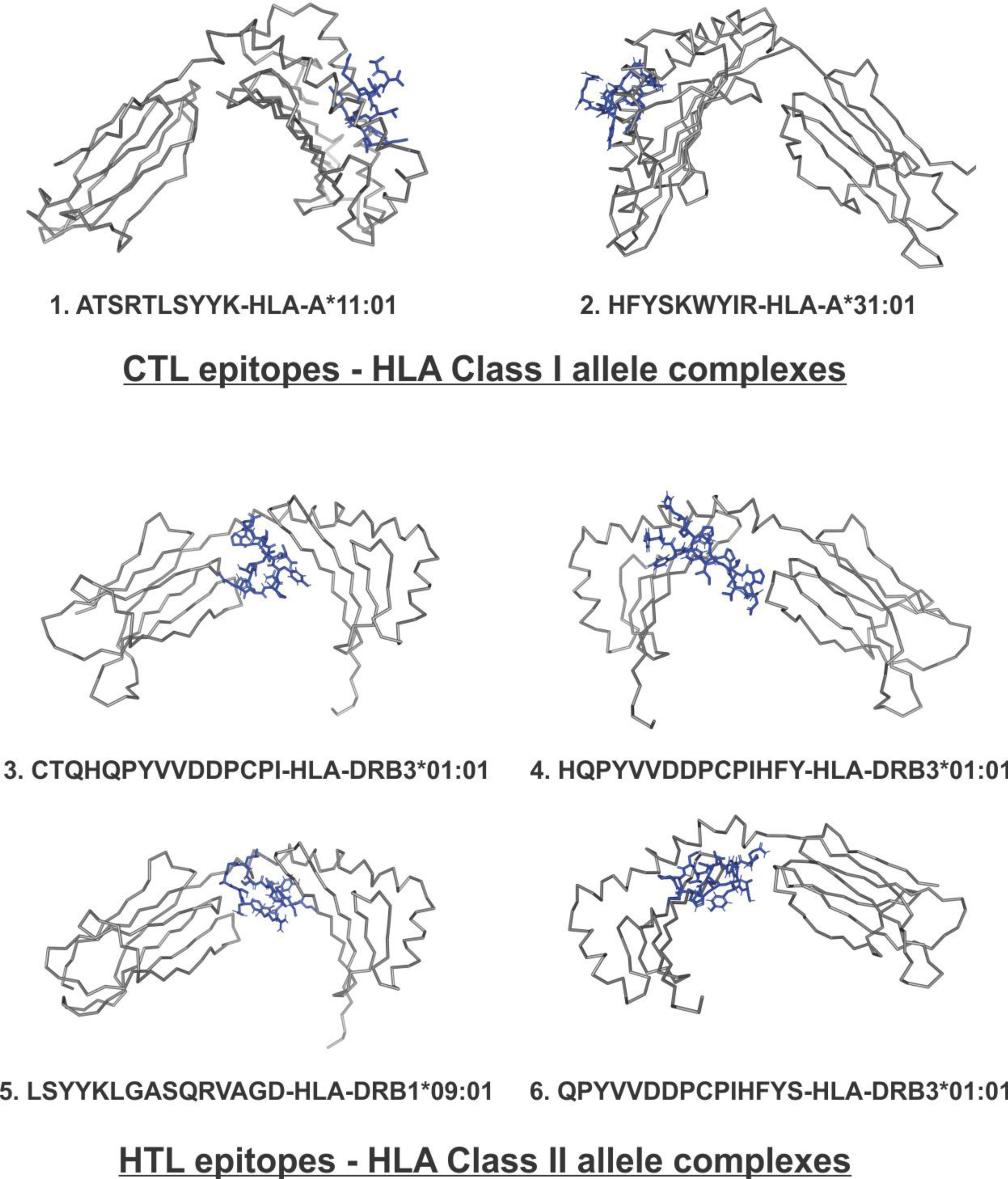
B-Factor of CTL and HTL epitope in complex with HLA class I and II allele. CTL and HTL Epitopes are shown in sticks and HLA Class I and II alleles are shown in ribbon. The HLA alleles are shown in gray. The regions of the epitope in the complex are shown in a rainbow (VIBGYOR), the regions in blue being very stable and the region towards red being relatively unstable. In the complexes shown above, most of the regions of epitopes are in blue indicating the complexes to be highly stable.

### Characterization and molecular interaction analysis of designed Multi-Epitope Vaccines with immune receptor

#### Characterization of designed Multi-Epitope Vaccines

##### Physicochemical property analysis of designed MEVs

ProtParam analysis for both the CTL and HTL MEVs was performed to analyze their physiochemical properties. The empirical physiochemical properties of the CTL and HTL MEVs are given in the table 5. The aliphatic index and grand average of hydropathicity (GRAVY) of both the MEVs indicate the globular and hydrophilic nature of both the MEVs. The instability index score of both the MEVs indicates the stable nature of the protein molecules (Table 5).

**Table 5.**
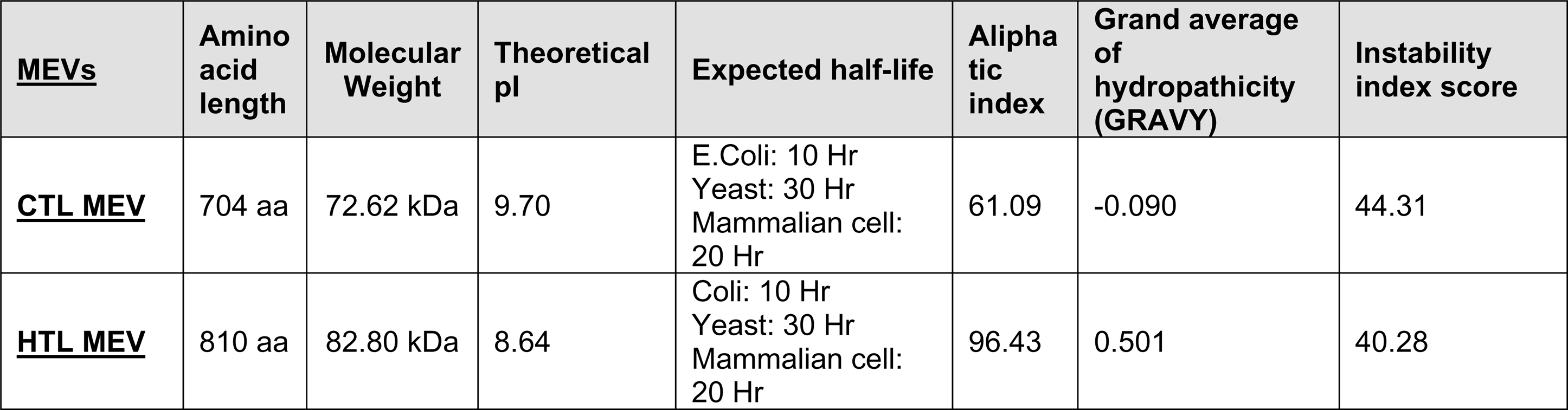
Physicochemical property analysis based on amino acid sequence of designed CTL and HTL multi-epitope vaccine.

##### Interferon-gamma inducing epitope prediction

Interferon-gamma (IFN-γ) inducing epitopes are involved in both the adaptive as well as in the innate immune response. The IFN-γ inducing 15 mer peptide epitopes were screened from the amino acid sequence of CTL and HTL MEVs by utilizing the IFNepitope server. A total of 20 CTL MEV and 20 HTL MEV INF-γ inducing POSITIVE epitopes with a score of 1 or more than 1 were shortlisted (Supplementary table S2).

##### MEVs allergenicity and antigenicity prediction

Both the CTL and HTL MEVs were found to be NON-ALLERGEN by the AlgPred analysis (scoring −0.95185601 and −1.1293352 respectively; threshold being −0.4). The CTL and HTL MEVs were also analyzed by VaxiJen to be probable ANTIGENS (prediction score 0.4485 and 0.4215 respectively; default threshold being 0.4). Hence with the mentioned analysis tools both the CTL and HTL MEVs are predicted to be non-allergic as well as potentially antigenic in nature.

##### Tertiary structure modeling and refinement of MEVs

3D homology models were generated for both the CTL and HTL MEVs by utilizing the I-TASSER modeling tool (Fig.7). The models were generated for CTL Multi-epitope vaccine (PDB hit 5n8pA, Norm. Z-score of 1.49, Cov of 0.92, TM-score of 0.916 and RMSD of 1.04 Å) and HTL Multi-epitope vaccine (PDB hit 5n8pA, Norm. Z-score of 1.52, Cov of 0.97, TM-score of 0.916 and RMSD of 1.04 Å). Both the generated CTL and HTL 3D models were further refined by ModRefiner to fix any gaps and then followed by GalaxyRefine refinement. The refinement by ModRefiner showed the TM-score of 0.9189 and 0.9498 for the CTL and HTL models respectively, hence being close to 1, the initial and the refined models were structurally similar. After refinement, the RMSD for CTL and HTL models with respect to the initial model was 3.367Å and 2.318Å respectively. Further, both the CTL and HTL MEVs models were refined by GalaxyRefine and model 1 was chosen based on best scorings parameters. The CTL MEV model refinement output model (Rama favoured was 83.6%, GDT-HA was 0.9371, RMSD was 0.459, MolProbity was 2.539, Clash score was 23.2, and Poor rotamers was 1.8) and the HTL MEV model refinement output model (Rama favoured was 87.7%, GDT-HA was 0.9552, RMSD was 0.402, MolProbity was 2.537, Clash score was 27.9, and Poor rotamers was 1.6) show a well-refined and acceptable models for both of the MEVs. After refinement, all the mentioned parameters have found to be improved significantly in comparison to the initial CTL and HTL MEV models (Supplementary table S3).

**Figure 6.**
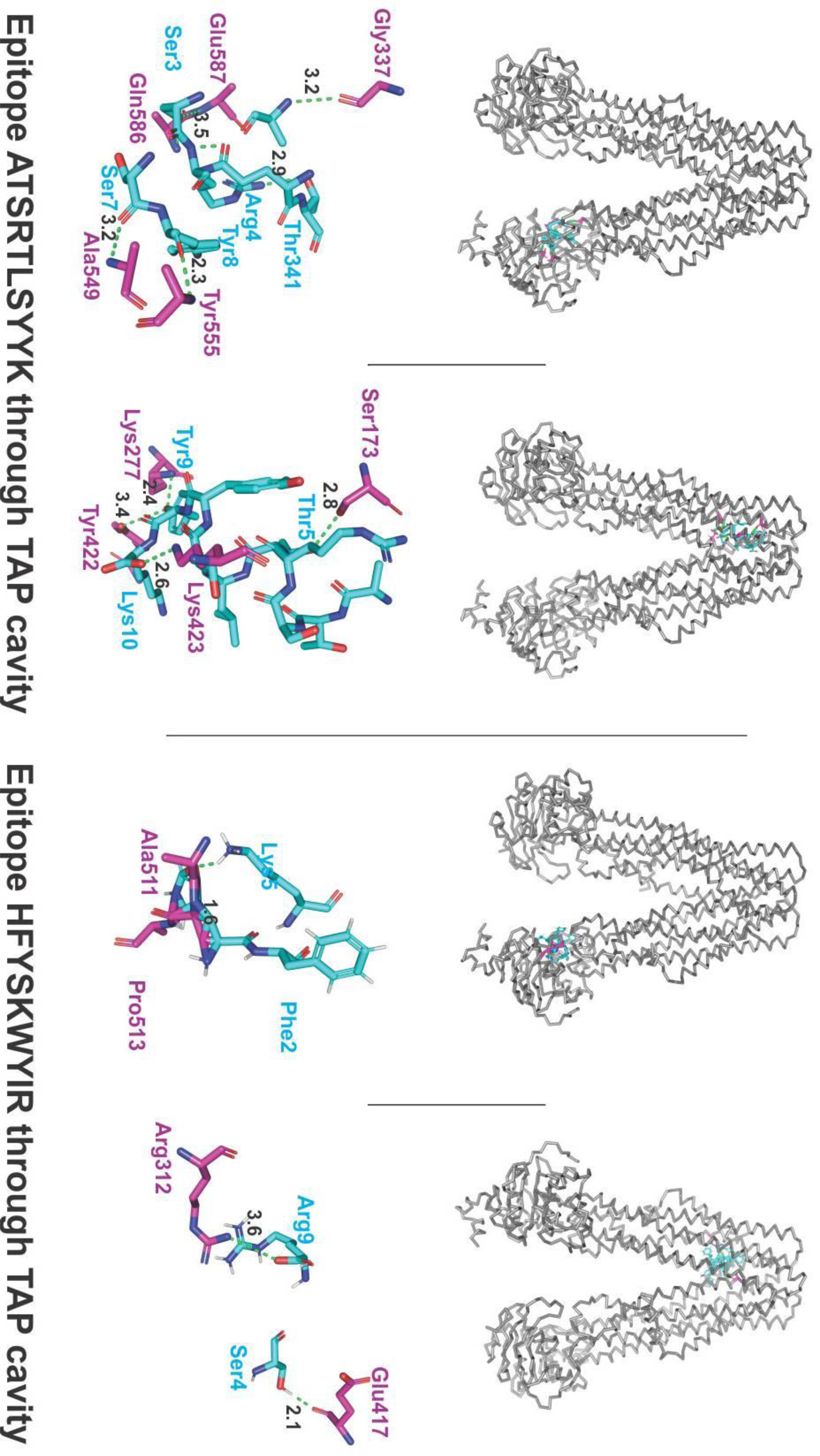
Molecular docking analysis of CTL epitopes within the TAP transporter cavity. Molecular interaction of CTL epitopes (cyan sticks) within the TAP cavity (gray ribbon/sticks) is shown. Detailed interaction between the residues of epitopes and TAP transporter residues have been shown with hydrogen formation shown with green dots. H bonds are shown in gereen dots with lengths in Angstroms.

**Figure 7.**
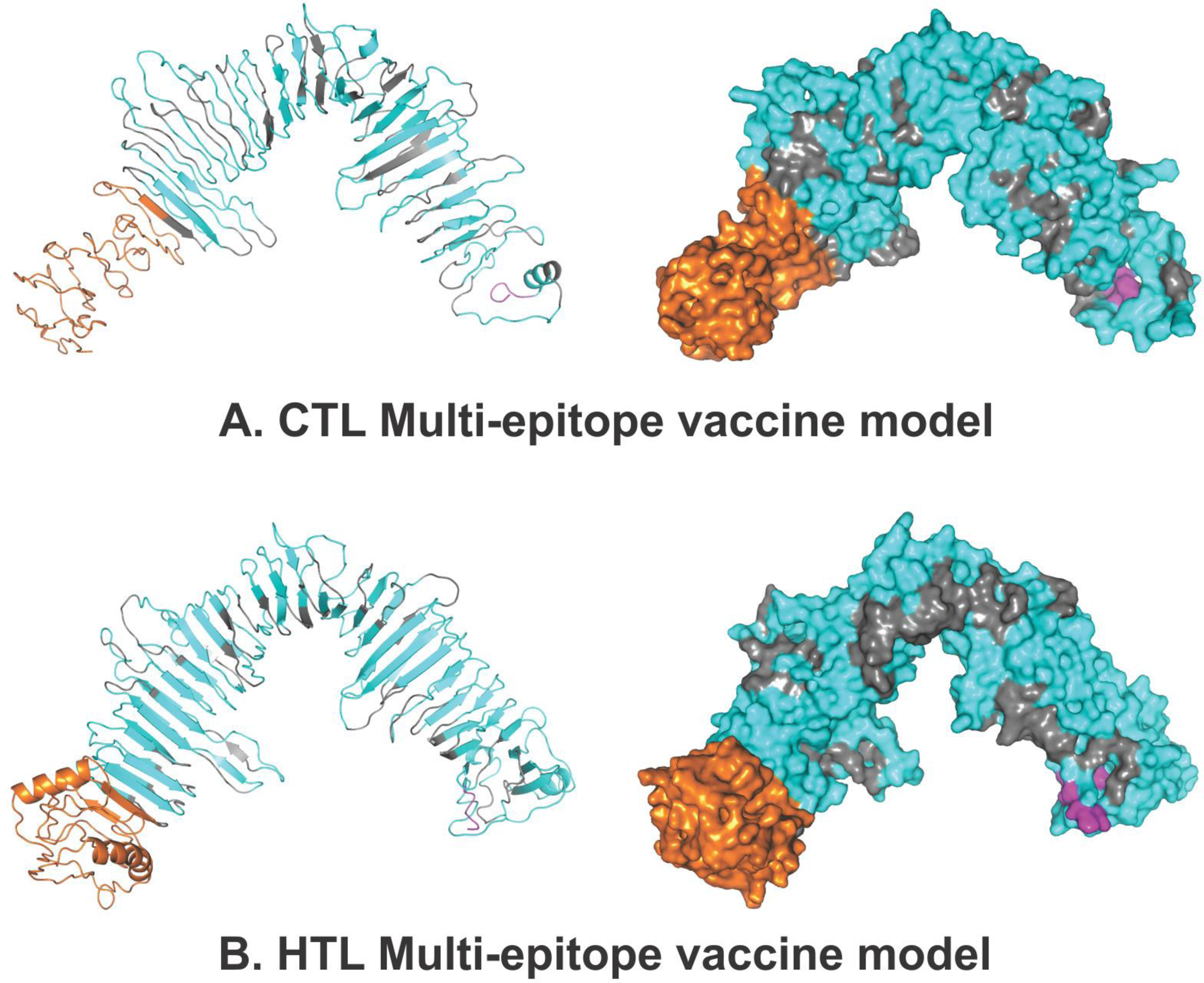
Tertiary structure modelling of CTL and HTL Multi-Epitope Vaccines. Tertiary structural models of CTL and HTL MEVs have been shown. The epitopes are shown in cyan. The adjuvant (Ov-ASP-1) is shown in orange. The linkers are shown in gray and 6xHis tag is shown in magenta. Both the cartoon and surface presentation of both the MEVs are shown.

##### Validation of CTL and HTL MEVs refined models

Both the CTL and HTL model were analyzed by the RAMPAGE analysis tool after refinement. The refined CTL MEV model was found to have 85.8% residues in favored region, 11.3% residues in allowed region, and only 3.0% residues in the outlier region; while the refined HTL MEV model was found to have 88.9% of residues in favored region, 8.9% residues in allowed region, and only 2.2% residues in the outlier region (Fig.8)

**Figure 8.**
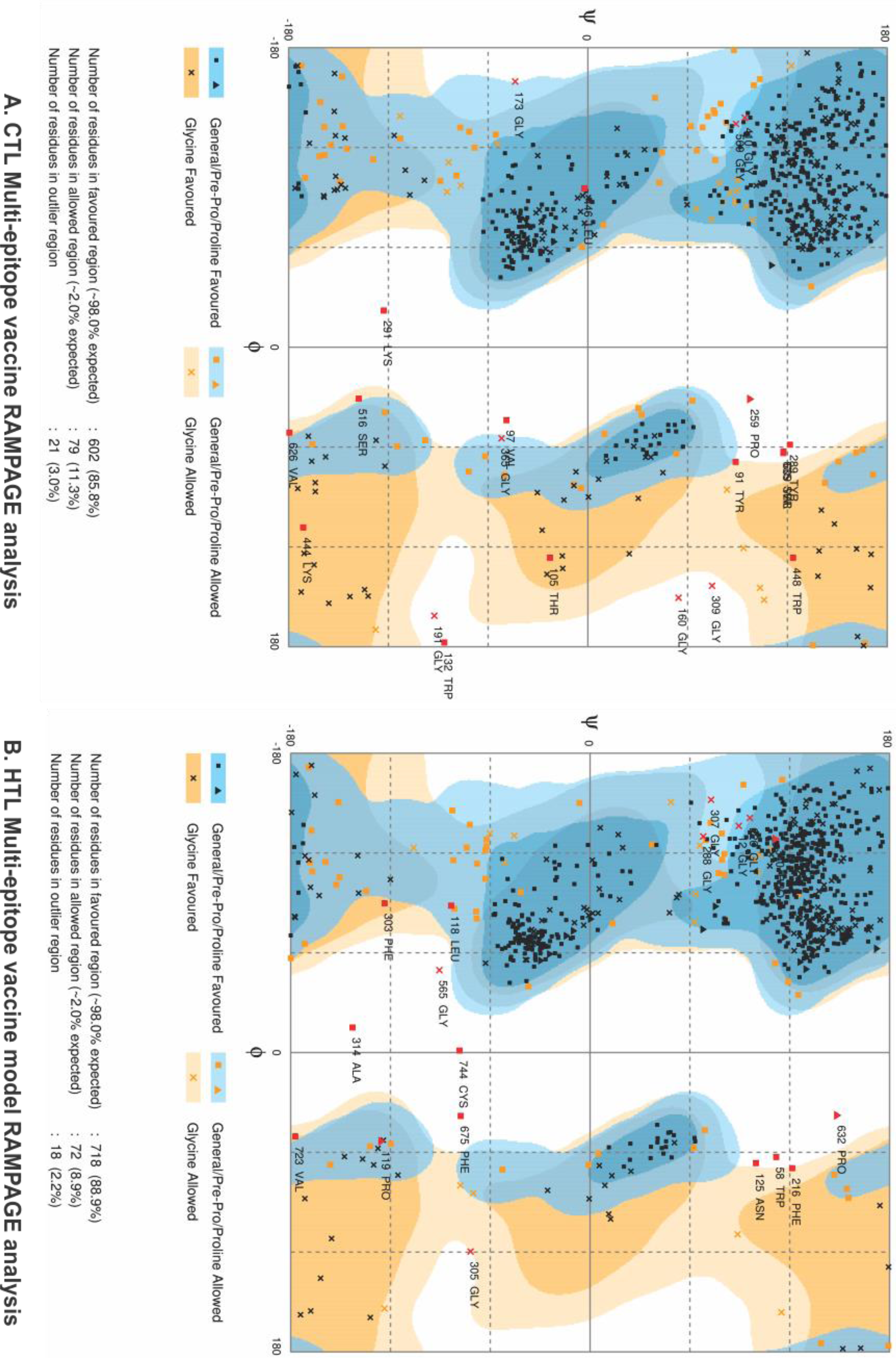
RAMPAGE analysis of CTL and HTL MEVs. The RAMPAGE analysis of both the CTL and HTL MEVs has been done and shown here.

##### Linear and Discontinuous B-cell epitope prediction from MEVs

The linear and discontinuous B-cell epitope prediction was performed to enlist potential linear and discontinuous epitopes from the refined 3D models of CTL and HTL MEVs utilizing the ElliPro tool available on IEDB server. The screening revealed that the CTL MEV carries 17 linear and 2 potential discontinuous B cell epitopes and that of HTL MEV carries 17 linear and 4 potential discontinuous epitopes. The high range of the PI (Protrusion Index) score of the linear and discontinuous epitopes from CTL and HTL MEVs show a high potential of the epitopes to cause humoral immune response (PI score: CTL MEV linear & discontinuous B cell epitopes −0.511 to 0.828 & 0.664 to 0.767 respectively; HTL MEV linear and discontinuous B cell epitopes −0.518 to 0.831 & 0.53 to 0.776 respectively) (Supplementary table S4,S5, S6, S7).

### Molecular interaction analysis of MEVs with immunological receptor

#### Molecular docking study of MEVs and TLR-3

Both the refined models of CTL and HTL MEVs were further studied for their molecular interaction with the ectodomain (ECD) of human TLR-3. Therefore, molecular docking of CTL and HTL MEVs model with the TLR-3 crystal structure model (PDB ID: 2A0Z) was performed utilizing the PatchDock tool. Generated docking conformation with the highest scores of 20776 and 20350 for CTL and HTL MEVs respectively were chosen for further study. The highest docking score indicates the best geometric shape complementarity fitting conformation of MEVs and the TLR-3 receptor as predicted by the PatchDock tool. Both the CTL and HTL MEVs were fitting into the ectodomain region of TLR-3 after docking (Fig.9A, 9C). The CTL and HTL MEVs have shown to form multiple hydrogen bonds within the ectodomain cavity region of TLR-3.

**Figure 9.**
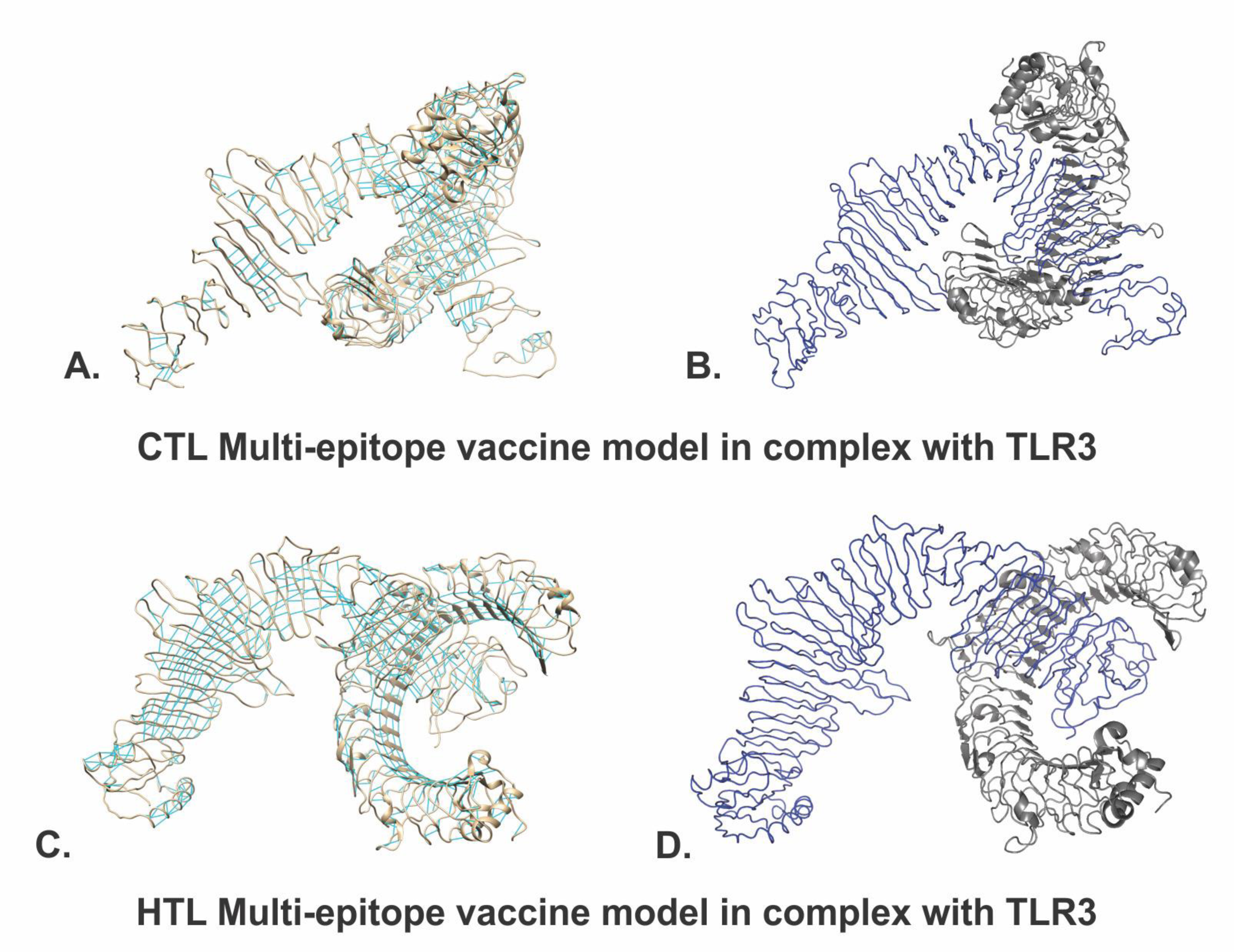
Molecular Docking study of CTL and HTL MEVs with TLR-3 receptor. (A), (C) The docking complex of CTL-TLR and HTL-TLR3 have been shown. The TLR3 is shown in the cartoon, and the MEVs are shown in the ribbon. Hydrogen bond formation is shown by cyan lines. (B), (D) B-factor presentation for the docked MEVs to the TLR3 receptor. The presentation is in VIBGYOR color, with blue showing low B-factor and red show high B-factor. Here most of the MEV regions are in blue showing low B-factor and hence indicate stable complex formation with TLR3 receptor. Images were generated by PyMol and UCSF Chimera (Pettersen et al., 2004).

The B-factor analysis of MEVs-TLR3 complexes was also performed. The B-factor indicates the displacement of the atomic positions from an average (mean) value i.e. the more flexible an atom is the larger the displacement from the mean position will be (mean-squares displacement) (Fig.9B, 9D). The B-factor analysis of the CTL and HTL MEVs bound to the TLR3 receptor shows that most of the regions of MEVs bound to TLR3 are stable nature in nature. The B-Factor analysis has been represented by the VIBGYOR color presentation with blue represents low B-factor and red represents high B-factor. Hence, the results suggest a stable complex formation tendency for both the CTL and HTL MEVs with the ectodomain of the human TLR-3 receptor (Fig.9B, 9D).

#### Molecular Dynamics (MD) Simulations study of MEVs and TLR-3 complex

Both the complexes of CTL MEV – TLR3 and the HTL MEV – TLR3 were further subjected for molecular dynamics simulation analysis to investigate the stability of the molecular interaction involved. Both the MEVs-TLR3 complexes have shown a very convincing and reasonably stable root mean square deviation (RMSD) values for Cα, Back bone, and all atom (CTL-MEV-TLR3 Complex: ∼4 to ∼7.5 Å; HTL-MEV-TLR3 Complex: ∼ 3.0 to ∼9.8 Å) which stabilizing towards the end, Fig 10. A & C. The RMSD of both complexes was maintained to the above mentioned RMSD range for a given time window of 10 ns at reasonably invariable temperature (∼278 K) and pressure (∼1 atm). Molecular docking and molecular dynamics simulation study of all the MEVs-TLRs complexes indicate a stable complex formation tendency. All most all the animo acid residues of the CTL and HTL MEVs in complexed with TLR3 have shown to have root mean square fluctuation (RMSF) in acceptable range (∼2 to ∼6 Å), Fig. 10 B & D. These results indicate both the CTL MEV – TLR3 and HTL MEV – TLR3 complexes to have a stable nature with acceptable molecular interaction tendency.

**Figure 10.**
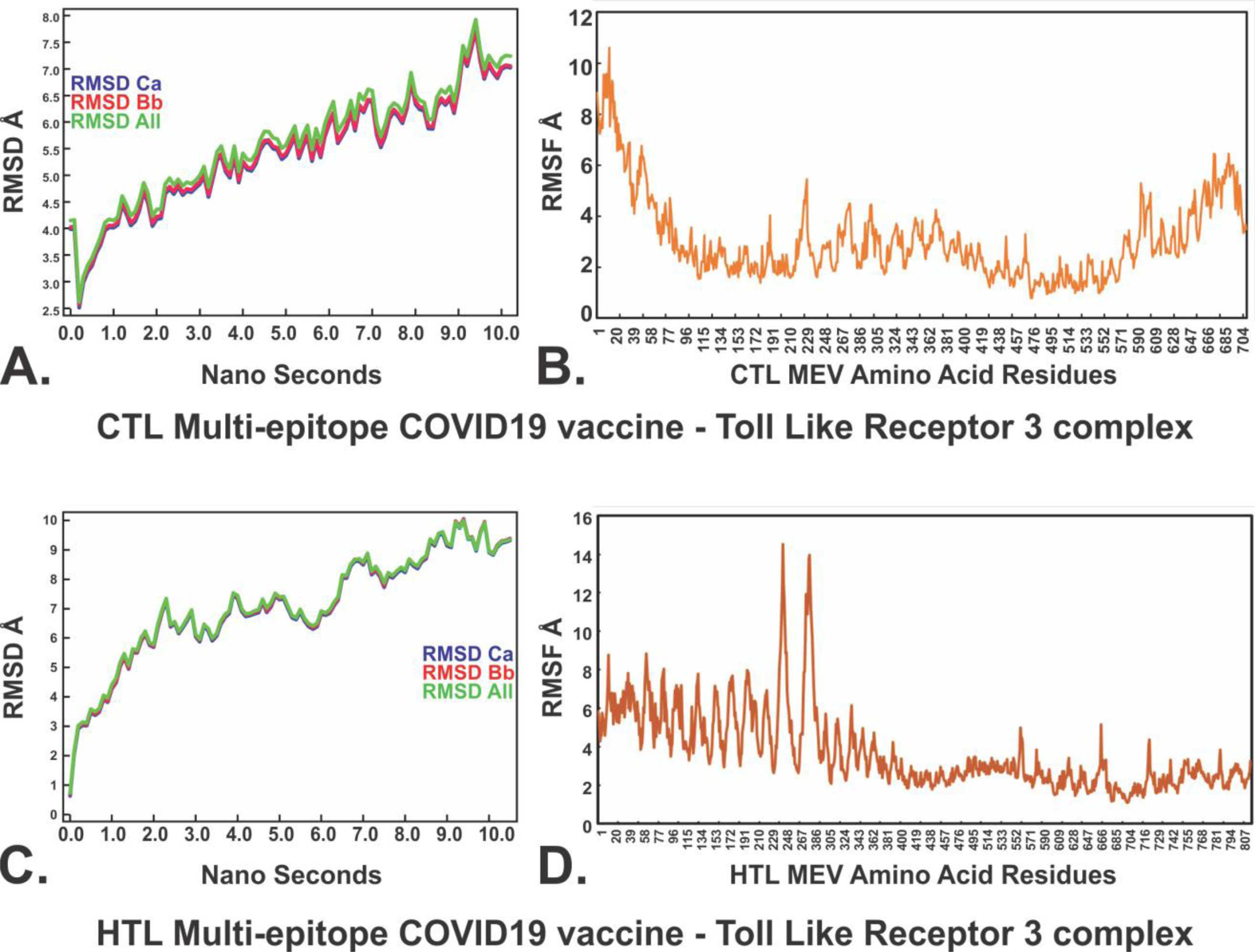
Molecular Dynamics simulation of CTL and HTL MEVs with TLR-3 receptor. (A), (C): Root Mean Square Deviation (RMSD) for Cα, Back bone and all atoms (RMSD Ca, RMSD Bb, & RMSD All) respectivally for The CTL MEV – TLR3 complex and The HTL MEV – TLR3 complex., docking complex of CTL-TLR and HTL-TLR3 have been shown. (B), (D): Root Mean Square Fluctuation (RMSF) for all the amino acid residues of CTL MEV and the HTL MEV in complex with TLR3 immune receptor.

### *In-silico* analysis of MEVs for cloning and expression potency

#### Analysis of cDNA of both the MEVs for cloning and expression in the mammalian host cell line

Complementary DNA optimized for CTL and HTL expression in the mammalian host cell line (Human) was generated by utilizing the Java Codon Adaptation Tool. Further, the generated optimized cDNA’s for both the MEVs were analyzed by utilizing the GenScript Rare Codon Analysis Tool. The analysis revealed that the codon-optimized cDNA of both the CTL and HTL MEVs bear all the crucial and favorable compositions for high-level expression in a mammalian cell line (CTL-MEV: GC content 70.40%, CAI (Codon Adaptation Index) score 1.00 and 0% tandem rare codons; HTL MEV: GC content 69.26%, CAI score 1.00 and 0% tandem rare codons). Ideally, the GC content of a cDNA should be 30% to 70%, CAI score that indicates the possibility of cDNA expression in a chosen expression system should be between 0.8-1.0, and the tandem rare codon frequency that indicates the presence of low-frequency codons in cDNA should be <30%. The tandem rare codons may hinder the proper expression of the cDNA or even interrupt the translational machinery of the chosen expression system. Hence as per the GenScript Rare Codon analysis, the cDNA of both the MEVs satisfies all the mentioned parameters and are predicted to have high expression in the mammalian host cell line (Human).

## CONCLUSION

In the present study, we have designed and proposed two multi-epitope vaccines derived from multiple CTL and HTL epitopes against SARS-CoV-2 (COVID19). The chosen CTL and HTL epitopes show significant sequence overlap with screened linear B cell epitopes. The shortlisted CTL and HTL epitopes were utilized to design CTL and HTL multi-epitope vaccine. Both the generated CTL and HTL multi-epitope vaccine tertiary models have shown to carry potential linear and discontinuous B cell epitopes as well as potential INF-γ epitopes. Hence the designed MEVs are predicted to have the potential to elicit humoral as well as cellular immune responses. Since Onchocerca volvulus activation-associated secreted protein-1 (Ov-ASP-1) binds to the APCs and trigger pro-inflammatory cytokine production via Toll-like receptor 3 (TLR3), the truncated (residues 10-153) Ov-ASP-1 has been utilized as an adjuvant at N terminal of both the CTL and HTL MEVs models. Chosen overlapping clustering epitopes were validated for their molecular interaction with their respective HLA allele binders by molecular docking studies. The molecular interaction of the chosen CTL epitopes within the TAP transporter cavity was also analyzed. Analysis of the average world population coverage by both the shortlisted CTL and HTL epitopes combined revealed coverage of 96.10% world population. The molecular interaction analysis of both the CTL and HTL MEVs with the immunoreceptor TLR3 has shown very convincing structural fitting of the MEVs into the ectodomain of TLR3 receptor cavity. This result was further confirmed by the molecular dynamics simulation studies of both the CTL MEV – TLR3 and HTL MEV – TLR3 complexes, indicating stable molecular complex formation tendency for both the MEVs in complex with TLR3. The cDNA for both the MEVs was generated considering codon-biasing for expression in the mammalian host cell line (Human). Both the cDNA has been optimized in respect of their GC content and zero tandem rare codons for the cDNA to have high expression possibility in the mammalian host cell line (Human). Hence for further studies, both the design of CTL and HTL MEVs could be cloned, expressed and tested for *in-vivo* validations and animal trials as potential vaccine candidates against SARS-CoV-2 infection.

**Supplementary table S1. Homology modeling for HLA alleles.** Tertiary structures of HLA alleles were modeled by homology modeling using SwissModel server. Templates were chosen with the highest sequence identity. Generated models with acceptable QMEAN values were chosen for further studies.

**Supplementary table S2. INF-γ epitopes from CTL and HTL MEVs.** INF-γ inducing (POSITIVE) epitopes from CTL and HTL MEVs were screened by using “Motif and SVM hybrid” (MERCI & SVM) approach.

**Supplementary table S3. Refinement models of CTL and HTL MEVs.** Both the CTL and HTL MEVs models were refined by GalaxyWEB server. After refinement, in particular the Rama favored residues increased significantly.

**Supplementary table S4. B Cell linear epitopes of CTL MEVs.** Linear B Cell epitopes predicted by ElliPro (IEDB) from CTL MEVs.

**Supplementary table S5.** B Cell discontinuous epitopes of CTL MEVs. Discontinuous B Cell epitopes predicted by ElliPro (IEDB) from CTL MEVs.

**Supplementary table S6. B Cell linear epitopes of HTL MEVs.** Linear B Cell epitopes predicted by ElliPro (IEDB) from HTL MEVs.

**Supplementary table S7.** B Cell discontinuous epitopes of HTL MEVs. Discontinuous B Cell epitopes predicted by ElliPro (IEDB) from HTL MEVs.

**Supplementary table S8. Shortlisted high scoring CTL epitopes (MHC-I Binding Predictions).** Selected high scoring CTL epitopes and their respective HLA alleles binders predicted by “MHC-I Binding Predictions” IEDB tool. *In-silico* analysis has shown all the selected epitopes to be non-toxic (Non-Toxin) as well as they show significant conservancy and high immunogenicity.

**Supplementary table S9. Shortlisted high scoring CTL epitopes (MHC-I Processing Predictions).** Selected high scoring CTL epitopes and their respective HLA alleles binders predicted by “MHC-I Processing Predictions” IEDB tool. The screening gives detailed and combined scoring “Total score” for Proteasomal cleavage/TAP transport/MHC class I combined. *In-silico* analysis have shown all the selected epitopes to be non-toxic (Non-Toxin) as well as they show significant conservancy with high immunogenicity.

**Supplementary table S10. Shortlisted high scoring HTL epitopes (MHC-II Binding Predictions).** Selected high “Percentile rank” HTL epitopes with their respective HLA class II alleles binders predicted by the “**MHC-II Binding Predictions**” tool of IEDB are listed. *In-silico* analysis has shown all the selected epitopes to be non-toxic (Non-Toxin) as well as they show significant conservancy.

**Supplementary table S11. Shortlisted B Cell epitopes (BepiPred Linear B Cell Prediction).** B cell linear epitopes with length of 4 to 20 amino acids, predicted by the “BepiPred Linear B Cell Prediction” IEDB tool, from eleven SARS-CoV-2 proteins, are listed here. *In-silico* analysis has shown all the selected epitopes to be non-toxic (Non-Toxin) as well as they show significant amino acid sequence conservancy.

## AUTHOR CONTRIBUTION

Protocol design: S.S., M.K.; Methodology performed by S.S., S.V., M.K., R.K.; Global Economic risk analysis: R.K.B.; Data analysis, scientific writing and revising the article: S.S., S.V., M.K., R.K., R.K.B., A.K.S., H.J.S., M.Kolbe, KCP.

## ADDITIONAL INFORMATION

Authors declare to have no competing interests.

## Abbreviations

(APCs): Antigen-presenting cell
(CAI): Codon Adaptation Index
(COVID19): Coronavirus ID 19
(CoV): Coronavirus
(CoV): Coverage
(Cryo-EM): Cryo-Electron Microscopy
(CTL): Cytotoxic T lymphocyte
(ECD): ectodomain
(E Protein): Envelope Protein
(ER): endoplasmic reticulum
(EBI): European Bioinformatics Institute
(N Protein): Nucleocapsid Protein
(ORF): Ope reading Frame
(GDT): global distance test
(GRAVY): Grand average of hydropathicity
(RMSD): root mean square deviation
(RMSF): root mean square fluctuation
(S Protein): Surface protein
(IC50): half maximal inhibitory concentration
(HTL): Helper T lymphocyte
(HLA): Human Leukocyte Antigen
(IEDB): Immune Epitope Database
(IPD): Immuno Polymorphism Database
(IFN-γ): Interferon Gama
(IMGT): International ImMunoGeneTics project
(MHC): major histocompatibility complex
(M Protein): Membrane Protein
(MERCI): Motif-EmeRging with ClassesIdentification
(MD simulation): Molecular dynamics simulation
(MEV): Multi-epitope Vaccine
(MSA): Multiple Sequence Alignment
(nsp): non-structural protein
(NCBI): National Center for Biotechnology Information
(pMHC): peptide-MHC
(SARS-CoV-2): Severe Acute Respiratory Syndrome Coronavirus 2
(SMM): Stabilization Matrix alignment method
(TM): Transmembrane
(PDB): Protein Data Bank
(QMEAN): Qualitative Model Energy ANalysis
(RMSD): root mean square deviation
(SVM): Support Vector Machine
(TLR3): Toll-Like Receptor 3
(TAP): Transporter associated with antigen processing
(uGDT): un-normalized global distance test
(YASARA): Yet Another Scientifc Artifcial Reality Application

