## Supplementary for "Structural basis to design multi-epitope vaccines against Novel Coronavirus 19 (COVID19) infection, the ongoing pandemic emergency: an in silico approach"

**Supplementary table S1. Homology modeling for HLA alleles.** Tertiary structures of HLA alleles were modeled by homology modeling using SwissModel server. Templates were chosen with the highest sequence identity. Generated models with acceptable QMEAN values were chosen for further studies.

| S.No | HLA Class I Allele | Template used for modeling | % sequence identity | QMEAN |
|---|---|---|---|---|
| 1 | A*31:01 | 6o9b.1.A | 94.29% | 0.73 |
| 2 | A*11:01 | 6o9b.1.A | 97.50% | 1.14 |
| S.No | HLA Class II Allele | Template used for modeling | % sequence identity | QMEAN |
| 3 | DRB3*01:01 | 4is6.1.B | 90.10% | (-)1.84 |
| 4 | DRB1*09:01 | 1bx2.1.B | 87.43% | (-)0.46 |

**Supplementary table S2. INF-γ epitopes from CTL and HTL MEVs.** INF-γ inducing (POSITIVE) epitopes from CTL and HTL MEVs were screened by using "Motif and SVM hybrid" (MERCI & SVM) approach.

| CLT Epitopes also predicted to be IFN-gamma epitopes | | | | | |
|---|---|---|---|---|---|
| S.No | Start-END | Sequence | Method | Result | Score |
| 1 | 67-82 | RWANQCIFGHSPRQQ | MERCI | POSITIVE | 2 |
| 2 | 68-83 | WANQCIFGHSPRQQR | MERCI | POSITIVE | 2 |
| 3 | 69-84 | ANQCIFGHSPRQQRE | MERCI | POSITIVE | 2 |
| 4 | 70-85 | NQCIFGHSPRQQREG | MERCI | POSITIVE | 2 |
| 5 | 71-86 | QCIFGHSPRQQREGV | MERCI | POSITIVE | 2 |
| 6 | 83-98 | EGVGENVYAYWSSVS | MERCI | POSITIVE | 1 |
| 7 | 84-99 | GVGENVYAYWSSVSV | MERCI | POSITIVE | 2 |
| 8 | 120-135 | LYENNPSNNMTWKVA | MERCI | POSITIVE | 1 |
| 9 | 121-136 | YENNPSNNMTWKVAG | MERCI | POSITIVE | 1 |
| 10 | 122-137 | ENNPSNNMTWKVAGQ | MERCI | POSITIVE | 2 |
| 11 | 413-428 | CTDDNALAYYGGGGS | MERCI | POSITIVE | 1 |
| 12 | 428-443 | TDDNALAYYGGGGSM | MERCI | POSITIVE | 1 |
| 13 | 429-444 | DDNALAYYGGGGSMY | MERCI | POSITIVE | 1 |
| 14 | 439-454 | GGGGSSIINNTVYTK | MERCI | POSITIVE | 1 |
| 15 | 440-455 | GGGSSIINNTVYTKG | MERCI | POSITIVE | 2 |
| 16 | 441-456 | GGSSIINNTVYTKGG | MERCI | POSITIVE | 3 |
| 17 | 442-457 | GSSIINNTVYTKGGG | MERCI | POSITIVE | 3 |
| 18 | 443-458 | SSIINNTVYTKGGGG | MERCI | POSITIVE | 3 |
| 19 | 444-459 | SIINNTVYTKGGGGS | MERCI | POSITIVE | 3 |
| 20 | 445-460 | IINNTVYTKGGGGSL | MERCI | POSITIVE | 3 |
| HLT Epitopes also predicted to be IFN-gamma epitopes | | | | | |
| S.No | Start-END | Sequence | Method | Result | Score |
| 1 | 67-82 | RWANQCIFGHSPRQQ | MERCI | POSITIVE | 2 |
| 2 | 68-83 | WANQCIFGHSPRQQR | MERCI | POSITIVE | 2 |
| 3 | 69-84 | ANQCIFGHSPRQQRE | MERCI | POSITIVE | 2 |
| 4 | 70-85 | NQCIFGHSPRQQREG | MERCI | POSITIVE | 2 |
| 5 | 71-86 | QCIFGHSPRQQREGV | MERCI | POSITIVE | 2 |
| 6 | 83-98 | EGVGENVYAYWSSVS | MERCI | POSITIVE | 1 |
| 7 | 84-99 | GVGENVYAYWSSVSV | MERCI | POSITIVE | 2 |
| 8 | 120-135 | LYENNPSNNMTWKVA | MERCI | POSITIVE | 1 |
| 9 | 121-136 | YENNPSNNMTWKVAG | MERCI | POSITIVE | 1 |
| 10 | 122-137 | ENNPSNNMTWKVAGQ | MERCI | POSITIVE | 2 |
| 11 | 585-600 | GGGGSFAWWTAFVTN | MERCI | POSITIVE | 1 |
| 12 | 586-601 | GGGSFAWWTAFVTNV | MERCI | POSITIVE | 1 |
| 13 | 587-602 | GGSFAWWTAFVTNVN | MERCI | POSITIVE | 1 |
| 14 | 588-603 | GSFAWWTAFVTNVNA | MERCI | POSITIVE | 1 |
| 15 | 589-604 | SFAWWTAFVTNVNAS | MERCI | POSITIVE | 1 |
| 16 | 560-575 | FAWWTAFVTNVNASS | MERCI | POSITIVE | 1 |
| 17 | 561-576 | AWWTAFVTNVNASSG | MERCI | POSITIVE | 1 |
| 18 | 806-821 | GGGSCTQHQPYVVDD | MERCI | POSITIVE | 1 |
| 19 | 807-822 | GGSCTQHQPYVVDDP | MERCI | POSITIVE | 1 |
| 20 | 808-823 | GSCTQHQPYVVDDPC | MERCI | POSITIVE | 1 |

**Supplementary table S3. Refinement models of CTL and HTL MEVs.** Both the CTL and HTL MEVs models were refined by GalaxyWEB server. After refinement, in particular the Rama favored residues increased significantly.

| Galaxy Refinement for CTL MEV | | | | | | |
|---|---|---|---|---|---|---|
| Model | GDT-HA | RMSD | Mol Probity | Clash score | Poor rotamers | Rama favored |
| Initial | 1.0000 | 0.000 | 3.749 | 100.5 | 4.5 | 75.8 |
| MODEL 1 | 0.9371 | 0.459 | 2.539 | 23.2 | 0.4 | 83.6 |
| Galaxy Refinement for HTL MEV | | | | | | |
| Initial | 1.0000 | 0.000 | 3.554 | 131.2 | 2.2 | 81.3 |
| MODEL 1 | 0.9552 | 0.402 | 2.537 | 27.9 | 0.7 | 87.7 |

**Supplementary table S4. B Cell linear epitopes of CTL MEVs.** Linear B Cell epitopes predicted by ElliPro (IEDB) from CTL MEVs.

| CTL Multi-Epitope vaccine model linear epitopes | | | | | |
|---|---|---|---|---|---|
| S.No. | Start | End | Peptide | Number of residues | Score |
| 1 | 1 | 97 | IVVAVTGYNCPGGKLTALERKKIVGQNNKYRSDLINGKLKNRNGTY MPRGKNMLELTWDCKLESSAQRWANQCIFGHSPRQQREGVGENVYAYWSSV | 97 | 0.828 |
| 2 | 604 | 704 | LRARSVSPKGGGGSFLAFLLFLVGGGGSHFYSKWYIRGGGGSWTAG AAAYYVGGGGSFPNITNLCPFGGGGSNYNYLYRLFRGGGGSNYLYRLFRHHHHHH | 101 | 0.812 |
| 3 | 400 | 415 | LNGSCGSVGGGGSCTD | 16 | 0.693 |
| 4 | 265 | 277 | SSEMVMCGGSLYG | 13 | 0.678 |
| 5 | 422 | 438 | GGGGSCTDDNALAYYGG | 17 | 0.662 |
| 6 | 584 | 596 | FGGGGSRARSVSP | 13 | 0.619 |
| 7 | 219 | 231 | GGGSKPRQKRTAT | 13 | 0.618 |
| 8 | 490 | 502 | WSMATYYGGGGSY | 13 | 0.618 |
| 9 | 294 | 304 | GGGSISNSWLM | 11 | 0.607 |
| 10 | 362 | 372 | FYGGGGSFLFV | 11 | 0.599 |
| 11 | 380 | 393 | GGGSRYFRLTLGVY | 14 | 0.599 |
| 12 | 320 | 327 | KGGGGSQE | 8 | 0.55 |
| 13 | 445 | 456 | GLPWNVVRGGGG | 12 | 0.548 |
| 14 | 99 | 109 | VEGLKKTAGTD | 11 | 0.547 |
| 15 | 342 | 349 | FNVPMEKG | 8 | 0.534 |
| 16 | 521 | 524 | ASFY | 4 | 0.528 |
| 17 | 464 | 477 | VYTKGGGGSLPVNV | 14 | 0.511 |

**Supplementary table S5. B Cell discontinuous epitopes of CTL MEVs.** Discontinuous B Cell epitopes predicted by ElliPro (IEDB) from CTL MEVs.

| CTL Multi-Epitope vaccine model discontinuous epitopes | | | |
|---|---|---|---|
| S.No. | Residues | Number of residues | Score |
| 1 | I1, V2, V3, A4, V5, T6, G7, Y8, N9, C10, P11, G12, G13, K14, L15, T16, A17, L18, E19, R20, K21, K22, I23, V24, G25, Q26, N27, N28, K29, Y30, R31, S32, D33, L34, I35, N36, G37, K38, L39, K40, N41, R42, N43, G44, T45, Y46, M47, P48, R49, G50, K51, N52, M53, L54, E55, L56, T57, W58, D59, C60, K61, L62, E63, S64, S65, A66, Q67, R68, W69, A70, N71, Q72, C73, I74, F75, G76, H77, S78, P79, R80, Q81, Q82, R83, E84, G85, V86, G87, E88, N89, V90, Y91, A92, Y93, W94, S95, S96, V97, S98, V99, E100, G101, L102, K103, K104, T105, G107, T108, D109, K112, W114, W115, N125, P126, S127, N128, N129, K133, V134, A135, G136 | 120 | 0.767 |
| 2 | G160, A187, R188, G189, G219, G220, G221, S222, K223, P224, R225, Q226, R228, T229, A230, T231, G246, G247, G248, G249, S250, G264, S265, S266, E267, M268, V269, M270, C271, G272, G273, S274, L275, Y276, G277, G294, G295, G296, S297, I298, S299, N300, S301, W302, L303, M304, K320, G321, G322, G323, G324, S325, Q326, E327, I328, T341, F342, N343, V344, P345, M346, E347, K348, G349, Y363, G364, G365, G366, G367, S368, F369, L370, F371, V372, A374, G380, G381, G382, S383, R384, Y385, F386, R387, L388, T389, L390, G391, V392, L400, N401, G402, S403, C404, G405, S406, V407, G408, G409, G410, G411, S412, C413, T414, D415, G422, G423, G424, G425, S426, C427, T428, D429, D430, N431, A432, L433, A434, Y435, Y436, G445, L446, P447, W448, N449, V450, V451, R452, G453, G454, G455, G456, G469, G470, G471, S472, L473, P474, V475, N476, V477, W490, M492, A493, T494, Y495, Y496, G497, G498, G499, G500, S501, Y502, A521, S522, F523, Y524, I546, P547, Y548, N549, S550, K564, V565, S566, I567, W568, F584, G585, G586, G587, G588, S589, R590, A591, R592, S593, V594, S595, P596, G598, G599, G600, G601, Q603, L604, R605, A606, R607, S608, V609, S610, P611, K612, G613, G614, G615, G616, S617, F618, L619, A620, F621, L622, L623, F624, L625, V626, G627, G628, G629, G630, S631, H632, F633, Y634, S635, K636, W637, Y638, I639, R640, G641, G642, G643, G644, S645, W646, T647, A648, G649, A650, A651, A652, Y653, Y654, V655, G656, G657, G658, G659, S660, F661, P662, N663, I664, T665, N666, L667, C668, P669, F670, G671, G672, G673, G674, S675, N676, Y677, N678, Y679, L680, Y681, R682, L683, F684, G686, G687, G688, G689, S690, N691, Y692, L693, Y694, R695, R698, H699, H700, H701, H702, H703, H704 | 282 | 0.664 |

**Supplementary table S6. B Cell linear epitopes of HTL MEVs.** Linear B Cell epitopes predicted by ElliPro (IEDB) from HTL MEVs.

| HTL Multi-Epitope vaccine model linear epitopes | | | | | |
|---|---|---|---|---|---|
| S.No. | Start | End | Peptide | Number of residues | Score |
| 1 | 691 | 810 | INVFAFPFTIYSLLGGGGSKTQSLLIVNNATNVVGGGGSLLIVNNATNVVIK VCGGGGSQSLLIVNNATNVVIKGGGGSSLLIVNNATNVVIKVGGGGSTQ SLLIVNNATNVVIHHHHHH | 120 | 0.831 |
| 2 | 1 | 138 | IVVAVTGYNCPGGKLTALERKKIVGQNNKYRSDLINGKLKNRNGTYMPR GKNMLELTWDCKLESSAQRWANQCIFGHSPRQQREGVGENVYAYWSS VSVEGLKKTAGTDAGKSWWSKLPKLYENNPSNNMTWKVAGQG | 138 | 0.777 |
| 3 | 452 | 468 | VSIWNLDYIINLIGGGG | 17 | 0.713 |
| 4 | 323 | 333 | FVGGGGSESPF | 11 | 0.69 |
| 5 | 474 | 489 | WNLDYIINLIIGGGGS | 16 | 0.685 |
| 6 | 434 | 444 | TVYSHLLLVAA | 11 | 0.679 |
| 7 | 543 | 551 | LYGGGGSCF | 9 | 0.651 |
| 8 | 416 | 424 | TIPIQASLP | 9 | 0.63 |
| 9 | 278 | 287 | SAFFGMSGGG | 10 | 0.608 |
| 10 | 351 | 360 | ILASFSASTS | 10 | 0.597 |
| 11 | 496 | 508 | WNLDYIINLGGGG | 13 | 0.586 |
| 12 | 399 | 404 | PAQYEL | 6 | 0.572 |
| 13 | 378 | 383 | SAPPAQ | 6 | 0.565 |
| 14 | 673 | 684 | FAFPFTIYSLLL | 12 | 0.556 |
| 15 | 515 | 527 | ALITLATCELGGG | 13 | 0.544 |
| 16 | 570 | 573 | LCFL | 4 | 0.54 |
| 17 | 655 | 660 | DDPCPI | 6 | 0.518 |

**Supplementary table S7. B Cell discontinuous epitopes of HTL MEVs.** Discontinuous B Cell epitopes predicted by ElliPro (IEDB) from HTL MEVs.

| HTL Multi-Epitope vaccine model discontinuous epitopes | | | |
|---|---|---|---|
| S. No. | Residues | Number of residues | Score |
| 1 | C639, P640, I641, D655, D656, P657, C658, P659, I660, H661, F662, F673, A674, F675, P676, F677, T678, I679, Y680, S681, L682, L683, L684, G687, G688, I691, N692, V693, F694, A695, F696, P697, F698, T699, I700, Y701, S702, L703, L704, G705, G706, G707, G708, S709, K710, T711, Q712, S713, L714, L715, I716, V717, N718, N719, A720, T721, N722, V723, V724, G725, G726, G727, G728, S729, L730, L731, I732, V733, N734, N735, A736, T737, N738, V739, V740, I741, K742, V743, C744, G745, G746, G747, G748, S749, Q750, S751, L752, L753, I754, V755, N756, N757, A758, T759, N760, V761, V762, I763, K764, G765, G766, G767, G768, S769, S770, L771, L772, I773, V774, N775, N776, A777, T778, N779, V780, V781, I782, K783, V784, G785, G786, G787, G788, S789, T790, Q791, S792, L793, L794, I795, V796, N797, N798, A799, T800, N801, V802, V803, I804, H805, H806, H807, H808, H809, H810 | 145 | 0.776 |
| 2 | I1, V2, V3, A4, V5, T6, G7, Y8, N9, C10, P11, G12, G13, K14, L15, T16, A17, L18, E19, R20, K21, K22, I23, V24, G25, Q26, N27, N28, K29, Y30, R31, S32, D33, L34, I35, N36, G37, K38, L39, K40, N41, R42, N43, G44, T45, Y46, M47, P48, R49, G50, K51, N52, M53, L54, E55, L56, T57, W58, D59, C60, K61, L62, E63, S64, S65, A66, Q67, R68, W69, A70, N71, Q72, C73, I74, F75, G76, H77, S78, P79, R80, Q81, Q82, R83, E84, G85, V86, G87, E88, N89, V90, Y91, A92, Y93, W94, S95, S96, V97, S98, V99, E100, G101, L102, K103, K104, T105, A106, G107, T108, D109, A110, G111, K112, S113, W114, W115, S116, K117, L118, P119, K120, L121, Y122, E123, N124, N125, P126, S127, N128, N129, M130, T131, W132, K133, V134, A135, G136, Q137, G138, E145, A146, A147, A148, L150, F152, A154, V156, L163, A164, G165, G168, S169 | 151 | 0.748 |
| 3 | S278, A279, F280, F281, G282, M283, S284, G285, G286, G287, F303, G304, G305, A322, F323, V324, G325, G326, G327, G328, S329, E330, S331, P332, F333, I350, I351, L352, A353, S354, F355, S356, A357, S358, T359, S360, S378, A379, P380, P381, A382, Q383, P399, A400, Q401, Y402, E403, L404, G405, G406, G407, T416, I417, P418, I419, Q420, A421, S422, L423, P424, T434, V435, Y436, S437, H438, L439, L440, L441, V442, A444, G446, V452, S453, I454, W455, N456, L457, D458, Y459, I460, I461, N462, L463, I464, G465, G466, G467, G468, W474, N475, L476, D477, Y478, I479, I480, N481, L482, I483, I484, G485, G486, G487, G488, S489, I495, W496, N497, L498, D499, Y500, I501, I502, N503, L504, G505, G506, G507, G508, A515, L516, I517, T518, A520, T521, C522, E523, L538, A539, L543, Y544, G545, G546, G547, G548, S549, C550, F551, L570, C571, F572, L573, F596, L597, L598, F599, P615, Y616, V617, V618, D619 | 150 | 0.609 |
| 4 | L524, G525, G526, G527, G528 | 5 | 0.53 |

**Supplementary table S8. Shortlisted high scoring CTL epitopes (MHC-I Binding Predictions).** Selected high scoring CTL epitopes and their respective HLA alleles binders predicted by “MHC-I Binding Predictions” IEDB tool. *In-silico* analysis has shown all the selected epitopes to be non-toxic (Non-Toxin) as well as they show significant conservancy and high immunogenicity.

| SARS-CoV-2 CTL epitopes (MHC-I Binding Predictions) | | | | | | | | | |
|---|---|---|---|---|---|---|---|---|---|
| S.No. | Proteins | Peptide | Allele | Length | Immunogenicity | Conservancy | Toxicity | Method used | Percentile Rank |
| 1 | E Protein | FLAFVVFLL | A*02:01 | 9 | 0.30188 | 99.59% (480/482) | Non-Toxin | Consensus (ann/comblib_sidney2008/smm) | 0.2 |
| 2 | E Protein | FLAFVVFLL | A*02:03 | 9 | 0.30188 | 99.59% (480/482) | Non-Toxin | Consensus (ann/smm) | 0.25 |
| 3 | E Protein | FLAFVVFLLV | A*02:01 | 10 | 0.30526 | 99.59% (480/482) | Non-Toxin | Consensus (ann/smm) | 0.15 |
| 4 | E Protein | FLAFVVFLLV | A*02:03 | 10 | 0.30526 | 99.59% (480/482) | Non-Toxin | Consensus (ann/smm) | 0.23 |
| 5 | E Protein | LAFVVFLLV | B*51:01 | 9 | 0.2141 | 99.59% (480/482) | Non-Toxin | Consensus (ann/comblib_sidney2008/smm) | 0.2 |
| 6 | E Protein | LVKPSFYVY | B*15:01 | 9 | -0.11106 | 99.59% (480/482) | Non-Toxin | Consensus (ann/comblib_sidney2008/smm) | 0.2 |
| 7 | E Protein | RVKNLNSSR | A*31:01 | 9 | -0.32968 | 99.59% (480/482) | Non-Toxin | Consensus (ann/smm) | 0.16 |
| 8 | E Protein | SLVKPSFYVY | B*15:01 | 10 | -0.2443 | 99.59% (480/482) | Non-Toxin | Consensus (ann/smm) | 0.28 |
| 9 | E Protein | SVLLFLAFV | A*02:06 | 9 | 0.19022 | 99.59% (480/482) | Non-Toxin | Consensus (ann/smm) | 0.33 |
| 10 | E Protein | TLAILTALR | A*68:01 | 9 | 0.1989 | 99.17% (478/482) | Non-Toxin | Consensus (ann/smm) | 0.21 |
| 11 | E Protein | VLLFLAFVV | A*02:01 | 9 | 0.26315 | 99.59% (480/482) | Non-Toxin | Consensus (ann/comblib_sidney2008/smm) | 0.3 |
| 12 | E Protein | VSLVKPSFY | A*30:02 | 9 | -0.25372 | 99.59% (480/482) | Non-Toxin | Consensus (ann/smm) | 0.33 |
| 13 | M Protein | FIASFRLFAR | A*68:01 | 10 | 0.12185 | 99.79% (476/477) | Non-Toxin | Consensus (ann/smm) | 0.12 |
| 14 | M Protein | LIFLWLLWPV | A*02:06 | 10 | 0.40176 | 98.32% (469/477) | Non-Toxin | Consensus (ann/smm) | 0.08 |
| 15 | M Protein | LSYFIASFR | A*68:01 | 9 | 0.21181 | 97.90% (467/477) | Non-Toxin | Consensus (ann/smm) | 0.11 |
| 16 | M Protein | MWLSYFIASF | A*23:01 | 10 | 0.00197 | 97.69% (466/477) | Non-Toxin | Consensus (ann/smm) | 0.11 |
| 17 | M Protein | MWLSYFIASFR | A*68:01 | 11 | 0.03554 | 97.69% (466/477) | Non-Toxin | ann | 0.09 |
| 18 | M Protein | MWLSYFIASFR | A*33:01 | 11 | 0.03554 | 97.69% (466/477) | Non-Toxin | ann | 0.1 |
| 19 | M Protein | QWNLVIGFLF | A*23:01 | 10 | 0.28076 | 99.16% (473/477) | Non-Toxin | Consensus (ann/smm) | 0.12 |
| 20 | M Protein | RFLYIIKLIF | A*23:01 | 10 | 0.11728 | 98.74% (471/477) | Non-Toxin | Consensus (ann/smm) | 0.12 |
| 21 | M Protein | RTRSMWSF | A*32:01 | 8 | -0.25178 | 99.58% (475/477) | Non-Toxin | ann | 0.1 |
| 22 | N Protein | AQFAPSASAF | B*15:01 | 10 | -0.17446 | 97.59% (486/498) | Non-Toxin | Consensus (ann/smm) | 0.16 |
| 23 | N Protein | AQFAPSASAFF | B*15:01 | 11 | -0.11074 | 97.59% (486/498) | Non-Toxin | ann | 0.12 |
| 24 | N Protein | FTALTQHGK | A*68:01 | 9 | -0.0226 | 99.40% (495/498) | Non-Toxin | Consensus (ann/smm) | 0.18 |
| 25 | N Protein | KMKDLSPR | A*31:01 | 8 | -0.22357 | 99.60% (496/498) | Non-Toxin | ann | 0.15 |
| 26 | N Protein | KTFPPTEPK | A*11:01 | 9 | 0.1306 | 96.99% (483/498) | Non-Toxin | Consensus (ann/smm) | 0.11 |
| 27 | N Protein | KTFPPTEPK | A*30:01 | 9 | 0.1306 | 96.99% (483/498) | Non-Toxin | Consensus (ann/comblib_sidney2008/smm) | 0.2 |
| 28 | N Protein | RQKRTATKAY | B*15:01 | 10 | -0.06462 | 97.79% (487/498) | Non-Toxin | Consensus (ann/smm) | 0.2 |
| 29 | N Protein | SSPDDQIGYY | A*01:01 | 10 | 0.07924 | 99.40% (495/498) | Non-Toxin | Consensus (ann/smm) | 0.2 |
| 30 | ORF-1ab | ASFYYVWKSY | A*30:02 | 10 | 0.00073 | 99.78% (455/456) | Non-Toxin | Consensus (ann/smm) | 0.07 |
| 31 | ORF-1ab | CLAYYFMRFRR | A*33:01 | 11 | 0.12614 | 100% (456/456) | Non-Toxin | ann | 0.07 |
| 32 | ORF-1ab | ETISLAGSY | A*26:01 | 9 | -0.1653 | 100% (456/456) | Non-Toxin | Consensus (ann/smm) | 0.1 |
| 33 | ORF-1ab | IPLMYKGLPW | B*53:01 | 10 | -0.37784 | 100% (456/456) | Non-Toxin | Consensus (ann/smm) | 0.07 |
| 34 | ORF-1ab | IPLMYKGLPW | B*53:01 | 10 | -0.37784 | 100% (456/456) | Non-Toxin | Consensus (ann/smm) | 0.07 |
| 35 | ORF-1ab | KMNYQVNGY | A*30:02 | 9 | -0.06542 | 100% (456/456) | Non-Toxin | Consensus (ann/smm) | 0.07 |

| | | | | | | | | | |
|---|---|---|---|---|---|---|---|---|---|
| 36 | ORF-1ab | KMNYQVNGY | A*30:02 | 9 | -0.06542 | 100% (456/456) | Non-Toxin | Consensus (ann/smm) | 0.07 |
| 37 | ORF-1ab | STNVTIATY | A*30:02 | 9 | 0.25822 | 99.34% (453/456) | Non-Toxin | Consensus (ann/smm) | 0.09 |
| 38 | ORF-1ab | YAYLRKHFSM | B*08:01 | 10 | -0.13937 | 100% (456/456) | Non-Toxin | Consensus (ann/smm) | 0.07 |
| 39 | ORF3a | FMRIFTIGTV | A*02:03 | 10 | 0.47908 | 99.37% (478/481) | Non-Toxin | Consensus (ann/smm) | 0.17 |
| 40 | ORF3a | FTIGTVTLK | A*68:01 | 9 | 0.18024 | 99.37% (478/481) | Non-Toxin | Consensus (ann/smm) | 0.16 |
| 41 | ORF3a | HVTFFIYNK | A*68:01 | 9 | 0.36278 | 99.79% (480/481) | Non-Toxin | Consensus (ann/smm) | 0.16 |
| 42 | ORF3a | IIMRLWLCWK | A*03:01 | 10 | 0.27346 | 96.67% (465/481) | Non-Toxin | Consensus (ann/smm) | 0.13 |
| 43 | ORF3a | VEHVTFFIY | B*44:03 | 9 | 0.3766 | 99.58% (479/481) | Non-Toxin | Consensus (ann/smm) | 0.18 |
| 44 | ORF3a | YFLQSINFVR | A*33:01 | 10 | -0.03483 | 96.88% (466/481) | Non-Toxin | Consensus (ann/smm) | 0.2 |
| 45 | ORF6 | AEILLIIM | B*40:01 | 8 | 0.25884 | 99.58% (479/481) | Non-Toxin | Consensus (ann/smm) | 0.41 |
| 46 | ORF6 | FQVTIAEIL | A*02:06 | 9 | 0.38115 | 99.17% (477/481) | Non-Toxin | Consensus (ann/smm) | 0.52 |
| 47 | ORF6 | ILLIIMRTFK | A*03:01 | 10 | 0.2388 | 99.58% (479/481) | Non-Toxin | Consensus (ann/smm) | 0.17 |
| 48 | ORF6 | IMRTFKVSI | A*32:01 | 9 | -0.09496 | 99.79% (480/481) | Non-Toxin | Consensus (ann/comblib_sidney2008/smm) | 0.44 |
| 49 | ORF6 | KVSIWNLDY | A*30:02 | 9 | 0.29343 | 99.58% (479/481) | Non-Toxin | Consensus (ann/smm) | 0.41 |
| 50 | ORF6 | LLIIMRTFK | A*03:01 | 9 | 0.156 | 99.58% (479/481) | Non-Toxin | Consensus (ann/smm) | 0.36 |
| 51 | ORF6 | LSKSLTENKY | A*30:02 | 10 | -0.24668 | 98.75% (475/481) | Non-Toxin | Consensus (ann/smm) | 0.48 |
| 52 | ORF6 | SIWNLDYII | A*32:01 | 9 | 0.15011 | 99.58% (479/481) | Non-Toxin | Consensus (ann/comblib_sidney2008/smm) | 0.2 |
| 53 | ORF6 | VTIAEILLI | B*58:01 | 9 | 0.28951 | 99.17% (477/481) | Non-Toxin | Consensus (ann/comblib_sidney2008/smm) | 0.34 |
| 54 | ORF7a | CVRGTTVLLK | A*03:01 | 10 | 0.14952 | 99.58% (478/480) | Non-Toxin | Consensus (ann/smm) | 0.17 |
| 55 | ORF7a | EVQELYSPI | A*26:01 | 9 | -0.09723 | 98.12% (471/480) | Non-Toxin | Consensus (ann/smm) | 0.21 |
| 56 | ORF7a | EVQELYSPIF | A*26:01 | 10 | -0.03858 | 98.12% (471/480) | Non-Toxin | Consensus (ann/smm) | 0.22 |
| 57 | ORF7a | FALTCFSTQF | B*53:01 | 10 | -0.09369 | 98.75% (474/480) | Non-Toxin | Consensus (ann/smm) | 0.22 |
| 58 | ORF7a | FLIVAAIVF | B*15:01 | 9 | 0.29611 | 98.96% (475/480) | Non-Toxin | Consensus (ann/comblib_sidney2008/smm) | 0.2 |
| 59 | ORF7a | FLIVAAIVFI | A*02:01 | 10 | 0.38946 | 98.33% (472/480) | Non-Toxin | Consensus (ann/smm) | 0.21 |
| 60 | ORF7a | ITLCFTLKRK | A*03:01 | 10 | -0.06825 | 97.92% (470/480) | Non-Toxin | Consensus (ann/smm) | 0.17 |
| 61 | ORF7a | QELYSPIFLI | B*44:02 | 10 | 0.03838 | 98.33% (472/480) | Non-Toxin | Consensus (ann/smm) | 0.22 |
| 62 | ORF7a | QELYSPIFLI | B*44:03 | 10 | 0.03838 | 98.33% (472/480) | Non-Toxin | Consensus (ann/smm) | 0.25 |
| 63 | ORF7a | RSVSPKLFI | B*58:01 | 9 | -0.30783 | 98.33% (472/480) | Non-Toxin | Consensus (ann/comblib_sidney2008/smm) | 0.2 |
| 64 | ORF7b | DFYLCFLAF | A*23:01 | 9 | 0.05884 | 98.31% (232/236) | Non-Toxin | Consensus (ann/smm) | 0.29 |
| 65 | ORF7b | FLAFLLFLVL | A*02:01 | 10 | 0.23386 | 98.31% (232/236) | Non-Toxin | Consensus (ann/smm) | 0.26 |
| 66 | ORF7b | FLLFLVLIML | A*02:01 | 10 | 0.14288 | 99.58% (235/236) | Non-Toxin | Consensus (ann/smm) | 0.3 |
| 67 | ORF7b | FYLCFLAFLL | A*23:01 | 10 | 0.1745 | 98.31% (232/236) | Non-Toxin | Consensus (ann/smm) | 0.13 |
| 68 | ORF7b | FYLCFLAFLL | A*24:02 | 10 | 0.1745 | 98.31% (232/236) | Non-Toxin | Consensus (ann/smm) | 0.28 |
| 69 | ORF7b | IELSLIDFY | B*44:03 | 9 | 0.03153 | 99.58% (235/236) | Non-Toxin | Consensus (ann/smm) | 0.19 |
| 70 | ORF7b | LVLIMLIIFW | B*53:01 | 10 | 0.25452 | 99.58% (235/236) | Non-Toxin | Consensus (ann/smm) | 0.32 |
| 71 | ORF7b | MIELSLIDFY | A*01:01 | 10 | 0.06184 | 99.58% (235/236) | Non-Toxin | Consensus (ann/smm) | 0.35 |
| 72 | ORF7b | MLIIFWFSL | A*02:01 | 9 | 0.50177 | 99.58% (235/236) | Non-Toxin | Consensus (ann/comblib_sidney2008/smm) | 0.2 |
| 73 | ORF7b | SLIDFYLCFL | A*02:01 | 10 | 0.18838 | 98.31% (232/236) | Non-Toxin | Consensus (ann/smm) | 0.14 |
| 74 | ORF7b | YLCFLAFLL | A*02:01 | 9 | 0.21865 | 98.31% (232/236) | Non-Toxin | Consensus (ann/comblib_sidney2008/smm) | 0.3 |
| 75 | ORF8 | CPIHFYSKW | B*53:01 | 9 | -0.07935 | 99.38% (477/480) | Non-Toxin | Consensus (ann/comblib_sidney2008/smm) | 0.2 |
| 76 | ORF8 | CPIHFYSKWY | B*53:01 | 10 | -0.02216 | 99.38% (477/480) | Non-Toxin | Consensus (ann/smm) | 0.2 |
| 77 | ORF8 | CSFYEDFLEY | A*01:01 | 10 | 0.31272 | 99.79% (479/480) | Non-Toxin | Consensus (ann/smm) | 0.24 |
| 78 | ORF8 | DFLEYHDVR | A*33:01 | 9 | 0.16684 | 99.58% (478/480) | Non-Toxin | Consensus (ann/smm) | 0.13 |
| 79 | ORF8 | FTINCQEPK | A*68:01 | 9 | -0.04683 | 100.00% (480/480) | Non-Toxin | Consensus (ann/smm) | 0.18 |
| 80 | ORF8 | GARKSAPLI | A*30:01 | 9 | -0.34031 | 99.17% (476/480) | Non-Toxin | Consensus (ann/comblib_sidney2008/smm) | 0.3 |

| | | | | | | | | | |
|---|---|---|---|---|---|---|---|---|---|
| 81 | ORF8 | GIITTVAAF | B*15:01 | 9 | 0.2148 | 99.17% (476/480) | Non-Toxin | Consensus (ann/comblib_sidney2008/smm) | 0.3 |
| 82 | ORF8 | GSLVVRCSFY | A*30:02 | 10 | 0.00657 | 100.00% (480/480) | Non-Toxin | Consensus (ann/smm) | 0.24 |
| 83 | ORF8 | IQYIDIGNY | A*30:02 | 9 | 0.30442 | 99.38% (477/480) | Non-Toxin | Consensus (ann/smm) | 0.12 |
| 84 | ORF8 | LGIITTVAAF | B*15:01 | 10 | 0.34746 | 99.17% (476/480) | Non-Toxin | Consensus (ann/smm) | 0.31 |
| 85 | ORF8 | LQSCTQHQPY | B*15:01 | 10 | -0.25674 | 96.04% (461/480) | Non-Toxin | Consensus (ann/smm) | 0.17 |
| 86 | ORF8 | NYTVSCLPF | A*23:01 | 9 | -0.17355 | 51.46% (247/480) | Non-Toxin | Consensus (ann/smm) | 0.23 |
| 87 | ORF8 | QSCTQHQPY | A*01:01 | 9 | -0.16503 | 96.04% (461/480) | Non-Toxin | Consensus (ann/smm) | 0.28 |
| 88 | ORF10 | FAFPFTIYSL | A*02:06 | 10 | 0.20414 | 99.79% (478/479) | Non-Toxin | Consensus (ann/smm) | 0.48 |
| 89 | ORF10 | FPFTIYSL | B*51:01 | 8 | 0.06356 | 100.00% (479/479) | Non-Toxin | Consensus (ann/smm) | 0.39 |
| 90 | ORF10 | FPFTIYSLL | B*51:01 | 9 | 0.05708 | 100.00% (479/479) | Non-Toxin | Consensus (ann/comblib_sidney2008/smm) | 0.3 |
| 91 | ORF10 | FPFTIYSLLL | B*53:01 | 10 | 0.03149 | 100.00% (479/479) | Non-Toxin | Consensus (ann/smm) | 0.13 |
| 92 | ORF10 | FPFTIYSLLL | B*51:01 | 10 | 0.03149 | 100.00% (479/479) | Non-Toxin | Consensus (ann/smm) | 0.14 |
| 93 | ORF10 | FTIYSLLLCR | A*68:01 | 10 | -0.18372 | 100.00% (479/479) | Non-Toxin | Consensus (ann/smm) | 0.12 |
| 94 | ORF10 | FTIYSLLLCR | A*33:01 | 10 | -0.18372 | 100.00% (479/479) | Non-Toxin | Consensus (ann/smm) | 0.35 |
| 95 | ORF10 | MGYINVFAFPF | A*23:01 | 11 | 0.40977 | 99.37% (476/479) | Non-Toxin | Consensus (ann/smm) | 0.41 |
| 96 | ORF10 | TIYSLLLCR | A*68:01 | 9 | -0.22977 | 100.00% (479/479) | Non-Toxin | Consensus (ann/smm) | 0.47 |
| 97 | ORF10 | YINVFAFPF | B*53:01 | 9 | 0.28259 | 99.37% (476/479) | Non-Toxin | Consensus (ann/comblib_sidney2008/smm) | 0.35 |
| 98 | ORF10 | YINVFAFPF | A*32:01 | 9 | 0.28259 | 99.37% (476/479) | Non-Toxin | Consensus (ann/comblib_sidney2008/smm) | 0.4 |
| 99 | S Protein | FVFLVLLPLV | A*02:06 | 10 | 0.02996 | 98.94% (467/472) | Non-Toxin | Consensus (ann/smm) | 0.1 |
| 100 | S Protein | GNYNYLYRLFR | A*33:01 | 11 | 0.08205 | 98.52% (465/472) | Non-Toxin | ann | 0.11 |
| 101 | S Protein | KSFTVEKGIY | A*30:02 | 10 | 0.11812 | 99.36% (469/472) | Non-Toxin | Consensus (ann/smm) | 0.11 |

**Supplementary table S9. Shortlisted high scoring CTL epitopes (MHC-I Processing Predictions).** Selected high scoring CTL epitopes and their respective HLA alleles binders predicted by "MHC-I Processing Predictions" IEDB tool. The screening gives detailed and combined scoring "Total score" for Proteasomal cleavage/TAP transport/MHC class I combined. *In-silico* analysis have shown all the selected epitopes to be non-toxic (Non-Toxin) as well as they show significant conservancy with high immunogenicity.

| SARS-CoV-2 CTL epitopes (MHC-I Processing Predictions) | | | | | | | | | | | | |
|---|---|---|---|---|---|---|---|---|---|---|---|---|
| S.No. | Proteins | Allele | Peptide Length | Peptide | Immunogenicity | Conservancy | Toxicity | Proteasome Score | TAP Score | MHC Score | Processing Score | Total Score | MHC IC50[nM] |
| 1 | E Protein | B*15:01 | 10 | SLVKPSFYVY | -0.2443 | 99.59% (480/482) | Non-Toxin | 1.51 | 1.36 | -1.39 | 2.86 | 1.47 | 24.6 |
| 2 | E Protein | B*15:01 | 9 | LVKPSFYVY | -0.11106 | 99.59% (480/482) | Non-Toxin | 1.51 | 1.35 | -1.42 | 2.86 | 1.44 | 26.3 |
| 3 | E Protein | A*30:02 | 9 | LVKPSFYVY | -0.11106 | 99.59% (480/482) | Non-Toxin | 1.51 | 1.35 | -1.61 | 2.86 | 1.25 | 40.4 |
| 4 | E Protein | A*02:01 | 9 | FLAFVVFLL | 0.30188 | 99.59% (480/482) | Non-Toxin | 1.45 | 0.41 | -0.81 | 1.86 | 1.05 | 6.5 |
| 5 | E Protein | B*15:01 | 9 | LTALRLCAY | 0.01886 | 99.17% (478/482) | Non-Toxin | 1.42 | 1.27 | -1.69 | 2.69 | 1 | 49.5 |
| 6 | M Protein | A*30:02 | 9 | ATSRTLSYY | -0.11604 | 98.95% (472/477) | Non-Toxin | 1.26 | 1.34 | -1.12 | 2.6 | 1.48 | 13.3 |
| 7 | M Protein | A*11:01 | 9 | ATSRTLSYY | -0.11604 | 98.95% (472/477) | Non-Toxin | 1.26 | 1.34 | -1.52 | 2.6 | 1.08 | 32.9 |
| 8 | M Protein | B*08:01 | 10 | FARTRSMWSF | -0.12986 | 99.58% (475/477) | Non-Toxin | 1.41 | 1.12 | -1.34 | 2.53 | 1.19 | 22.1 |
| 9 | M Protein | A*02:01 | 10 | FLWLLWPVTL | 0.31272 | 98.53% (470/477) | Non-Toxin | 1.85 | 0.46 | -1.16 | 2.31 | 1.16 | 14.3 |
| 10 | M Protein | A*68:01 | 9 | LSYFIASFR | 0.21181 | 97.90% (467/477) | Non-Toxin | 0.84 | 0.72 | -0.46 | 1.56 | 1.1 | 2.9 |
| 11 | M Protein | A*23:01 | 11 | LSYFIASFRLF | 0.2706 | 97.90% (467/477) | Non-Toxin | 1.25 | 1.18 | -1.36 | 2.43 | 1.07 | 23 |

| | | | | | | | | | | | | | |
|---|---|---|---|---|---|---|---|---|---|---|---|---|---|
| 12 | M Protein | A*23:01 | 10 | MWLSYFIASF | 0.00197 | 97.69% (466/477) | Non-Toxin | 1.38 | 1.26 | -0.91 | 2.63 | 1.73 | 8.1 |
| 13 | M Protein | A*24:02 | 10 | MWLSYFIASF | 0.00197 | 97.69% (466/477) | Non-Toxin | 1.38 | 1.26 | -1.27 | 2.63 | 1.36 | 18.7 |
| 14 | M Protein | B*15:01 | 10 | MWLSYFIASF | 0.00197 | 97.69% (466/477) | Non-Toxin | 1.38 | 1.26 | -1.33 | 2.63 | 1.3 | 21.6 |
| 15 | M Protein | A*23:01 | 10 | RFLYIIKLIF | 0.11728 | 98.74% (471/477) | Non-Toxin | 1.25 | 1.35 | -1.33 | 2.6 | 1.27 | 21.3 |
| 16 | M Protein | A*30:01 | 8 | RTRSMWSF | -0.25178 | 99.58% (475/477) | Non-Toxin | 1.41 | 1.28 | -1.51 | 2.69 | 1.18 | 32.7 |
| 17 | M Protein | A*30:02 | 9 | SGFAAYSRY | 0.00261 | 98.32% (469/477) | Non-Toxin | 1.53 | 1.17 | -1.4 | 2.69 | 1.3 | 25 |
| 18 | M Protein | B*15:01 | 10 | SQRVAGDSGF | 0.0305 | 98.53% (470/477) | Non-Toxin | 1.33 | 1.24 | -1.35 | 2.58 | 1.23 | 22.4 |
| 19 | M Protein | A*23:01 | 9 | SYFIASFRL | 0.18333 | 98.74% (471/477) | Non-Toxin | 1.45 | 0.62 | -0.99 | 2.08 | 1.09 | 9.8 |
| 20 | M Protein | B*35:01 | 9 | VATSRTLSY | -0.17295 | 98.95% (472/477) | Non-Toxin | 1.34 | 1.31 | -1.38 | 2.65 | 1.27 | 23.9 |
| 21 | M Protein | A*30:02 | 9 | YSRYRIGNY | 0.21358 | 98.53% (470/477) | Non-Toxin | 1.36 | 1.37 | -1.61 | 2.73 | 1.12 | 40.6 |
| 22 | N Protein | B*15:01 | 10 | AQFAPSASAF | -0.17446 | 97.59% (486/498) | Non-Toxin | 1.23 | 1.25 | -0.7 | 2.48 | 1.78 | 5 |
| 23 | N Protein | B*15:01 | 11 | AQFAPSASAFF | -0.11074 | 97.59% (486/498) | Non-Toxin | 1.3 | 1.25 | -1.46 | 2.55 | 1.08 | 29 |
| 24 | N Protein | A*30:02 | 10 | KDLSPRWYFY | 0.14332 | 99.60% (496/498) | Non-Toxin | 1.58 | 1.21 | -1.68 | 2.79 | 1.1 | 48.3 |
| 25 | N Protein | A*30:02 | 10 | KMKDLSPRWY | -0.05692 | 99.60% (496/498) | Non-Toxin | 1.31 | 1.35 | -1.56 | 2.66 | 1.1 | 36.2 |
| 26 | N Protein | A*68:02 | 9 | NTASWFTAL | 0.22775 | 99.60% (496/498) | Non-Toxin | 1.42 | 0.49 | -0.79 | 1.91 | 1.12 | 6.2 |
| 27 | N Protein | B*15:01 | 10 | RQKRTATKAY | -0.06462 | 97.79% (487/498) | Non-Toxin | 1.51 | 1.34 | -1.55 | 2.85 | 1.3 | 35.5 |
| 28 | N Protein | A*01:01 | 10 | SSPDDQIGYY | 0.07924 | 99.40% (495/498) | Non-Toxin | 1.24 | 1.36 | -1.16 | 2.6 | 1.45 | 14.3 |
| 29 | ORF1ab | A*01:01 | 10 | ACTDDNALAY | 0.10055 | 99.78% (455/456) | Non-Toxin | 1.49 | 1.33 | -0.82 | 2.82 | 2 | 6.6 |
| 30 | ORF1ab | A*01:01 | 9 | CTDDNALAY | 0.07355 | 99.78% (455/456) | Non-Toxin | 1.49 | 1.23 | -0.45 | 2.72 | 2.27 | 2.8 |
| 31 | ORF1ab | A*26:01 | 9 | ETISLAGSY | -0.1653 | 100% (456/456) | Non-Toxin | 1.4 | 1.21 | -0.53 | 2.61 | 2.08 | 3.4 |
| 32 | ORF1ab | B*35:01 | 9 | FAVDAAKAY | -0.04849 | 98.90% (451/456) | Non-Toxin | 1.73 | 1.35 | -0.46 | 3.08 | 2.62 | 2.9 |
| 33 | ORF1ab | B*15:01 | 9 | ILMTARTVY | 0.12576 | 100% (456/456) | Non-Toxin | 1.65 | 1.34 | -1.1 | 2.99 | 1.89 | 12.5 |
| 34 | ORF1ab | B*15:01 | 10 | LILMTARTVY | 0.0012 | 100% (456/456) | Non-Toxin | 1.65 | 1.36 | -1.23 | 3.01 | 1.78 | 16.9 |
| 35 | ORF1ab | B*15:01 | 10 | LMSNLGMPSY | -0.30933 | 99.56% (454/456) | Non-Toxin | 1.55 | 1.34 | -1 | 2.9 | 1.89 | 10.1 |
| 36 | ORF1ab | B*35:01 | 9 | LPSLATVAY | 0.06748 | 98.02% (447/456) | Non-Toxin | 1.47 | 1.15 | -0.34 | 2.62 | 2.28 | 2.2 |
| 37 | ORF1ab | A*32:01 | 9 | RMYIFFASF | 0.29328 | 99.12% (452/456) | Non-Toxin | 1.32 | 1.36 | -0.82 | 2.68 | 1.86 | 6.6 |
| 38 | ORF1ab | A*30:02 | 10 | RMYIFFASFY | 0.32633 | 99.12% (452/456) | Non-Toxin | 1.28 | 1.53 | -0.97 | 2.81 | 1.83 | 9.4 |
| 39 | ORF1ab | A*03:01 | 10 | RMYIFFASFY | 0.32633 | 99.12% (452/456) | Non-Toxin | 1.28 | 1.53 | -1 | 2.81 | 1.8 | 10.1 |
| 40 | ORF1ab | B*44:03 | 9 | SEFSSLPSY | -0.40603 | 99.34% (453/456) | Non-Toxin | 1.49 | 1.35 | -1.03 | 2.84 | 1.82 | 10.6 |
| 41 | ORF1ab | B*15:01 | 9 | VMYMGTLSY | -0.21438 | 97.58% (445/456) | Non-Toxin | 1.3 | 1.5 | -0.8 | 2.79 | 1.99 | 6.3 |
| 42 | ORF1ab | A*03:01 | 9 | VMYMGTLSY | -0.21438 | 97.58% (445/456) | Non-Toxin | 1.3 | 1.5 | -0.98 | 2.79 | 1.82 | 9.5 |
| 43 | ORF3a | B*44:03 | 11 | AGLEAPFLYLY | 0.21841 | 96.67% (465/481) | Non-Toxin | 1.51 | 1.29 | -1.63 | 2.8 | 1.17 | 42.9 |
| 44 | ORF3a | A*23:01 | 10 | LVYFLQSINF | -0.05419 | 96.88% (466/481) | Non-Toxin | 1.3 | 1.31 | -1.39 | 2.61 | 1.23 | 24.3 |
| 45 | ORF6 | A*30:02 | 9 | KVSIWNLDY | 0.29343 | 99.58% (479/481) | Non-Toxin | 1.21 | 1.33 | -1.46 | 2.54 | 1.08 | 28.8 |
| 46 | ORF7a | B*35:01 | 9 | CPDGVKHVY | -0.07008 | 98.54% (473/480) | Non-Toxin | 1.43 | 1.11 | -1.34 | 2.54 | 1.2 | 22 |
| 47 | ORF7b | A*02:01 | 9 | IIFWFSLEL | 0.2683 | 99.58% (235/236) | Non-Toxin | 1.65 | 0.55 | -1.2 | 2.21 | 1.01 | 15.7 |
| 48 | ORF7b | A*02:01 | 10 | SLIDFYLCFL | 0.18838 | 98.31% (232/236) | Non-Toxin | 1.7 | 0.5 | -0.74 | 2.2 | 1.46 | 5.5 |
| 49 | ORF7b | A*02:03 | 10 | SLIDFYLCFL | 0.18838 | 98.31% (232/236) | Non-Toxin | 1.7 | 0.5 | -0.79 | 2.2 | 1.41 | 6.2 |
| 50 | ORF7b | A*02:06 | 10 | SLIDFYLCFL | 0.18838 | 98.31% (232/236) | Non-Toxin | 1.7 | 0.5 | -0.93 | 2.2 | 1.27 | 8.6 |
| 51 | ORF8 | B*15:01 | 9 | GIITTVAAF | 0.2148 | 99.17% (476/480) | Non-Toxin | 1.27 | 1.13 | -1.26 | 2.4 | 1.14 | 18.1 |
| 52 | ORF8 | A*33:01 | 9 | HFYSKWYIR | -0.09452 | 99.38% (477/480) | Non-Toxin | 1.08 | 0.76 | -0.68 | 1.84 | 1.16 | 4.8 |
| 53 | ORF8 | A*31:01 | 9 | HFYSKWYIR | -0.09452 | 99.38% (477/480) | Non-Toxin | 1.08 | 0.76 | -0.79 | 1.84 | 1.05 | 6.2 |
| 54 | ORF8 | B*15:01 | 10 | LGIITTVAAF | 0.34746 | 99.17% (476/480) | Non-Toxin | 1.27 | 0.97 | -1.1 | 2.24 | 1.14 | 12.7 |
| 55 | ORF10 | A*68:01 | 10 | NVFAFPFTIY | 0.40129 | 99.58% (477/479) | Non-Toxin | 1.47 | 1.4 | -1.84 | 2.86 | 1.02 | 69.7 |

| | | | | | | | | | | | | |
|---|---|---|---|---|---|---|---|---|---|---|---|---|
| 56 | S Protein | B*35:01 | 9 | CVADYSVLY | -0.09595 | 100.00% (472/472) | Non-Toxin | 1.51 | 1.38 | -1.39 | 2.89 | 1.5 | 24.4 |
| 57 | S Protein | B*35:01 | 9 | FAMQMAYRF | -0.28061 | 100.00% (472/472) | Non-Toxin | 1.45 | 1.05 | -0.8 | 2.5 | 1.7 | 6.3 |
| 58 | S Protein | B*58:01 | 10 | KRSFIEDLLF | 0.29624 | 99.36% (469/472) | Non-Toxin | 1.23 | 1.26 | -1.11 | 2.49 | 1.37 | 12.9 |
| 59 | S Protein | A*23:01 | 10 | KWPWYIWLGF | 0.56424 | 99.58% (470/472) | Non-Toxin | 1.26 | 1.25 | -1.04 | 2.51 | 1.47 | 11 |
| 60 | S Protein | B*35:01 | 11 | LQIPFAMQMAY | -0.22124 | 100.00% (472/472) | Non-Toxin | 1.42 | 1.4 | -1.25 | 2.82 | 1.57 | 17.8 |
| 61 | S Protein | A*01:01 | 9 | LTDEMIAQY | 0.02757 | 99.58% (470/472) | Non-Toxin | 1.21 | 1.21 | -0.72 | 2.42 | 1.71 | 5.2 |
| 62 | S Protein | B*35:01 | 10 | QIPFAMQMAY | -0.25308 | 100.00% (472/472) | Non-Toxin | 1.42 | 1.35 | -1.16 | 2.77 | 1.61 | 14.5 |
| 63 | S Protein | A*30:02 | 9 | RISNCVADY | -0.02787 | 100.00% (472/472) | Non-Toxin | 1.16 | 1.47 | -1.26 | 2.63 | 1.37 | 18.2 |
| 64 | S Protein | B*58:01 | 9 | RSFIEDLLF | 0.27446 | 99.58% (470/472) | Non-Toxin | 1.23 | 1.32 | -0.72 | 2.54 | 1.82 | 5.3 |
| 65 | S Protein | B*15:01 | 10 | RVYSTGSNVF | -0.23394 | 100.00% (472/472) | Non-Toxin | 1.51 | 1.32 | -1.02 | 2.83 | 1.81 | 10.5 |
| 66 | S Protein | B*35:01 | 9 | SANNCTFEY | 0.13273 | 98.94% (467/472) | Non-Toxin | 1.18 | 1.3 | -1.11 | 2.48 | 1.37 | 12.8 |
| 67 | S Protein | A*30:02 | 10 | YTNSFTRGVY | 0.08467 | 99.58% (470/472) | Non-Toxin | 1.36 | 1.28 | -1.26 | 2.64 | 1.38 | 18.2 |

**Supplementary table S10. Shortlisted high scoring HTL epitopes (MHC-II Binding Predictions).** Selected high “Percentile rank” HTL epitopes with their respective HLA class II alleles binders predicted by the “**MHC-II Binding Predictions**” tool of IEDB are listed. *In-silico* analysis has shown all the selected epitopes to be non-toxic (Non-Toxin) as well as they show significant conservancy.

| SARS-CoV-2 HTL epitopes (MHC-II Binding Predictions) | | | | | | | |
|---|---|---|---|---|---|---|---|
| S.No | Proteins | Allele | Peptide | Conservancy | Toxicity | Method used | Percentile Rank |
| 1 | E Protein | DPA1*03:01/DPB1*04:02 | FLAFVVFLLVTLAIL | 99.59% (480/482) | Non-Toxin | Consensus (comb.lib./smm/nn) | 0.06 |
| 2 | E Protein | DPA1*03:01/DPB1*04:02 | LFLAFVVFLLVTLAI | 99.59% (480/482) | Non-Toxin | Consensus (comb.lib./smm/nn) | 0.03 |
| 3 | E Protein | DPA1*01:03/DPB1*02:01 | LFLAFVVFLLVTLAI | 99.59% (480/482) | Non-Toxin | Consensus (comb.lib./smm/nn) | 0.04 |
| 4 | E Protein | DPA1*01/DPB1*04:01 | LFLAFVVFLLVTLAI | 99.59% (480/482) | Non-Toxin | Consensus (comb.lib./smm) | 0.08 |
| 5 | E Protein | DPA1*02:01/DPB1*01:01 | LFLAFVVFLLVTLAI | 99.59% (480/482) | Non-Toxin | Consensus (comb.lib./smm/nn) | 0.1 |
| 6 | E Protein | DPA1*01:03/DPB1*02:01 | NSVLLFLAFVVFLLV | 99.59% (480/482) | Non-Toxin | Consensus (comb.lib./smm/nn) | 0.04 |
| 7 | E Protein | DPA1*03:01/DPB1*04:02 | NSVLLFLAFVVFLLV | 99.59% (480/482) | Non-Toxin | Consensus (comb.lib./smm/nn) | 0.06 |
| 8 | E Protein | DPA1*01/DPB1*04:01 | NSVLLFLAFVVFLLV | 99.59% (480/482) | Non-Toxin | Consensus (comb.lib./smm) | 0.07 |
| 9 | E Protein | DPA1*02:01/DPB1*01:01 | NSVLLFLAFVVFLLV | 99.59% (480/482) | Non-Toxin | Consensus (comb.lib./smm/nn) | 0.1 |
| 10 | E Protein | DPA1*02:01/DPB1*01:01 | VNSVLLFLAFVVFLL | 99.59% (480/482) | Non-Toxin | Consensus (comb.lib./smm/nn) | 0.1 |
| 11 | M Protein | DQA1*01:01/DQB1*05:01 | IKLIFLWLLWPVTLA | 98.11% (468/477) | Non-Toxin | Consensus (comb.lib./smm/nn) | 0.07 |
| 12 | M Protein | DQA1*01:01/DQB1*05:01 | KLIFLWLLWPVTLAC | 98.11% (468/477) | Non-Toxin | Consensus (comb.lib./smm/nn) | 0.11 |
| 13 | M Protein | DRB1*09:01 | RTLSYYKLGASQRVA | 99.16% (473/477) | Non-Toxin | Consensus (comb.lib./smm/nn) | 0.06 |
| 14 | M Protein | DRB1*01:01 | RTLSYYKLGASQRVA | 99.16% (473/477) | Non-Toxin | Consensus (comb.lib./smm/nn) | 0.67 |
| 15 | M Protein | DRB1*09:01 | SRTLSYYKLGASQRV | 99.16% (473/477) | Non-Toxin | Consensus (comb.lib./smm/nn) | 0.07 |
| 16 | M Protein | DRB1*09:01 | SYYKLGASQRVAGDS | 98.74% (471/477) | Non-Toxin | Consensus (comb.lib./smm/nn) | 0.48 |
| 17 | M Protein | DRB1*09:01 | TLSYYKLGASQRVAG | 98.95% (472/477) | Non-Toxin | Consensus (comb.lib./smm/nn) | 0.07 |
| 18 | M Protein | DRB1*01:01 | TLSYYKLGASQRVAG | 98.95% (472/477) | Non-Toxin | Consensus (comb.lib./smm/nn) | 0.67 |
| 19 | M Protein | DPA1*01:03/DPB1*02:01 | VGLMWLSYFIASFRL | 97.48% (465/477) | Non-Toxin | Consensus (comb.lib./smm/nn) | 0.07 |
| 20 | N Protein | DQA1*01:02/DQB1*06:02 | ANNAAIVLQLPQGTT | 97.59% (486/498) | Non-Toxin | Consensus (comb.lib./smm/nn) | 0.2 |
| 21 | N Protein | DRB1*11:01 | DQIGYYRRATRRIRG | 99.40% (495/498) | Non-Toxin | Consensus (smm/nn/sturniolo) | 0.42 |
| 22 | N Protein | DRB5*01:01 | DQIGYYRRATRRIRG | 99.40% (495/498) | Non-Toxin | Consensus (smm/nn/sturniolo) | 0.58 |

| | | | | | | | |
|---|---|---|---|---|---|---|---|
| 23 | N Protein | DQA1*01:02/DQB1*06:02 | GTRNPANNAAIVLQL | 97.59% (486/498) | Non-Toxin | Consensus (comb.lib./smm/nn) | 0.05 |
| 24 | N Protein | DRB1*07:01 | GTWLTYTGAIKLDDK | 97.59% (486/498) | Non-Toxin | Consensus (comb.lib./smm/nn) | 0.58 |
| 25 | N Protein | DRB1*11:01 | GYYRRATRRIRGGDG | 99.40% (495/498) | Non-Toxin | Consensus (smm/nn/sturniolo) | 0.42 |
| 26 | N Protein | DRB1*11:01 | IGYYRRATRRIRGGD | 99.40% (495/498) | Non-Toxin | Consensus (smm/nn/sturniolo) | 0.42 |
| 27 | N Protein | DQA1*01:02/DQB1*06:02 | NNAAIVLQLPQGTTL | 97.59% (486/498) | Non-Toxin | Consensus (comb.lib./smm/nn) | 0.42 |
| 28 | N Protein | DQA1*01:02/DQB1*06:02 | NPANNAAIVLQLPQG | 97.59% (486/498) | Non-Toxin | Consensus (comb.lib./smm/nn) | 0.03 |
| 29 | N Protein | DQA1*01:02/DQB1*06:02 | PANNAAIVLQLPQGT | 97.59% (486/498) | Non-Toxin | Consensus (comb.lib./smm/nn) | 0.04 |
| 30 | N Protein | DRB1*09:01 | QIAQFAPSASAFFGM | 97.39% (485/498) | Non-Toxin | Consensus (comb.lib./smm/nn) | 0.01 |
| 31 | N Protein | DRB1*11:01 | QIGYYRRATRRIRGG | 99.40% (495/498) | Non-Toxin | Consensus (smm/nn/sturniolo) | 0.39 |
| 32 | N Protein | DRB5*01:01 | QIGYYRRATRRIRGG | 99.40% (495/498) | Non-Toxin | Consensus (smm/nn/sturniolo) | 0.58 |
| 33 | N Protein | DQA1*01:02/DQB1*06:02 | RNPANNAAIVLQLPQ | 97.59% (486/498) | Non-Toxin | Consensus (comb.lib./smm/nn) | 0.04 |
| 34 | N Protein | DRB1*07:01 | SGTWLTYTGAIKLDD | 97.39% (485/498) | Non-Toxin | Consensus (comb.lib./smm/nn) | 0.58 |
| 35 | N Protein | DRB1*07:01 | TPSGTWLTYTGAIKL | 97.39% (485/498) | Non-Toxin | Consensus (comb.lib./smm/nn) | 0.58 |
| 36 | N Protein | DQA1*01:02/DQB1*06:02 | TRNPANNAAIVLQLP | 97.39% (485/498) | Non-Toxin | Consensus (comb.lib./smm/nn) | 0.04 |
| 37 | N Protein | DRB1*07:01 | TWLTYTGAIKLDDKD | 97.59% (486/498) | Non-Toxin | Consensus (comb.lib./smm/nn) | 0.58 |
| 38 | ORF1ab | DPA1*01:03/DPB1*02:01 | WFLAYILFTRFFYVL | 100% (456/456) | Non-Toxin | Consensus (comb.lib./smm/nn) | 0.01 |
| 39 | ORF1ab | DPA1*01:03/DPB1*02:01 | FLAYILFTRFFYVLG | 100% (456/456) | Non-Toxin | Consensus (comb.lib./smm/nn) | 0.01 |
| 40 | ORF1ab | DPA1*01:03/DPB1*02:01 | LAYILFTRFFYVLGL | 100% (456/456) | Non-Toxin | Consensus (comb.lib./smm/nn) | 0.01 |
| 41 | ORF1ab | DPA1*01:03/DPB1*02:01 | AYILFTRFFYVLGLA | 100% (456/456) | Non-Toxin | Consensus (comb.lib./smm/nn) | 0.01 |
| 42 | ORF1ab | DRB1*15:01 | AMPNMLRIMASLVLA | 100% (456/456) | Non-Toxin | Consensus (smm/nn/sturniolo) | 0.01 |
| 43 | ORF1ab | DRB1*15:01 | MPNMLRIMASLVLAR | 100% (456/456) | Non-Toxin | Consensus (smm/nn/sturniolo) | 0.01 |
| 44 | ORF1ab | DRB1*15:01 | PNMLRIMASLVLARK | 100% (456/456) | Non-Toxin | Consensus (smm/nn/sturniolo) | 0.01 |
| 45 | ORF1ab | DRB1*15:01 | NMLRIMASLVLARKH | 100% (456/456) | Non-Toxin | Consensus (smm/nn/sturniolo) | 0.01 |
| 46 | ORF1ab | DRB3*02:02 | FAWWTAFVTNVNASS | 98.46% (449/456) | Non-Toxin | NetMHCIIpan | 0.01 |
| 47 | ORF1ab | DPA1*01:03/DPB1*02:01 | AAIMQLFFSYFAVHF | 100% (456/456) | Non-Toxin | Consensus (comb.lib./smm/nn) | 0.05 |
| 48 | ORF1ab | DQA1*01:02/DQB1*06:02 | AFASEAARVVRSIFS | 99.56% (454/456) | Non-Toxin | Consensus (comb.lib./smm/nn) | 0.08 |
| 49 | ORF1ab | DPA1*01:03/DPB1*02:01 | AIMQLFFSYFAVHFI | 100% (456/456) | Non-Toxin | Consensus (comb.lib./smm/nn) | 0.05 |
| 50 | ORF1ab | DRB3*02:02 | ANYIFWRNTNPIQLS | 98.90% (451/456) | Non-Toxin | NetMHCIIpan | 0.05 |
| 51 | ORF1ab | DQA1*05:01/DQB1*03:01 | ASIVAGGIVAIVVTC | 100% (456/456) | Non-Toxin | Consensus (comb.lib./smm/nn) | 0.03 |
| 52 | ORF1ab | DRB3*02:02 | AWWTAFVTNVNASSS | 98.46% (449/456) | Non-Toxin | NetMHCIIpan | 0.01 |
| 53 | ORF1ab | DQA1*05:01/DQB1*03:01 | DISASIVAGGIVAIV | 100% (456/456) | Non-Toxin | Consensus (comb.lib./smm/nn) | 0.03 |
| 54 | ORF1ab | DPA1*03:01/DPB1*04:02 | EETKFLTENLLLYID | 99.78% (455/456) | Non-Toxin | Consensus (comb.lib./smm/nn) | 0.03 |
| 55 | ORF1ab | DQA1*05:01/DQB1*02:01 | EIDFLELAMDEFIER | 99.12% (452/456) | Non-Toxin | Consensus (comb.lib./smm/nn) | 0.04 |
| 56 | ORF1ab | DPA1*03:01/DPB1*04:02 | ETKFLTENLLLYIDI | 99.78% (455/456) | Non-Toxin | Consensus (comb.lib./smm/nn) | 0.03 |
| 57 | ORF1ab | DQA1*01:01/DQB1*05:01 | FISNSWLMWLIINLV | 100% (456/456) | Non-Toxin | Consensus (comb.lib./smm/nn) | 0.05 |
| 58 | ORF1ab | DRB1*07:01 | FSAVGNICYTPSKLI | 99.56% (454/456) | Non-Toxin | Consensus (comb.lib./smm/nn) | 0.06 |
| 59 | ORF1ab | DRB1*07:01 | FTPLVPFWITIAYII | 100% (456/456) | Non-Toxin | Consensus (comb.lib./smm/nn) | 0.03 |
| 60 | ORF1ab | DQA1*01:01/DQB1*05:01 | HFISNSWLMWLIINL | 100% (456/456) | Non-Toxin | Consensus (comb.lib./smm/nn) | 0.05 |
| 61 | ORF1ab | DQA1*05:01/DQB1*02:01 | IDFLELAMDEFIERY | 99.34% (453/456) | Non-Toxin | Consensus (comb.lib./smm/nn) | 0.05 |
| 62 | ORF1ab | DQA1*05:01/DQB1*03:01 | ISASIVAGGIVAIVV | 100% (456/456) | Non-Toxin | Consensus (comb.lib./smm/nn) | 0.03 |
| 63 | ORF1ab | DQA1*01:01/DQB1*05:01 | ISNSWLMWLIINLVQ | 100% (456/456) | Non-Toxin | Consensus (comb.lib./smm/nn) | 0.07 |
| 64 | ORF1ab | DPA1*03:01/DPB1*04:02 | KLINIIIWFLLLSVC | 97.80% (446/456) | Non-Toxin | Consensus (comb.lib./smm/nn) | 0.09 |
| 65 | ORF1ab | DPA1*02:01/DPB1*05:01 | KVTLVFLFVAAIFYL | 99.78% (455/456) | Non-Toxin | Consensus (comb.lib./smm/nn) | 0.04 |

| | | | | | | | |
|---|---|---|---|---|---|---|---|
| 66 | ORF1ab | DPA1*01:03/DPB1*02:01 | LAAIMQLFFSYFAVH | 100% (456/456) | Non-Toxin | Consensus (comb.lib./smm/nn) | 0.07 |
| 67 | ORF1ab | DPA1*03:01/DPB1*04:02 | LEETKFLTENLLLYI | 99.78% (455/456) | Non-Toxin | Consensus (comb.lib./smm/nn) | 0.03 |
| 68 | ORF1ab | DPA1*03:01/DPB1*04:02 | LINIIIWFLLLSVCL | 97.80% (446/456) | Non-Toxin | Consensus (comb.lib./smm/nn) | 0.09 |
| 69 | ORF1ab | DRB1*07:01 | LVPFWITIAYIICIS | 100% (456/456) | Non-Toxin | Consensus (comb.lib./smm/nn) | 0.03 |
| 70 | ORF1ab | DQA1*05:01/DQB1*02:01 | MEIDFLELAMDEFIE | 99.12% (452/456) | Non-Toxin | Consensus (comb.lib./smm/nn) | 0.04 |
| 71 | ORF1ab | DPA1*02:01/DPB1*14:01 | MNLKYAISAKNRART | 99.78% (455/456) | Non-Toxin | NetMHCIIpan | 0.08 |
| 72 | ORF1ab | DPA1*01:03/DPB1*02:01 | MQLFFSYFAVHFISN | 100% (456/456) | Non-Toxin | Consensus (comb.lib./smm/nn) | 0.05 |
| 73 | ORF1ab | DPA1*01:03/DPB1*02:01 | MYIFFASFYYVWKSY | 99.12% (452/456) | Non-Toxin | Consensus (comb.lib./smm/nn) | 0.04 |
| 74 | ORF1ab | DRB1*11:01 | NEFYAYLRKHFSMMI | 100% (456/456) | Non-Toxin | Consensus (smm/nn/sturniolo) | 0.02 |
| 75 | ORF1ab | DRB3*02:02 | NYIFWRNTNPIQLSS | 98.90% (451/456) | Non-Toxin | NetMHCIIpan | 0.04 |
| 76 | ORF1ab | DRB1*07:01 | PLVPFWITIAYIICI | 100% (456/456) | Non-Toxin | Consensus (comb.lib./smm/nn) | 0.02 |
| 77 | ORF1ab | DQA1*05:01/DQB1*02:01 | QMEIDFLELAMDEFI | 99.12% (452/456) | Non-Toxin | Consensus (comb.lib./smm/nn) | 0.03 |
| 78 | ORF1ab | DPA1*02:01/DPB1*14:01 | QMNLKYAISAKNRAR | 99.78% (455/456) | Non-Toxin | NetMHCIIpan | 0.07 |
| 79 | ORF1ab | DRB1*01:01 | QQESPFVMMSAPPAQ | 99.78% (455/456) | Non-Toxin | Consensus (comb.lib./smm/nn) | 0.1 |
| 80 | ORF1ab | DQA1*01:01/DQB1*05:01 | QSTQWSLFFFLYENA | 92.10% (420/456) | Non-Toxin | Consensus (comb.lib./smm/nn) | 0.1 |
| 81 | ORF1ab | DPA1*01:03/DPB1*02:01 | QWSLFFFLYENAFLP | 92.10% (420/456) | Non-Toxin | Consensus (comb.lib./smm/nn) | 0.04 |
| 82 | ORF1ab | DPA1*01:03/DPB1*02:01 | RMYIFFASFYYVWKS | 99.12% (452/456) | Non-Toxin | Consensus (comb.lib./smm/nn) | 0.04 |
| 83 | ORF1ab | DQA1*05:01/DQB1*03:01 | SASIVAGGIVAIVVT | 100% (456/456) | Non-Toxin | Consensus (comb.lib./smm/nn) | 0.03 |
| 84 | ORF1ab | DRB1*07:01 | SAVGNICYTPSKLIE | 99.56% (454/456) | Non-Toxin | Consensus (comb.lib./smm/nn) | 0.09 |
| 85 | ORF1ab | DQA1*05:01/DQB1*03:01 | SIVAGGIVAIVVTCL | 100% (456/456) | Non-Toxin | Consensus (comb.lib./smm/nn) | 0.03 |
| 86 | ORF1ab | DPA1*03:01/DPB1*04:02 | SKLINIIIWFLLLSV | 97.80% (446/456) | Non-Toxin | Consensus (comb.lib./smm/nn) | 0.09 |
| 87 | ORF1ab | DQA1*01:01/DQB1*05:01 | SNSWLMWLIINLVQM | 100% (456/456) | Non-Toxin | Consensus (comb.lib./smm/nn) | 0.07 |
| 88 | ORF1ab | DQA1*05:01/DQB1*02:01 | SQMEIDFLELAMDEF | 99.12% (452/456) | Non-Toxin | Consensus (comb.lib./smm/nn) | 0.05 |
| 89 | ORF1ab | DPA1*01:03/DPB1*02:01 | STQWSLFFFLYENAF | 92.10% (420/456) | Non-Toxin | Consensus (comb.lib./smm/nn) | 0.05 |
| 90 | ORF1ab | DQA1*01:01/DQB1*05:01 | STQWSLFFFLYENAF | 92.10% (420/456) | Non-Toxin | Consensus (comb.lib./smm/nn) | 0.1 |
| 91 | ORF1ab | DPA1*03:01/DPB1*04:02 | TKFLTENLLLYIDIN | 99.78% (455/456) | Non-Toxin | Consensus (comb.lib./smm/nn) | 0.05 |
| 92 | ORF1ab | DPA1*03:01/DPB1*04:02 | TLEETKFLTENLLLY | 99.78% (455/456) | Non-Toxin | Consensus (comb.lib./smm/nn) | 0.05 |
| 93 | ORF1ab | DRB1*07:01 | TPLVPFWITIAYIIC | 100% (456/456) | Non-Toxin | Consensus (comb.lib./smm/nn) | 0.03 |
| 94 | ORF1ab | DPA1*01:03/DPB1*02:01 | TQWSLFFFLYENAFL | 92.10% (420/456) | Non-Toxin | Consensus (comb.lib./smm/nn) | 0.04 |
| 95 | ORF1ab | DQA1*01:01/DQB1*05:01 | TQWSLFFFLYENAFL | 92.10% (420/456) | Non-Toxin | Consensus (comb.lib./smm/nn) | 0.09 |
| 96 | ORF1ab | DRB1*11:01 | VNEFYAYLRKHFSMM | 100% (456/456) | Non-Toxin | Consensus (smm/nn/sturniolo) | 0.05 |
| 97 | ORF1ab | DRB1*01:01 | VQQESPFVMMSAPPA | 99.78% (455/456) | Non-Toxin | Consensus (comb.lib./smm/nn) | 0.1 |
| 98 | ORF1ab | DQA1*01:02/DQB1*06:02 | YAFASEAARVVRSIF | 99.78% (455/456) | Non-Toxin | Consensus (comb.lib./smm/nn) | 0.08 |
| 99 | ORF1ab | DPA1*01:03/DPB1*02:01 | YIFFASFYYVWKSYV | 99.56% (454/456) | Non-Toxin | Consensus (comb.lib./smm/nn) | 0.05 |
| 100 | ORF1ab | DRB3*02:02 | YIFWRNTNPIQLSSY | 98.90% (451/456) | Non-Toxin | NetMHCIIpan | 0.05 |
| 101 | ORF1ab | DPA1*01:03/DPB1*02:01 | YILFTRFFYVLGLAA | 100% (456/456) | Non-Toxin | Consensus (comb.lib./smm/nn) | 0.08 |
| 102 | ORF3a | DPA1*01/DPB1*04:01 | APFLYLYALVYFLQS | 96.88% (466/481) | Non-Toxin | Consensus (comb.lib./smm) | 0.12 |
| 103 | ORF3a | DPA1*02:01/DPB1*14:01 | DFVRATATIPIQASL | 99.37% (478/481) | Non-Toxin | NetMHCIIpan | 0.12 |
| 104 | ORF3a | DPA1*01:03/DPB1*02:01 | DTGVEHVTFFIYNKI | 99.37% (478/481) | Non-Toxin | Consensus (comb.lib./smm/nn) | 1.1 |
| 105 | ORF3a | DPA1*02:01/DPB1*05:01 | DTGVEHVTFFIYNKI | 99.37% (478/481) | Non-Toxin | Consensus (comb.lib./smm/nn) | 1.6 |
| 106 | ORF3a | DRB1*04:05 | FFIYNKIVDEPEEHV | 99.58% (479/481) | Non-Toxin | Consensus (smm/nn/sturniolo) | 0.94 |
| 107 | ORF3a | DPA1*01/DPB1*04:01 | FLYLYALVYFLQSIN | 96.67% (465/481) | Non-Toxin | Consensus (comb.lib./smm) | 0.12 |
| 108 | ORF3a | DPA1*01:03/DPB1*02:01 | GVEHVTFFIYNKIVD | 99.37% (478/481) | Non-Toxin | Consensus (comb.lib./smm/nn) | 1.2 |

| | | | | | | | |
|---|---|---|---|---|---|---|---|
| 109 | ORF3a | DRB1*04:05 | HVTFFIYNKIVDEPE | 99.58% (479/481) | Non-Toxin | Consensus (smm/nn/sturniolo) | 0.84 |
| 110 | ORF3a | DPA1*01/DPB1*04:01 | PFLYLYALVYFLQSI | 96.67% (465/481) | Non-Toxin | Consensus (comb.lib./smm) | 0.12 |
| 111 | ORF3a | DRB1*04:05 | TFFIYNKIVDEPEEH | 99.58% (479/481) | Non-Toxin | Consensus (smm/nn/sturniolo) | 0.81 |
| 112 | ORF3a | DPA1*01:03/DPB1*02:01 | TGVEHVTFFIYNKIV | 99.37% (478/481) | Non-Toxin | Consensus (comb.lib./smm/nn) | 0.93 |
| 113 | ORF3a | DPA1*02:01/DPB1*05:01 | TGVEHVTFFIYNKIV | 99.37% (478/481) | Non-Toxin | Consensus (comb.lib./smm/nn) | 1.4 |
| 114 | ORF3a | DRB1*04:05 | VTFFIYNKIVDEPEE | 99.58% (479/481) | Non-Toxin | Consensus (smm/nn/sturniolo) | 0.81 |
| 115 | ORF3a | DPA1*01/DPB1*04:01 | YLYALVYFLQSINFV | 96.46% (464/481) | Non-Toxin | Consensus (comb.lib./smm) | 0.14 |
| 116 | ORF6 | DRB1*15:01 | EILLIIMRTFKVSIW | 99.58% (479/481) | Non-Toxin | Consensus (smm/nn/sturniolo) | 0.1 |
| 117 | ORF6 | DRB1*15:01 | ILLIIMRTFKVSIWN | 99.38% (478/481) | Non-Toxin | Consensus (smm/nn/sturniolo) | 0.1 |
| 118 | ORF6 | DQA1*01:01/DQB1*05:01 | VSIWNLDYIINLIIK | 99.38% (478/481) | Non-Toxin | Consensus (comb.lib./smm/nn) | 0.05 |
| 119 | ORF7a | DRB1*01:01 | KIILFLALITLATCE | 99.58% (478/480) | Non-Toxin | Consensus (comb.lib./smm/nn) | 0.16 |
| 120 | ORF7b | DRB4*01:01 | AFLLFLVLIMLIIFW | 99.58% (235/236) | Non-Toxin | Consensus (comb.lib./smm/nn) | 0.14 |
| 121 | ORF7b | DPA1*03:01/DPB1*04:02 | DFYLCFLAFLLFLVL | 98.31% (232/236) | Non-Toxin | Consensus (comb.lib./smm/nn) | 0.03 |
| 122 | ORF7b | DPA1*01:03/DPB1*02:01 | DFYLCFLAFLLFLVL | 98.31% (232/236) | Non-Toxin | Consensus (comb.lib./smm/nn) | 0.08 |
| 123 | ORF7b | DPA1*01/DPB1*04:01 | DFYLCFLAFLLFLVL | 98.31% (232/236) | Non-Toxin | Consensus (comb.lib./smm) | 0.09 |
| 124 | ORF7b | DPA1*03:01/DPB1*04:02 | FLAFLLFLVLIMLII | 97.88% (231/236) | Non-Toxin | Consensus (comb.lib./smm/nn) | 0.06 |
| 125 | ORF7b | DRB4*01:01 | FLAFLLFLVLIMLII | 97.88% (231/236) | Non-Toxin | Consensus (comb.lib./smm/nn) | 0.14 |
| 126 | ORF7b | DRB4*01:01 | FLLFLVLIMLIIFWF | 99.58% (235/236) | Non-Toxin | Consensus (comb.lib./smm/nn) | 0.19 |
| 127 | ORF7b | DPA1*03:01/DPB1*04:02 | FYLCFLAFLLFLVLI | 97.88% (231/236) | Non-Toxin | Consensus (comb.lib./smm/nn) | 0.03 |
| 128 | ORF7b | DPA1*01:03/DPB1*02:01 | FYLCFLAFLLFLVLI | 97.88% (231/236) | Non-Toxin | Consensus (comb.lib./smm/nn) | 0.06 |
| 129 | ORF7b | DPA1*01/DPB1*04:01 | FYLCFLAFLLFLVLI | 97.88% (231/236) | Non-Toxin | Consensus (comb.lib./smm) | 0.09 |
| 130 | ORF7b | DPA1*01:03/DPB1*02:01 | IDFYLCFLAFLLFLV | 98.31% (232/236) | Non-Toxin | Consensus (comb.lib./smm/nn) | 0.06 |
| 131 | ORF7b | DPA1*03:01/DPB1*04:02 | IDFYLCFLAFLLFLV | 98.31% (232/236) | Non-Toxin | Consensus (comb.lib./smm/nn) | 0.16 |
| 132 | ORF7b | DPA1*03:01/DPB1*04:02 | LAFLLFLVLIMLIIF | 99.58% (235/236) | Non-Toxin | Consensus (comb.lib./smm/nn) | 0.08 |
| 133 | ORF7b | DRB4*01:01 | LAFLLFLVLIMLIIF | 99.58% (235/236) | Non-Toxin | Consensus (comb.lib./smm/nn) | 0.12 |
| 134 | ORF7b | DPA1*01:03/DPB1*02:01 | LIDFYLCFLAFLLFL | 98.31% (232/236) | Non-Toxin | Consensus (comb.lib./smm/nn) | 0.05 |
| 135 | ORF7b | DPA1*01:03/DPB1*02:01 | LSLIDFYLCFLAFLL | 98.31% (232/236) | Non-Toxin | Consensus (comb.lib./smm/nn) | 0.15 |
| 136 | ORF7b | DQA1*01:01/DQB1*05:01 | LVLIMLIIFWFSLEL | 99.58% (235/236) | Non-Toxin | Consensus (comb.lib./smm/nn) | 0.14 |
| 137 | ORF7b | DPA1*01:03/DPB1*02:01 | SLIDFYLCFLAFLLF | 98.31% (232/236) | Non-Toxin | Consensus (comb.lib./smm/nn) | 0.07 |
| 138 | ORF8 | DQA1*01:01/DQB1*05:01 | LVVRCSFYEDFLEYH | 99.58% (478/480) | Non-Toxin | Consensus (comb.lib./smm/nn) | 0.45 |
| 139 | ORF8 | DRB3*01:01 | QHQPYVVDDPCPIHF | 99.17% (476/480) | Non-Toxin | Consensus (comb.lib./smm/nn) | 0.08 |
| 140 | ORF8 | DRB3*01:01 | TQHQPYVVDDPCPIH | 99.17% (476/480) | Non-Toxin | Consensus (comb.lib./smm/nn) | 0.08 |
| 141 | ORF10 | DPA1*01/DPB1*04:01 | FAFPFTIYSLLLCRM | 99.79% (478/479) | Non-Toxin | Consensus (comb.lib./smm) | 0.56 |
| 142 | ORF10 | DPA1*02:01/DPB1*01:01 | FAFPFTIYSLLLCRM | 99.79% (478/479) | Non-Toxin | Consensus (comb.lib./smm/nn) | 1.1 |
| 143 | ORF10 | DPA1*03:01/DPB1*04:02 | FAFPFTIYSLLLCRM | 99.79% (478/479) | Non-Toxin | Consensus (comb.lib./smm/nn) | 1.1 |
| 144 | ORF10 | DPA1*01:03/DPB1*02:01 | INVFAFPFTIYSLLL | 99.37% (476/479) | Non-Toxin | Consensus (comb.lib./smm/nn) | 0.29 |
| 145 | ORF10 | DPA1*01/DPB1*04:01 | INVFAFPFTIYSLLL | 99.37% (476/479) | Non-Toxin | Consensus (comb.lib./smm) | 0.46 |
| 146 | ORF10 | DPA1*02:01/DPB1*01:01 | INVFAFPFTIYSLLL | 99.37% (476/479) | Non-Toxin | Consensus (comb.lib./smm/nn) | 0.72 |
| 147 | ORF10 | DPA1*01:03/DPB1*02:01 | NVFAFPFTIYSLLLC | 99.58% (477/479) | Non-Toxin | Consensus (comb.lib./smm/nn) | 0.4 |
| 148 | ORF10 | DPA1*01/DPB1*04:01 | NVFAFPFTIYSLLLC | 99.58% (477/479) | Non-Toxin | Consensus (comb.lib./smm) | 0.56 |
| 149 | ORF10 | DPA1*02:01/DPB1*01:01 | NVFAFPFTIYSLLLC | 99.58% (477/479) | Non-Toxin | Consensus (comb.lib./smm/nn) | 0.69 |
| 150 | ORF10 | DPA1*01/DPB1*04:01 | VFAFPFTIYSLLLCR | 99.58% (477/479) | Non-Toxin | Consensus (comb.lib./smm) | 0.56 |
| 151 | ORF10 | DPA1*02:01/DPB1*01:01 | VFAFPFTIYSLLLCR | 99.58% (477/479) | Non-Toxin | Consensus (comb.lib./smm/nn) | 0.66 |

| | | | | | | | |
|---|---|---|---|---|---|---|---|
| 152 | ORF10 | DPA1*01:03/DPB1*02:01 | VFAFPFTIYSLLLCR | 99.58% (477/479) | Non-Toxin | Consensus (comb.lib./smm/nn) | 0.71 |
| 153 | ORF10 | DPA1*03:01/DPB1*04:02 | VFAFPFTIYSLLLCR | 99.58% (477/479) | Non-Toxin | Consensus (comb.lib./smm/nn) | 1.1 |
| 154 | S protein | DRB3*01:01 | ADSFVIRGDEVRQIA | 99.79% (471/472) | Non-Toxin | Consensus (comb.lib./smm/nn) | 0.49 |
| 155 | S protein | DRB1*07:01 | AIPTNFTISVTTEIL | 100.00% (472/472) | Non-Toxin | Consensus (comb.lib./smm/nn) | 0.4 |
| 156 | S protein | DRB3*01:01 | DSFVIRGDEVRQIAP | 99.79% (471/472) | Non-Toxin | Consensus (comb.lib./smm/nn) | 0.51 |
| 157 | S protein | DRB3*02:02 | EGVFVSNGTHWFVTQ | 99.36% (469/472) | Non-Toxin | NetMHCIIpan | 0.21 |
| 158 | S protein | DRB5*01:01 | FVFKNIDGYFKIYSK | 98.09% (463/472) | Non-Toxin | Consensus (smm/nn/sturniolo) | 0.17 |
| 159 | S protein | DRB1*11:01 | GNYNYLYRLFRKSNL | 98.52% (465/472) | Non-Toxin | Consensus (smm/nn/sturniolo) | 0.22 |
| 160 | S protein | DRB1*07:01 | IAIPTNFTISVTTEI | 100.00% (472/472) | Non-Toxin | Consensus (comb.lib./smm/nn) | 0.47 |
| 161 | S protein | DRB5*01:01 | INITRFQTLLALHRS | 99.36% (469/472) | Non-Toxin | Consensus (smm/nn/sturniolo) | 0.32 |
| 162 | S protein | DRB5*01:01 | ITRFQTLLALHRSYL | 99.36% (469/472) | Non-Toxin | Consensus (smm/nn/sturniolo) | 0.26 |
| 163 | S protein | DPA1*02:01/DPB1*14:01 | ITRFQTLLALHRSYL | 99.36% (469/472) | Non-Toxin | NetMHCIIpan | 0.43 |
| 164 | S protein | DRB1*13:02 | LIVNNATNVVIKVCE | 99.36% (469/472) | Non-Toxin | Consensus (smm/nn/sturniolo) | 0.03 |
| 165 | S protein | DRB1*01:01 | LSFELLHAPATVCGP | 98.09% (463/472) | Non-Toxin | Consensus (comb.lib./smm/nn) | 0.03 |
| 166 | S protein | DRB5*01:01 | NITRFQTLLALHRSY | 99.36% (469/472) | Non-Toxin | Consensus (smm/nn/sturniolo) | 0.32 |
| 167 | S protein | DPA1*02:01/DPB1*14:01 | NITRFQTLLALHRSY | 99.36% (469/472) | Non-Toxin | NetMHCIIpan | 0.45 |
| 168 | S protein | DRB1*07:01 | PTNFTISVTTEILPV | 100.00% (472/472) | Non-Toxin | Consensus (comb.lib./smm/nn) | 0.51 |
| 169 | S protein | DPA1*02:01/DPB1*14:01 | QLIRAAEIRASANLA | 99.36% (469/472) | Non-Toxin | NetMHCIIpan | 0.31 |
| 170 | S protein | DPA1*02:01/DPB1*14:01 | QQLIRAAEIRASANL | 99.36% (469/472) | Non-Toxin | NetMHCIIpan | 0.2 |
| 171 | S protein | DRB5*01:01 | REFVFKNIDGYFKIY | 97.88% (462/472) | Non-Toxin | Consensus (smm/nn/sturniolo) | 0.17 |
| 172 | S protein | DRB3*02:02 | REGVFVSNGTHWFVT | 99.58% (470/472) | Non-Toxin | NetMHCIIpan | 0.2 |
| 173 | S protein | DRB1*01:01 | SFELLHAPATVCGPK | 98.09% (463/472) | Non-Toxin | Consensus (comb.lib./smm/nn) | 0.09 |
| 174 | S protein | DRB3*01:01 | SFVIRGDEVRQIAPG | 99.79% (471/472) | Non-Toxin | Consensus (comb.lib./smm/nn) | 0.51 |
| 175 | S protein | DRB1*13:02 | SKTQSLLIVNNATNV | 100.00% (472/472) | Non-Toxin | Consensus (smm/nn/sturniolo) | 0.03 |
| 176 | S protein | DRB1*01:01 | VLSFELLHAPATVCG | 98.09% (463/472) | Non-Toxin | Consensus (comb.lib./smm/nn) | 0.03 |
| 177 | S protein | DRB1*01:01 | VVLSFELLHAPATVC | 98.09% (463/472) | Non-Toxin | Consensus (comb.lib./smm/nn) | 0.03 |
| 178 | S protein | DRB1*01:01 | VVVLSFELLHAPATV | 98.09% (463/472) | Non-Toxin | Consensus (comb.lib./smm/nn) | 0.09 |
| 179 | S protein | DRB3*01:01 | YADSFVIRGDEVRQI | 99.79% (471/472) | Non-Toxin | Consensus (comb.lib./smm/nn) | 0.49 |
| 180 | S protein | DQA1*05:01/DQB1*03:01 | YIWLGFIAGLIAIVM | 99.58% (470/472) | Non-Toxin | Consensus (comb.lib./smm/nn) | 0.51 |

**Supplementary table S11. Shortlisted B Cell epitopes (BepiPred Linear B Cell Prediction).** B cell linear epitopes with length of 4 to 20 amino acids, predicted by the “BepiPred Linear B Cell Prediction” IEDB tool, from eleven SARS-CoV-2 proteins, are listed here. *In-silico* analysis has shown all the selected epitopes to be non-toxic (Non-Toxin) as well as they show significant amino acid sequence conservancy.

| SARS-CoV-2 B Cell epitopes (Bepipred Linear Epitope Prediction) | | | | | |
|---|---|---|---|---|---|
| S.No. | Proteins | Peptide | Conservancy | Toxicity | Length |
| 1 | E protein | SEET | 99.59% (480/482) | Non-Toxin | 4 |
| 2 | E protein | YVYSRVKNLNSSRVP | 99.38% (479/482) | Non-Toxin | 15 |
| 3 | M protein | PLLESE | 99.16% (473/477) | Non-Toxin | 6 |
| 4 | M protein | KLGASQRVAGDS | 98.74% (471/477) | Non-Toxin | 12 |
| 5 | M protein | NGTITVEELKKLLEQW | 98.53% (470/477) | Non-Toxin | 16 |
| 6 | M protein | YRIGNYKLNTDHSSSSDNIA | 98.53% (470/477) | Non-Toxin | 20 |
| 7 | N protein | DPNFKD | 97.19% (484/498) | Non-Toxin | 6 |
| 8 | N protein | AGLPYGANK | 99.40% (495/498) | Non-Toxin | 9 |
| 9 | N protein | NGPQNQRNAPRI | 96.18% (479/498) | Non-Toxin | 12 |
| 10 | N protein | SKQLQQSMSSADS | 97.59% (486/498) | Non-Toxin | 13 |
| 11 | ORF1ab | IQYG | 99.78% (455/456) | Non-Toxin | 4 |
| 12 | ORF1ab | YLKH | 100% (456/456) | Non-Toxin | 4 |
| 13 | ORF1ab | RKVP | 100% (456/456) | Non-Toxin | 4 |
| 14 | ORF1ab | YQCG | 100% (456/456) | Non-Toxin | 4 |
| 15 | ORF1ab | ELSR | 100% (456/456) | Non-Toxin | 4 |
| 16 | ORF1ab | TFCA | 99.78% (455/456) | Non-Toxin | 4 |
| 17 | ORF1ab | GKIV | 100% (456/456) | Non-Toxin | 4 |
| 18 | ORF1ab | EGNC | 100% (456/456) | Non-Toxin | 4 |
| 19 | ORF1ab | LPYP | 100% (456/456) | Non-Toxin | 4 |
| 20 | ORF1ab | SLRC | 100% (456/456) | Non-Toxin | 4 |
| 21 | ORF1ab | KVQI | 100% (456/456) | Non-Toxin | 4 |
| 22 | ORF1ab | HDIG | 100% (456/456) | Non-Toxin | 4 |
| 23 | ORF1ab | WNAD | 98.90% (451/456) | Non-Toxin | 4 |
| 24 | ORF1ab | EIPVA | 99.56% (454/456) | Non-Toxin | 5 |
| 25 | ORF1ab | KGEDI | 99.78% (455/456) | Non-Toxin | 5 |
| 26 | ORF1ab | NLYDK | 100% (456/456) | Non-Toxin | 5 |
| 27 | ORF1ab | KFNPP | 98.68% (450/456) | Non-Toxin | 5 |
| 28 | ORF1ab | GQQQT | 98.46% (449/456) | Non-Toxin | 5 |
| 29 | ORF1ab | KFADD | 99.34% (453/456) | Non-Toxin | 5 |
| 30 | ORF1ab | AAVNS | 100% (456/456) | Non-Toxin | 5 |
| 31 | ORF1ab | SKCEE | 99.12% (452/456) | Non-Toxin | 5 |
| 32 | ORF1ab | IEYTD | 100% (456/456) | Non-Toxin | 5 |
| 33 | ORF1ab | IQPIG | 100% (456/456) | Non-Toxin | 5 |
| 34 | ORF1ab | NSGSD | 100% (456/456) | Non-Toxin | 5 |
| 35 | ORF1ab | EFTPF | 100% (456/456) | Non-Toxin | 5 |

| | | | | | |
|---|---|---|---|---|---|
| 36 | ORF1ab | LLAKD | 100% (456/456) | Non-Toxin | 5 |
| 37 | ORF1ab | EEMLD | 100% (456/456) | Non-Toxin | 5 |
| 38 | ORF1ab | RSEDK | 100% (456/456) | Non-Toxin | 5 |
| 39 | ORF1ab | TGHML | 100% (456/456) | Non-Toxin | 5 |
| 40 | ORF1ab | AHSCN | 99.78% (455/456) | Non-Toxin | 5 |
| 41 | ORF1ab | QTGSS | 99.56% (454/456) | Non-Toxin | 5 |
| 42 | ORF1ab | LPVLQV | 98.90% (451/456) | Non-Toxin | 6 |
| 43 | ORF1ab | FGDSVE | 99.34% (453/456) | Non-Toxin | 6 |
| 44 | ORF1ab | ARTAPH | 99.78% (455/456) | Non-Toxin | 6 |
| 45 | ORF1ab | AGKASC | 100% (456/456) | Non-Toxin | 6 |
| 46 | ORF1ab | DFIDTK | 99.78% (455/456) | Non-Toxin | 6 |
| 47 | ORF1ab | QYELKH | 99.78% (455/456) | Non-Toxin | 6 |
| 48 | ORF1ab | FPDLNG | 99.12% (452/456) | Non-Toxin | 6 |
| 49 | ORF1ab | SLDTYP | 100% (456/456) | Non-Toxin | 6 |
| 50 | ORF1ab | SSFKWD | 99.34% (453/456) | Non-Toxin | 6 |
| 51 | ORF1ab | KRNRAT | 99.78% (455/456) | Non-Toxin | 6 |
| 52 | ORF1ab | KGFCKL | 100% (456/456) | Non-Toxin | 6 |
| 53 | ORF1ab | AQVAKS | 100% (456/456) | Non-Toxin | 6 |
| 54 | ORF1ab | VESSSK | 99.34% (453/456) | Non-Toxin | 6 |
| 55 | ORF1ab | LYDKLQ | 100% (456/456) | Non-Toxin | 6 |
| 56 | ORF1ab | ENKTTL | 100% (456/456) | Non-Toxin | 6 |
| 57 | ORF1ab | DKGVAP | 100% (456/456) | Non-Toxin | 6 |
| 58 | ORF1ab | ATNGPLK | 100% (456/456) | Non-Toxin | 7 |
| 59 | ORF1ab | NKVENMT | 99.78% (455/456) | Non-Toxin | 7 |
| 60 | ORF1ab | TRQVVNV | 100% (456/456) | Non-Toxin | 7 |
| 61 | ORF1ab | EVGFVVP | 100% (456/456) | Non-Toxin | 7 |
| 62 | ORF1ab | DASGKPV | 100% (456/456) | Non-Toxin | 7 |
| 63 | ORF1ab | DYYRSLP | 99.34% (453/456) | Non-Toxin | 7 |
| 64 | ORF1ab | GVSFSTF | 100% (456/456) | Non-Toxin | 7 |
| 65 | ORF1ab | MAFPSGK | 99.56% (454/456) | Non-Toxin | 7 |
| 66 | ORF1ab | SQGLLPP | 99.56% (454/456) | Non-Toxin | 7 |
| 67 | ORF1ab | VQSKMSD | 99.78% (455/456) | Non-Toxin | 7 |
| 68 | ORF1ab | AVDINKL | 99.12% (452/456) | Non-Toxin | 7 |
| 69 | ORF1ab | DRDAAMQ | 99.12% (452/456) | Non-Toxin | 7 |
| 70 | ORF1ab | FPKSDGT | 100% (456/456) | Non-Toxin | 7 |
| 71 | ORF1ab | HPNPKGF | 100% (456/456) | Non-Toxin | 7 |
| 72 | ORF1ab | GNKIADK | 99.56% (454/456) | Non-Toxin | 7 |
| 73 | ORF1ab | YTPHTVL | 99.78% (455/456) | Non-Toxin | 7 |
| 74 | ORF1ab | FEKGDYG | 99.78% (455/456) | Non-Toxin | 7 |
| 75 | ORF1ab | VDSSQGS | 100% (456/456) | Non-Toxin | 7 |
| 76 | ORF1ab | MYKGLPW | 100% (456/456) | Non-Toxin | 7 |
| 77 | ORF1ab | IIGDELK | 99.78% (455/456) | Non-Toxin | 7 |
| 78 | ORF1ab | QAWQPGV | 99.78% (455/456) | Non-Toxin | 7 |

| | | | | | |
|---|---|---|---|---|---|
| 79 | ORF1ab | KPREQID | 98.46% (449/456) | Non-Toxin | 7 |
| 80 | ORF1ab | VPGFNEKT | 99.12% (452/456) | Non-Toxin | 8 |
| 81 | ORF1ab | RELNGGAY | 100% (456/456) | Non-Toxin | 8 |
| 82 | ORF1ab | FTLKGGAP | 99.78% (455/456) | Non-Toxin | 8 |
| 83 | ORF1ab | KTVGELGD | 98.68% (450/456) | Non-Toxin | 8 |
| 84 | ORF1ab | NVPMEKLK | 99.34% (453/456) | Non-Toxin | 8 |
| 85 | ORF1ab | GNATEVPA | 100% (456/456) | Non-Toxin | 8 |
| 86 | ORF1ab | SGGQPITN | 100% (456/456) | Non-Toxin | 8 |
| 87 | ORF1ab | EIIKSQDL | 100% (456/456) | Non-Toxin | 8 |
| 88 | ORF1ab | YTVELGTEV | 100% (456/456) | Non-Toxin | 9 |
| 89 | ORF1ab | IPTKKAGGT | 100% (456/456) | Non-Toxin | 9 |
| 90 | ORF1ab | GLNGYTVEE | 98.68% (450/456) | Non-Toxin | 9 |
| 91 | ORF1ab | YFYTSKTTV | 100% (456/456) | Non-Toxin | 9 |
| 92 | ORF1ab | MSMTYGQQF | 99.78% (455/456) | Non-Toxin | 9 |
| 93 | ORF1ab | AKNVSLDNV | 99.78% (455/456) | Non-Toxin | 9 |
| 94 | ORF1ab | GGVTRDIAS | 100% (456/456) | Non-Toxin | 9 |
| 95 | ORF1ab | VNLHSSRLS | 100% (456/456) | Non-Toxin | 9 |
| 96 | ORF1ab | RNRDVDTDF | 100% (456/456) | Non-Toxin | 9 |
| 97 | ORF1ab | LTKHPNQEY | 100% (456/456) | Non-Toxin | 9 |
| 98 | ORF1ab | TEETFKLSY | 98.90% (451/456) | Non-Toxin | 9 |
| 99 | ORF1ab | IPGIPKDMT | 100% (456/456) | Non-Toxin | 9 |
| 100 | ORF1ab | FHTPAFDKS | 98.24% (448/456) | Non-Toxin | 9 |
| 101 | ORF1ab | KGHFDGQQG | 100% (456/456) | Non-Toxin | 9 |
| 102 | ORF1ab | GVVQQLPET | 99.78% (455/456) | Non-Toxin | 9 |
| 103 | ORF1ab | IERYKLEGY | 99.34% (453/456) | Non-Toxin | 9 |
| 104 | ORF1ab | VEKGVLPQLE | 99.78% (455/456) | Non-Toxin | 10 |
| 105 | ORF1ab | SEKSYELQTP | 99.56% (454/456) | Non-Toxin | 10 |
| 106 | ORF1ab | CHNSEVGPEH | 99.12% (452/456) | Non-Toxin | 10 |
| 107 | ORF1ab | CVKSREETGL | 99.56% (454/456) | Non-Toxin | 10 |
| 108 | ORF1ab | SYKDWSYSGQ | 99.78% (455/456) | Non-Toxin | 10 |
| 109 | ORF1ab | ALLTKSSEYK | 99.78% (455/456) | Non-Toxin | 10 |
| 110 | ORF1ab | QLTGYKKPAS | 99.56% (454/456) | Non-Toxin | 10 |
| 111 | ORF1ab | ILKPANNSLK | 99.56% (454/456) | Non-Toxin | 10 |
| 112 | ORF1ab | YNYEPLTQDH | 98.24% (448/456) | Non-Toxin | 10 |
| 113 | ORF1ab | EQAVANGDSE | 99.78% (455/456) | Non-Toxin | 10 |
| 114 | ORF1ab | PEANMDQESF | 99.78% (455/456) | Non-Toxin | 10 |
| 115 | ORF1ab | PCGTGTSTDV | 97.58% (445/456) | Non-Toxin | 10 |
| 116 | ORF1ab | FQEKDEDDNL | 98.90% (451/456) | Non-Toxin | 10 |
| 117 | ORF1ab | TFSNYQHEET | 100% (456/456) | Non-Toxin | 10 |
| 118 | ORF1ab | FFKFRIDGDM | 99.56% (454/456) | Non-Toxin | 10 |
| 119 | ORF1ab | LKYAISAKNR | 99.78% (455/456) | Non-Toxin | 10 |
| 120 | ORF1ab | YASQGLVASI | 99.56% (454/456) | Non-Toxin | 10 |
| 121 | ORF1ab | WTETDLTKGP | 99.78% (455/456) | Non-Toxin | 10 |

| | | | | | |
|---|---|---|---|---|---|
| 122 | ORF1ab | QVNGYPNMFI | 100% (456/456) | Non-Toxin | 10 |
| 123 | ORF1ab | YKRDAPAHIS | 100% (456/456) | Non-Toxin | 10 |
| 124 | ORF1ab | RVDGQVDLFR | 98.90% (451/456) | Non-Toxin | 10 |
| 125 | ORF1ab | HTKKWKYPQVN | 99.12% (452/456) | Non-Toxin | 11 |
| 126 | ORF1ab | KRPINPTDQSS | 99.34% (453/456) | Non-Toxin | 11 |
| 127 | ORF1ab | LFSTVFPPTSF | 72.58% (331/456) | Non-Toxin | 11 |
| 128 | ORF1ab | VGKPRPPLNRN | 99.34% (453/456) | Non-Toxin | 11 |
| 129 | ORF1ab | RVLSNLNLPGC | 98.90% (451/456) | Non-Toxin | 11 |
| 130 | ORF1ab | DIAKKPTETIC | 99.78% (455/456) | Non-Toxin | 11 |
| 131 | ORF1ab | YHNESGLKTILR | 99.78% (455/456) | Non-Toxin | 12 |
| 132 | ORF1ab | EGETLPTEVLTE | 99.56% (454/456) | Non-Toxin | 12 |
| 133 | ORF1ab | KHYTPSFKKGAK | 99.56% (454/456) | Non-Toxin | 12 |
| 134 | ORF1ab | RQGFVDSDVETK | 99.78% (455/456) | Non-Toxin | 12 |
| 135 | ORF1ab | SDVLLPLTQYNR | 99.78% (455/456) | Non-Toxin | 12 |
| 136 | ORF1ab | KLQNNELSPVAL | 99.56% (454/456) | Non-Toxin | 12 |
| 137 | ORF1ab | MHAASGNLLLDK | 99.78% (455/456) | Non-Toxin | 12 |
| 138 | ORF1ab | KPGGTSSGDATT | 100% (456/456) | Non-Toxin | 12 |
| 139 | ORF1ab | TNDNTSRYWEPE | 100% (456/456) | Non-Toxin | 12 |
| 140 | ORF1ab | PYVCNAPGCDVT | 100% (456/456) | Non-Toxin | 12 |
| 141 | ORF1ab | GCHATREAVGTN | 98.24% (448/456) | Non-Toxin | 12 |
| 142 | ORF1ab | SVKGLQPSVGPK | 99.78% (455/456) | Non-Toxin | 12 |
| 143 | ORF1ab | LNDFVSDADSTL | 99.78% (455/456) | Non-Toxin | 12 |
| 144 | ORF1ab | YILPSIISNEKQE | 100% (456/456) | Non-Toxin | 13 |
| 145 | ORF1ab | GLNLEEAARYMRS | 100% (456/456) | Non-Toxin | 13 |
| 146 | ORF1ab | FDNLKTLLSLREV | 99.78% (455/456) | Non-Toxin | 13 |
| 147 | ORF1ab | EQFKKGVQIPCTC | 99.56% (454/456) | Non-Toxin | 13 |
| 148 | ORF1ab | EDMLNPNYEDLLI | 99.56% (454/456) | Non-Toxin | 13 |
| 149 | ORF1ab | RNVATLQAENVTG | 100% (456/456) | Non-Toxin | 13 |
| 150 | ORF1ab | PKTKNVTKENDSK | 98.90% (451/456) | Non-Toxin | 13 |
| 151 | ORF1ab | YNKYKYFSGAMDTT | 99.56% (454/456) | Non-Toxin | 14 |
| 152 | ORF1ab | EEAIRHVRAWIGFD | 100% (456/456) | Non-Toxin | 14 |
| 153 | ORF1ab | CGETSWQTGDFVKAT | 99.34% (453/456) | Non-Toxin | 15 |
| 154 | ORF1ab | LQPLEQPTSEAVEAP | 99.56% (454/456) | Non-Toxin | 15 |
| 155 | ORF1ab | VSELLTPLGIDLDEW | 99.78% (455/456) | Non-Toxin | 15 |
| 156 | ORF1ab | SKHTDFSSEIIGYKA | 98.90% (451/456) | Non-Toxin | 15 |
| 157 | ORF1ab | PAPRTLLTKGTLEPE | 99.34% (453/456) | Non-Toxin | 15 |
| 158 | ORF1ab | KRNIKPVPEVKILNN | 98.90% (451/456) | Non-Toxin | 15 |
| 159 | ORF1ab | RFKESPFELEDFIPM | 99.34% (453/456) | Non-Toxin | 15 |
| 160 | ORF1ab | GTENLTKEGATTCGYLP | 99.12% (452/456) | Non-Toxin | 17 |
| 161 | ORF1ab | DFDTWFSQRGGSYTNDK | 99.78% (455/456) | Non-Toxin | 17 |
| 162 | ORF1ab | TANPKTPKYKFVRIQPG | 99.12% (452/456) | Non-Toxin | 17 |
| 163 | ORF1ab | DDYFNKKDWYDFVENPD | 98.90% (451/456) | Non-Toxin | 17 |
| 164 | ORF1ab | LQKEKVNINIVGDFKLNE | 100% (456/456) | Non-Toxin | 18 |

| | | | | | |
|---|---|---|---|---|---|
| 165 | ORF1ab | WDTIANYAKPFLNKVVST | 98.68% (450/456) | Non-Toxin | 18 |
| 166 | ORF1ab | AVTAYNGYLTSSSKTPEEH | 100% (456/456) | Non-Toxin | 19 |
| 167 | ORF1ab | GCSCDQLREPMLQSADAQS | 98.90% (451/456) | Non-Toxin | 19 |
| 168 | ORF1ab | PNNTDFSRVSAKPPPGDQF | 99.78% (455/456) | Non-Toxin | 19 |
| 169 | ORF1ab | RMLLEKCDLQNYGDSATLP | 100% (456/456) | Non-Toxin | 19 |
| 170 | ORF1ab | IVSTIQRKYKGIKIQEGVVD | 99.12% (452/456) | Non-Toxin | 20 |
| 171 | ORF1ab | GTTQTACTDDNALAYYNTTK | 99.78% (455/456) | Non-Toxin | 20 |
| 172 | ORF1ab | IQLSSYSLFDMSKFPLKLRG | 98.46% (449/456) | Non-Toxin | 20 |
| 173 | ORF3a | SKNPLL | 96.25% (463/481) | Non-Toxin | 6 |
| 174 | ORF3a | PYNSVT | 97.92% (471/481) | Non-Toxin | 6 |
| 175 | ORF3a | STQLSTDTGV | 99.58%(479/481) | Non-Toxin | 10 |
| 176 | ORF3a | KIITLKKRWQL | 99.37% (478/481) | Non-Toxin | 11 |
| 177 | ORF3a | QGEIKDATPSDF | 99.37% (478/481) | Non-Toxin | 12 |
| 178 | ORF6 | LTENKYSQLDEEQP | 99.17% (477/481) | Non-Toxin | 14 |
| 179 | ORF7a | LYHYQECVR | 99.79% (479/480) | Non-Toxin | 9 |
| 180 | ORF7b | ELQDHNE | 100.00% (236/236) | Non-Toxin | 7 |
| 181 | ORF8 | QEPKL | 100.00% (480/480) | Non-Toxin | 5 |
| 182 | ORF8 | EDFLEY | 99.79% (479/480) | Non-Toxin | 6 |
| 183 | ORF8 | RVGARKSAP | 99.17% (476/480) | Non-Toxin | 9 |
| 184 | ORF8 | DEAGSKSPIQYIDIGN | 98.96% (475/480) | Non-Toxin | 16 |
| 185 | S protein | LDPL | 99.58% (470/472) | Non-Toxin | 4 |
| 186 | S protein | QTLE | 100.00% (472/472) | Non-Toxin | 4 |
| 187 | S protein | LGKY | 100.00% (472/472) | Non-Toxin | 4 |
| 188 | S protein | TNTSN | 100.00% (472/472) | Non-Toxin | 5 |
| 189 | S protein | EAEVQ | 99.15% (468/472) | Non-Toxin | 5 |
| 190 | S protein | EQDKNTQ | 99.15% (468/472) | Non-Toxin | 7 |
| 191 | S protein | SNKKFLPF | 100.00% (472/472) | Non-Toxin | 8 |
| 192 | S protein | PDPSKPSK | 98.31% (464/472) | Non-Toxin | 8 |
| 193 | S protein | GQSKRVDFC | 100.00% (472/472) | Non-Toxin | 9 |
| 194 | S protein | TPGDSSSGWTA | 99.58% (470/472) | Non-Toxin | 11 |
| 195 | S protein | RVYSTGSNVFQ | 99.79% (471/472) | Non-Toxin | 11 |
| 196 | S protein | VNNSYECDIPI | 100.00% (472/472) | Non-Toxin | 11 |
| 197 | S protein | RNFYEPQIITTD | 99.36% (469/472) | Non-Toxin | 12 |
| 198 | S protein | MDLEGKQGNFKNL | 98.52% (465/472) | Non-Toxin | 13 |
| 199 | S protein | KQIYKTPPIKDFGGF | 98.09% (463/472) | Non-Toxin | 15 |
| 200 | S protein | LADAGFIKQYGDCLG | 99.36% (469/472) | Non-Toxin | 15 |
| 201 | S protein | KHTPINLVRDLPQGFS | 99.58% (470/472) | Non-Toxin | 16 |
| 202 | S protein | YTMSLGAENSVAYSNN | 100.00% (472/472) | Non-Toxin | 16 |
| 203 | S protein | DPFLGVYYHKNNKSWME | 98.09% (463/472) | Non-Toxin | 17 |
| 204 | S protein | NCTEVPVAIHADQLTPT | 100.00% (472/472) | Non-Toxin | 17 |
| 205 | S protein | KSFTVEKGIYQTSNFRVQP | 99.36% (469/472) | Non-Toxin | 19 |
| 206 | S protein | ASYQTQTNSPRRARSVASQ | 100.00% (472/472) | Non-Toxin | 19 |
